# Characterization and orthogonality assessment of two quorum sensing systems for synthetic biology applications

**DOI:** 10.1101/2025.04.03.643276

**Authors:** De Baets Jasmine, De Paepe Brecht, De Mey Marjan

**Affiliations:** Centre for Synthetic Biology, Ghent University, Ghent, 9000, Belgium

**Keywords:** Quorum sensing, genetic circuits, synthetic biology, characterization, orthogonality

## Abstract

Quorum sensing systems have a broad range of applications within the field of synthetic biology. However, a bottleneck is the optimization and tuning of these systems due to the lack of standardization and complete characterization. In this research, two quorum sensing systems, namely the LasI/LasR and the EsaI/EsaR system, were fully characterized in the model host organism *Escherichia coli*. Furthermore, insight was gained in the interplay between the various parts of these systems. To further expand the range of possibilities with these quorum sensing systems, the orthogonality of the two systems was assessed to allow simultaneous use within the same cell without interfering crosstalk. This assessment was performed on three levels: promoter, signal and synthase crosstalk. It was demonstrated that LasR is able to interact with the promoter of the EsaI/EsaR system, albeit to a low extent. Additionally, LasR was able to respond to the autoinducers produced by EsaI. To solve the promoter crosstalk, a nucleotide change was introduced into the binding site of EsaR within the promoter region. Additionally, LasR mutants were created rationally and screened for decreased response to EsaI while retaining functionality. The best performing mutant, LasR(P117S), was further characterized. In conclusion, we have further unlocked the potential of quorum sensing systems for synthetic biology applications by obtaining two functional, characterized and orthogonal quorum sensing systems.

**Highlights:**

– Characterization of two LuxR-type quorum sensing systems
– Assessing the orthogonality of the EsaI/EsaR and LasI/LasR quorum sensing system
– Eliminating the crosstalk between the EsaI/EsaR and LasI/LasR quorum sensing system

## 1. Introduction

Even though bacteria are simple, single-celled organisms, they are able to communicate and interact with neighboring cells by a mechanism called quorum sensing. Quorum sensing allows bacteria to detect the cell density of surrounding bacteria by producing and sensing autoinducer (AI) molecules that are released in the environment. This renders them the ability to form a more complex multicellular population with a common goal, *e.g.*, biofilm formation or virulence [1]. There are three well-investigated types of quorum sensing. The most common and described quorum sensing system (QS system) occurs in gram-negative bacteria by synthesizing *N*-acyl-homoserine lactones (AHLs), small molecules that freely diffuse through the cell membrane [2,3]. The second group of AIs consists of modified oligopeptides produced by gram-positive bacteria. The transport of these molecules requires ATP-binding cassette transporters and their detection occurs through specific two-component systems [4]. Thirdly, autoinducer-2 (AI-2) molecules, derivatives of 4,5-dihydroxy-2,3-pentanedione, are used by both gram-positive and –negative bacteria for intra– and interspecies communication [3]. Quorum sensing has drawn a lot of attention from synthetic biologists, because it opens the possibilities for engineering bacteria on a population level (Figure 1) [5]. Especially the limited number of genes required for introducing an AHL-activated QS system allows easy implementation in different hosts and functions as a starting point of many genetic circuits. These genetic circuits cover a broad range of possible applications, such as a cell density-based toggle switch [6], the autoinducible activation of a production pathway at high cell density [7] and regulating the dynamics of two populations in a bacterial consortium [8,9]. However, the main interest still seems to go toward the autoinducible expression of production pathways to obtain a decoupling of the growth and production stage during a fermentation process [10].

**Figure 1.**
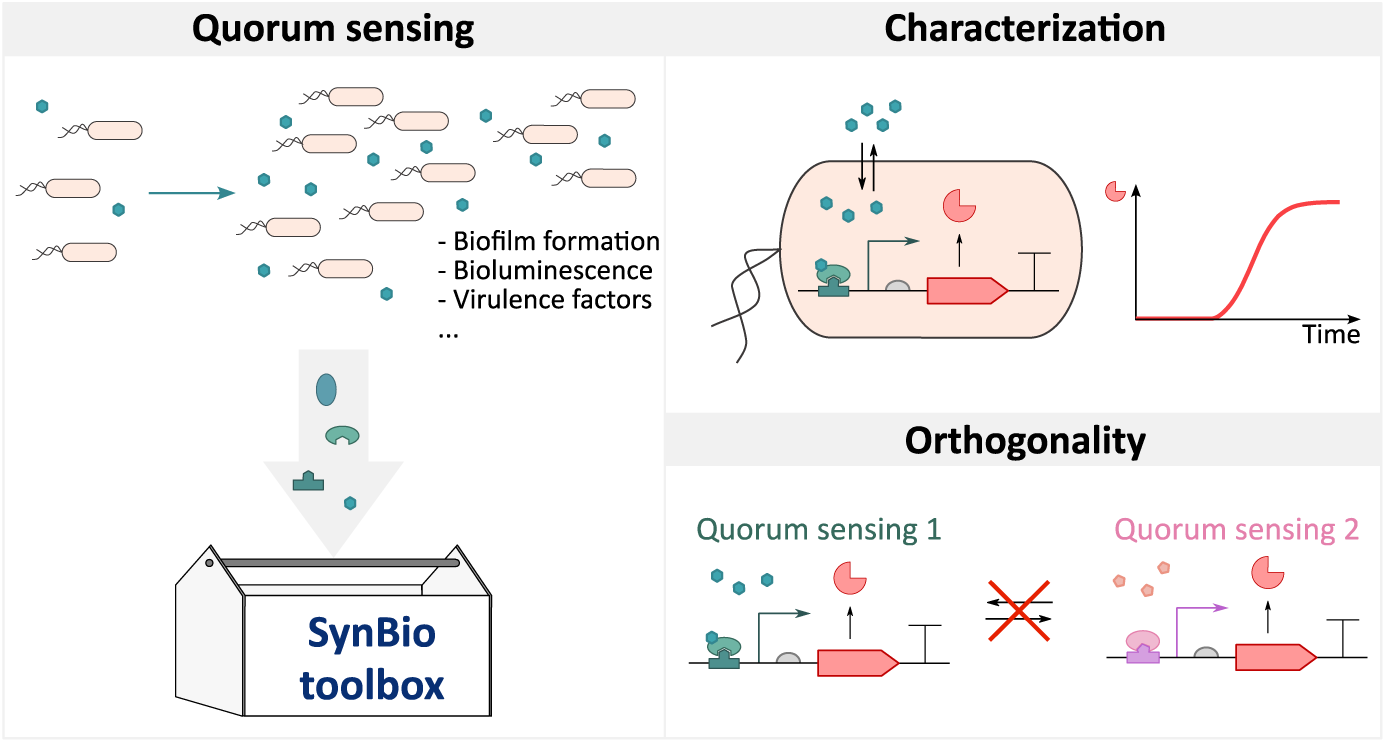
Graphical abstract of this research. Quorum sensing systems were sourced for genetic parts with potential for synthetic biology (SynBio) applications. These quorum sensing systems were characterized and their orthogonality was assessed.

Plenty of QS systems have been thoroughly characterized, albeit with the aim of unraveling the molecular mechanisms of these systems in their organism of origin. Therefore, this information might not be relevant for synthetic biologists aiming to implement these systems into genetic circuits expressed by model organisms, such as *Escherichia coli*. Assembly platforms such as the MoBioS system [11], could aid in obtaining a high-throughput and standardized manner to characterize these QS systems for synthetic biology applications. This standardized characterization could aid the fast construction of new quorum sensing-based applications by reducing the optimization time. For some applications, the combination of multiple QS systems is required to obtain the desired outcome. For example, quorum sensing-regulated co-cultivation could benefit from combining communication in two directions. Additionally, more complex genetic circuits can be obtained by layering or combining parts of different quorum sensing systems. However, due to the structural simplicity of the AI molecules and the homology of the transcription factors (TFs) within the AHL-based quorum sensing systems, issues with orthogonality might appear. Even though there has been research about the orthogonality of multiple QS systems, not all levels of crosstalk are always considered [12]. Additionally, to our knowledge, only one naturally fully orthogonal QS pair was described up until now, namely BjaI/BjaR combined with EsaI/TraR [13]. By tuning the expression level of the TF or finding mutants of the TF or promoter sequences, more orthogonal pairs were created [14,15]. However, the range of available fully orthogonal QS pairs with interesting characteristics remains limited.

In this study, two different AHL-based QS systems were investigated: the LasI/LasR and EsaI/EsaR system. The LasI/LasR system has been thoroughly researched in biomedical contexts due to its involvement in the virulence of *Pseudomonas aeruginosa* PAO1. However, its application in genetic circuitry remains limited. In contrast, the EsaI/EsaR system is frequently applied, mainly because it regulates a bidirectional promoter with opposite regulation. This gives a lot of freedom to create more diverse and complex circuitry. Besides their interesting features, these two systems might be orthogonal because of the big difference in acyl chain length between the inducers used by the two systems. In this research, we have fully characterized both QS systems (Figure 1). For this, we first analyzed the systems in the absence of their cognate synthase and with the extracellular addition of the AI molecules. Next, we introduced the synthase protein to obtain a functional, autoinducible system. Furthermore, we have investigated the full orthogonality between the two systems (Figure 1). After observing various types of crosstalk, a new transcription factor LasR and promoter P_esaR_ mutant were sought and successfully characterized.

## 2. Material & Methods

### 2.1 Strains and media

Enzymes and related products are purchased from New England Biolabs (County Road, Ipswich, MA, USA), other chemicals from Sigma-Aldrich (Brusselsesteenweg, Overijse, Belgium) unless stated differently. Protocols as described by the vendors were applied unless mentioned otherwise.

Newly assembled plasmids were introduced in One Shot^®^ Top10 Chemically Competent *E. coli* cells (Invitrogen, Carlsbad, California, USA). All experiments were performed in *E. coli* K12 MG1655 cells (mutant version of ATCC 47076, additional deletions resulting in Δ*ynaJ* Δ*uspE* Δ*fnr* Δ*cct* Δ*abgT* Δ*abgB* Δ*abgA* Δ*abgR* Δ*smrA* Δ*ydaM* Δ*ydaN* Δ*dbpA* Δ*ttcA* Δ*intR* Δ*recT* Δ*recE*).

Lysogenic Broth (LB) was used for growth during the cloning process and for preculture plates for microtiter plate experiments. This medium is composed of 10 g/L Tryptone (BioKar Diagnostics, Allonne, France), 5 g/L Yeast Extract (Becton Dickinson, Erembodegem-Dorp, Erembodegem, Belgium) and 5 g/L NaCl. 12 g/L agar (BioKar Diagnostics, Allonne, France) is added to LB to make LB-agar plates. Kanamycin was added when necessary in a 1000x dilution of the filter sterilized stock solution resulting in a final concentration of 50 µg/mL. Cultures were grown at 30°C at 200 rpm (LS-X (5 cm orbit), Adolf Kühner AG, Switzerland). Experiments were performed in MOPS EZ Rich Defined Medium (M2105) (Teknova, Hollister, CA, USA) with 0.2% glucose as carbon source. The medium was prepared according to the protocol provided by the vendor.

The following chemical inducers were added to the medium when needed: *N*-(3-oxododecanoyl)-L-homoserine lactone (O9139), *N*-(3-oxodecanoyl)-L-homoserine lactone (O9014), *N*-(3-oxooctanoyl)-L-homoserine lactone (O1764), *N*-(β-ketocaproyl)-L-homoserine lactone (K3007) from Sigma-Aldrich (Brusselsesteenweg, Overijse, Belgium). These inducers were solubilized in dimethyl sulfoxide (472301) (Sigma-Aldrich, Brusselsesteenweg, Overijse, Belgium) to a stock solution of 1 mM and stored at –80°C. Further dilutions of all inducers were done in filter sterilized MOPS EZ Rich Defined Medium.

Phosphate buffered saline (PBS) (P5368) from Sigma-Aldrich (Brusselsesteenweg, Overijse, Belgium) was used to wash the precultures to remove all autoinducers.

### 2.2 Plasmid construction

An overview of all the used plasmids is given in Supplementary Table S1. An overview of the DNA-sequence of all regulatory parts and genes is given in Supplementary Tables S2 and S3.

Plasmids for the characterization with extracellular induction and for the promoter and signal crosstalk were created in the MoBioS backbone by Golden Gate with the Type II restriction enzyme PaqCI (New England Biolabs, Inc., USA) as described in Demeester *et al.* (2023) [11,16]. The sequence of the inserted parts is given in Supplementary File 2.

For the creation of the plasmids for the autoinduced strains, each of the synthases was introduced into the backbone of the MoBioS system via Golden Gate assembly with the Type II restriction enzyme BsaI-HFv2 (New England Biolabs, Inc., USA). For LasI, a codon optimized version was used. These new synthase-MoBioS plasmids could be used as new versions of the MoBioS platform which again allowed the introduction of the different transcription factors and promoter regions via PaqCI Golden Gate. The plasmids for testing the LasR and P_esaR/esaS_ mutants were assembled using Circular Polymerase Extension Cloning (CPEC) using Q5 DNA polymerase (New England Biolabs, Ipswich, MA, USA) [17]. The linear DNA-fragments were obtained by performing polymerase chain reactions (PCR) using PrimeStar HS (Takara, Westburg, Leusden, The Netherlands). Oligonucleotides were purchased from IDT (Leuven, Belgium). A list of all oligonucleotides (IDT, Leuven, Belgium) used during this project is given in Supplementary Table S4.

For the construction of the transcription factor-sfGFP fusions, the transcription factor and a non-functional promoter (as described by Demeester *et al.* (2023) [11]) region were added as inserts for PaqCI Golden Gate with the MoBioS platform as acceptor plasmid. Afterwards, sfGFP with a glycine-rich linker (GGSGGGSG) was attached to the C-terminal of the transcription factor via CPEC.

One Shot^®^ Top10 Chemically Competent *E. coli* cells (Invitrogen, Carlsbad, California, USA) were transformed with the constructed plasmids. Successful transformation was checked through colony PCR. The plasmids of overnight growth cultures of positive colonies were prepped using the QIAprep Spin Miniprep Kit (Qiagen, Venlo, The Netherlands) and sent for Sanger sequencing to Macrogen (Macrogen Inc., Amsterdam, The Netherlands). Finished plasmids were then transformed into *E. coli* K12 MG1655.

Cells were made electrocompetent using the glycerol/mannitol density gradient wash protocol made by Warren (2011) [18]. After adding DNA, cells were electroporated at 1.8 kV, 200 Ω and 25 µF in a chilled electroporation cuvette (Bio-Rad, USA) with a 1-mm gap-width. After electroporation, 900 µL LB medium was added for resuscitation, followed by plating on LB-agar plates containing the required antibiotic.

### 2.3 *In vivo* fluorescence experiments

Precultures were inoculated in a transparent flat-bottomed 96-well plate (Greiner Bio-One, Vilvoorde, Belgium) filled with 150 µL LB medium supplemented with kanamycin when needed. These preculture plates were incubated at 30 °C for 16 to 18 hours while shaking at 800 rpm using the Digital Microplate Shaker (Thermo Fisher Scientific, Erembodegem, Belgium).

For strains expressing the synthase, an extra wash step was introduced after the preculture plate was grown overnight to remove all autoinducers that were produced during this growth. The precultures were centrifuged at 4000 rpm for 30 minutes with the Rotanta 46 RSC centrifuge (Hettich Benelux, Geldermalsen, The Netherlands) at 4°C. Supernatant was removed and the pellets were resuspended in 150 µL PBS solution. The same centrifugation and resuspension step was repeated. Afterwards a 300 times dilution of the washed (for the synthase expressing strains) and overnight grown (for the other strains) precultures plates were made into a black flat-bottomed 96-well plate (Greiner Bio-One, Vilvoorde, Belgium). The inducers were added to the desired concentration.

To follow growth and fluorescence, the plates were incubated for 24h at 30°C at an orbit of 2 mm in the Tecan Infinite M200 Pro Plate reader (Tecan Benelux, Mechelen, Belgium). Optical density at 600 nm (OD600) and fluorescence measurements were taken every 10 minutes. The used excitation and emission wavelengths for mKate2 are 588 and 633 nm, respectively. For sfGFP, 480 nm and 510 nm were used for excitation and emission, respectively.

Similarly, the assay for screening the expression library of the synthase EsaI was performed on an integrated robotic system. Cultures were incubated in the Inheco Incubator Shaker MP (integrated in the explorer G3 workstation) at 30°C and 800 rpm. The optical density and fluorescence were measured every 20 minutes in the PerkinElmer Ensight multimode plate reader (integrated in the explorer G3 workstation) using the same absorbance, excitation and emission wavelengths. For this experiment, the lid of the microtiter plate was coated with a Triton X-100 solution to reduce condensation. The Triton X-100 solution was made by mixing 20 mL ethanol with 80 mL sterile water and adding 50 µL Triton X-100. This solution was poured onto the lid and incubated for one minute. The solution was then discarded and the lid was fully dried in the fume hood before using it.

### 2.3 Data processing and statistical analysis

The data was analyzed in Python version 3.12 using Jupyter Notebooks with the pandas package. The obtained fluorescence values (*Fluo*) were normalized for optical density (*OD*_600_) per time point by Equation 1. Background fluorescence was taken into account by measuring the optical density (*OD*_600,*WT*_) and fluorescence (*Fluo*_*WT*_) of the wild type strain *E. coli* K12 MG1655 in MOPS EZ Rich defined medium. The corrected and normalized fluorescence was then averaged over the three repeats and the standard error was obtained.

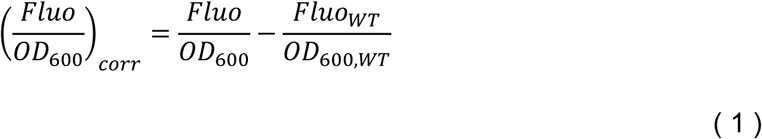

To obtain the Hill function parameters of the response curves, the Hill function for an activator (Equation 2) or a repressor (Equation 3) were fitted to the average corrected fluorescence using the weighted nonlinear least-squares algorithm (SciPy, curve_fit, Levenberg–Marquardt algorithm) [19,20].

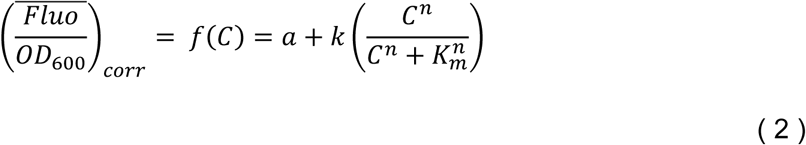

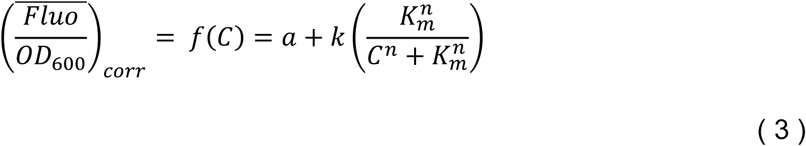

with *C* = the concentration of autoinducer in the growth medium (nM); *a* = the basal normalized fluorescent signal (leaky expression, a.u.); *k* = the relative maximum normalized fluorescent signal (a.u.); *M* = *a* + *k* = the maximum normalized fluorescent signal (a.u.); n = the Hill coefficient (cooperativity, sigmoid character); *K*_*m*_ = the Hill constant (half-maximal autoinducer concentration, nM).

For the experiments assessing orthogonality with the extracellular addition of the autoinducers, ANOVA tests followed by post-hoc Tukey test were performed with the statsmodels package in Python. In all other cases where two fluorescent outputs were compared, a Welsch two-sample t-test with unequal variance was performed (SciPy, stats) with a significance level of 0.05.

### 2.4 Confocal light scanning microscopy

For high resolution microscopy analysis, the Zeiss LSM 780 confocal scanning light microscope with Airyscan technology was used, provided by the Bio-Imaging Facility at VIB Ghent. For this, a µ-slide 8 well coverslip (ibidi, Am Klopferspitz, Martinsried, Germany) was coated with 200 µL 0.01% filter-sterilized poly-L-lysine solution (Sigma Aldrich bvba, Belgium). The cells to be visualized were grown overnight on LB medium and then subcultivated in EZ rich medium with the correct autoinducer concentration. Cultures were then 10 times diluted in transparent EZ rich medium. Next, 500 µL of the diluted cell culture was applied to each well together with 1 µL FM4-64 membrane dye solution (1 mg/mL stock) (Thermo Fisher Scientific, Erembodegem, Belgium). After incubation for 30 minutes at room temperature, the wells were pipetted dry. Subsequently, a few droplets glycerol (Chem-Lab Analytical BVBA, Belgium) were added as mounting medium. In the microscopic analysis, both the red (membrane) and green (proteins) channels were visualized. Microscopy images were analyzed using Fiji (an image processing package of ImageJ). To analyze the localization of the proteins, an intensity plot of the cross-section of single cells was made.

## 3. Results and discussion

### 3.1 Characterization of the LasR– and EsaR-based quorum sensing system with extracellular induction

The LasI/LasR and EsaI/EsaR system (Figure 2), belonging to the LuxI/LuxR-family of QS systems, both possess promising characteristics for synthetic biology applications. The molecular mechanism of the LasI/LasR system in its host organism *Pseudomonas aeruginosa* PAO1 has been completely unraveled [21–26], providing an enormous source of information for optimizing its application. The system contains a unidirectional promoter that is upregulated by LasR at high cell densities. The EsaI/EsaR system, originating from *Pantoea stewartii* ssp. *stewartii*, contains a bidirectional promoter with opposite regulation [27], providing interesting building blocks for new genetic circuitry. Furthermore, the autoinducers (AIs) used by both systems differ a lot in acyl chain length, compared to other quorum sensing (QS) systems from the LuxI/LuxR-family, indicating possible orthogonality. Namely, the LasI synthase produces 3-oxo-dodecanoyl homoserine lactone (3OC12-HSL), whereas EsaI synthesizes 3-oxo-hexanoyl homoserine lactone (3OC6-HSL). Although AHLs are generally assumed to freely diffuse through the cell membrane, it was demonstrated that this is not the case for AHLs with longer acyl chains, such as 3OC12-AHL. Besides diffusion, efflux pumps aid in transporting the AHL molecules outside of the cell [28]. Upon detection of 3OC12-HSL, the transcription factor LasR will dimerize and the activated protein will bind to the *las*-box, a 20 base pair palindromic sequence centered 40.5 base pairs upstream of the transcription start site of the LasI-promoter, and recruit RNA-polymerase to the core promoter leading to transcription (Figure 2A) [21,23,25]. In the EsaI/EsaR system, the dimeric transcription factor EsaR functions both as a transcriptional activator and repressor of a bidirectional promoter in the absence of the inducer, resulting in the activation and repression of P_esaS_ and P_esaR_, respectively (Figure 2B). The shared *esa*-box, the 20 base pair palindromic transcription factor binding site of EsaR, overlaps with the –10 box of P_esaR_ leading to steric hindrance of the RNA-polymerase. On the other hand, the *esa*-box is situated about 60 base pairs upstream of the transcription start site of P_esaS_, where bound EsaR acts as an activator. In the presence of 3OC6-HSL, at high cell densities, EsaR will undergo a conformational change and is no longer capable of binding the bidirectional P_esaR/esaS_ promoter [27].

**Figure 2.**
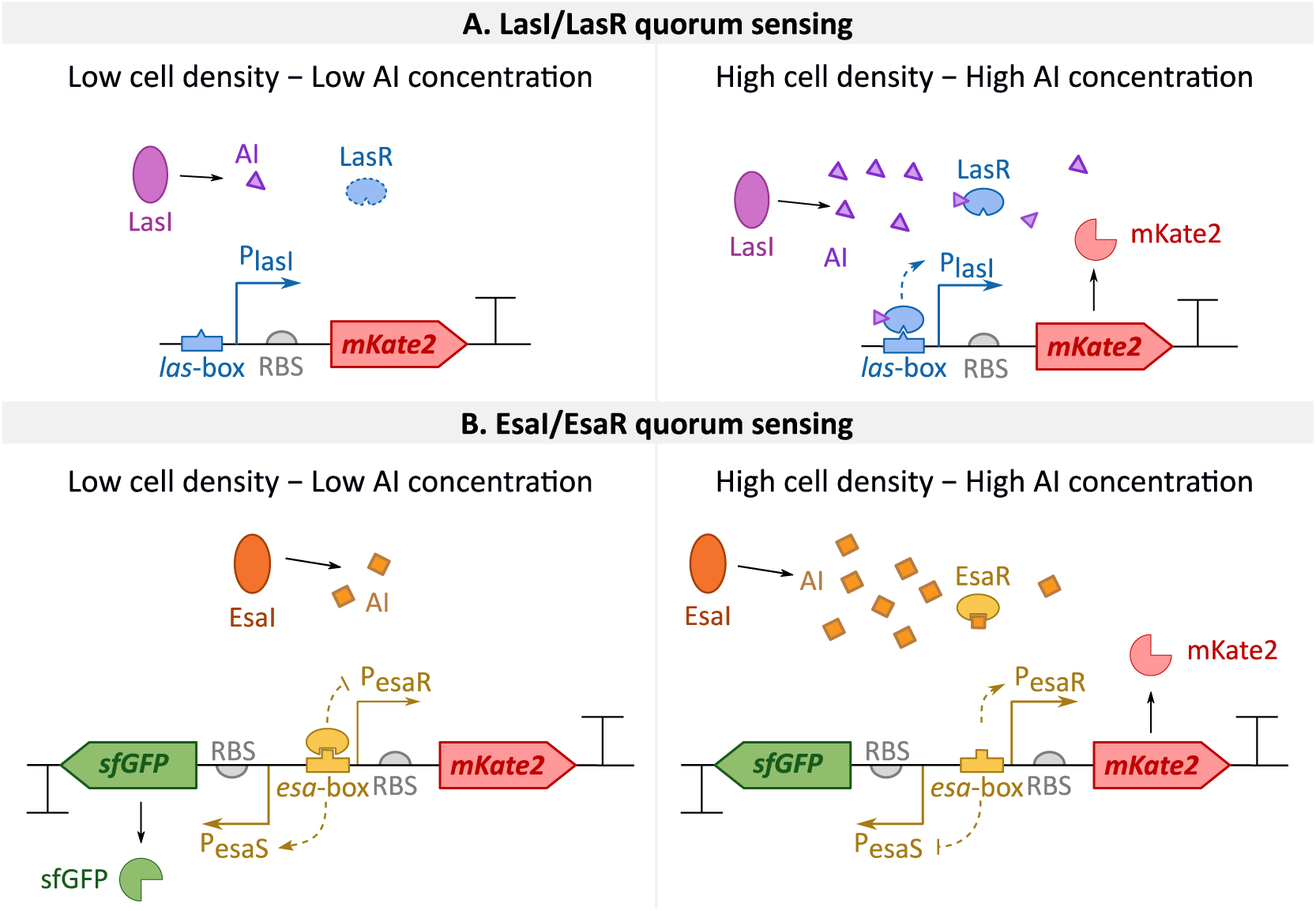
Overview of the principle of the two used quorum sensing systems in this research. **A.** Principle of the LasI/LasR system [25]. The synthase LasI produces an autoinducer (AI). At high cell densities, this AI will reach high concentrations and form a complex with LasR. This transcription factor will now activate the P_lasI_ promoter, here regulating expression of the red fluorescent protein mKate2. LasI and LasR are constitutively expressed. **B.** Principle of the EsaI/EsaR system [27]. The synthase EsaI produces the autoinducer. At low concentrations of this AI molecule, EsaR binds the P_esaR/esaS_ promoter and activates and represses P_esaS_ and P_esaR_, respectively, leading to production of the green fluorescent protein sfGFP. At high concentrations EsaR releases the promoter and the activity reverses, leading to mKate2 production from P_esaR_. EsaI and EsaR are constitutively expressed. Genetic parts are depicted according to SBOL conventions [29,30]. RBS = ribosome binding site.

In order to make predictable genetic circuits and minimize optimization times, a thorough characterization of the used QS systems is required. By breaking down the QS systems into various levels of interaction, more insight can be obtained. First, the interaction between the AI, TF and corresponding promoter P_QS_ is characterized in the absence of the synthase. By replacing the synthase with the addition of different AI concentrations to the growth medium while retaining constitutive expression of the transcription factor, a response curve can be created (Figure 4A). Furthermore, this was done for three different expression levels of the transcription factor, as this is known to influence the response curve [9,31,32]. As such, three different ribosome binding site (RBS) sequences, referred to as RBS-low, RBS-medium and RBS-high, were designed *in silico* for each transcription factor with the Salis RBS calculator, theoretically spanning a range of strengths [33]. The corresponding *in vivo* expression strength of each RBS was measured by creating a transcription factor-sfGFP fusion protein, allowing to use green fluorescence intensity as a proxy for protein levels of the transcription factor (Figure 3A). The quantification of this fusion protein was done both in the absence and presence of the corresponding AI, since some transcription factors from the LuxR-family require the presence of their AI to form a stable protein complex and will be degraded in its absence.

**Figure 3.**
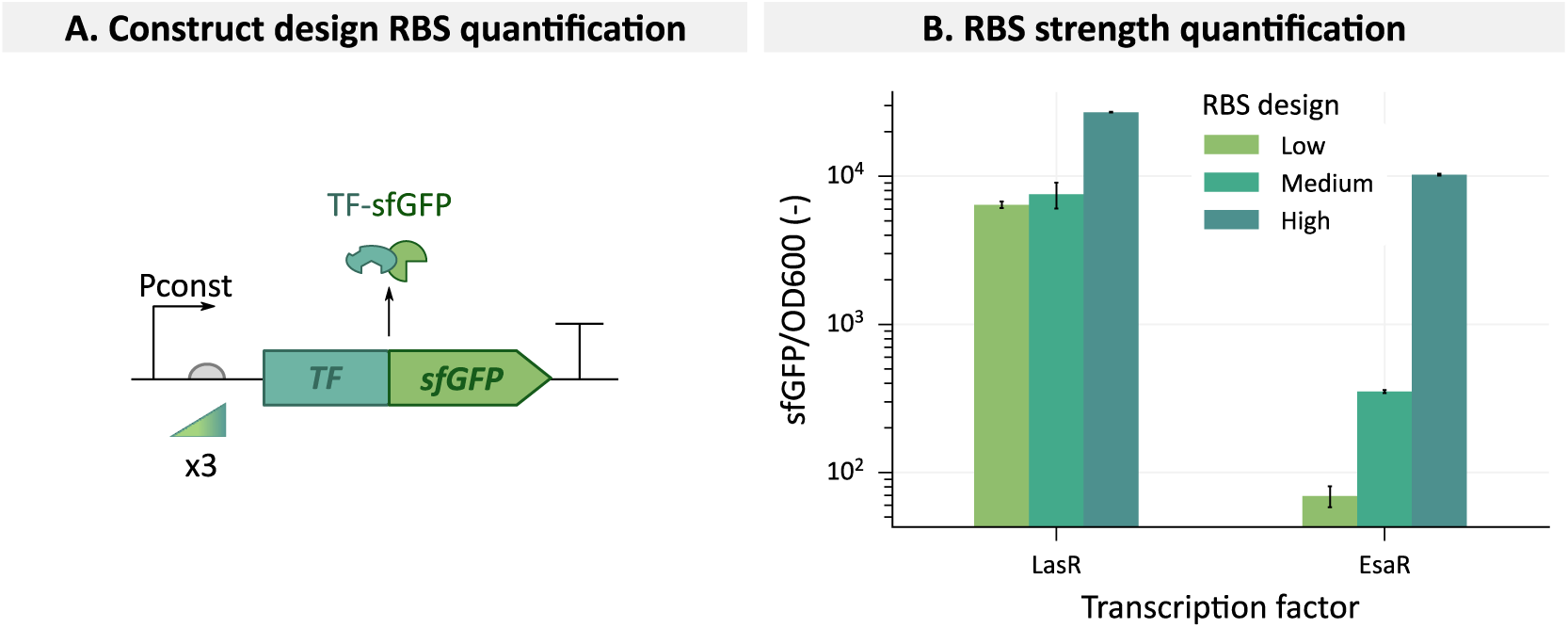
Quantification of the expression level of the transcription factor-sfGFP fusion, controlled by three different ribosome binding site (RBS) sequences for both LasR and EsaR. **A.** Construct design of the transcription factor (TF)-sfGFP fusions. **B.** Quantification of the sfGFP-fusion proteins. Fluorescent values were obtained in the stationary phase and normalized for cell growth determined by optical density at 600 nm (OD600). Bars represent the mean and error bars the standard error of the mean based on three biological replicates.

**Figure 4.**
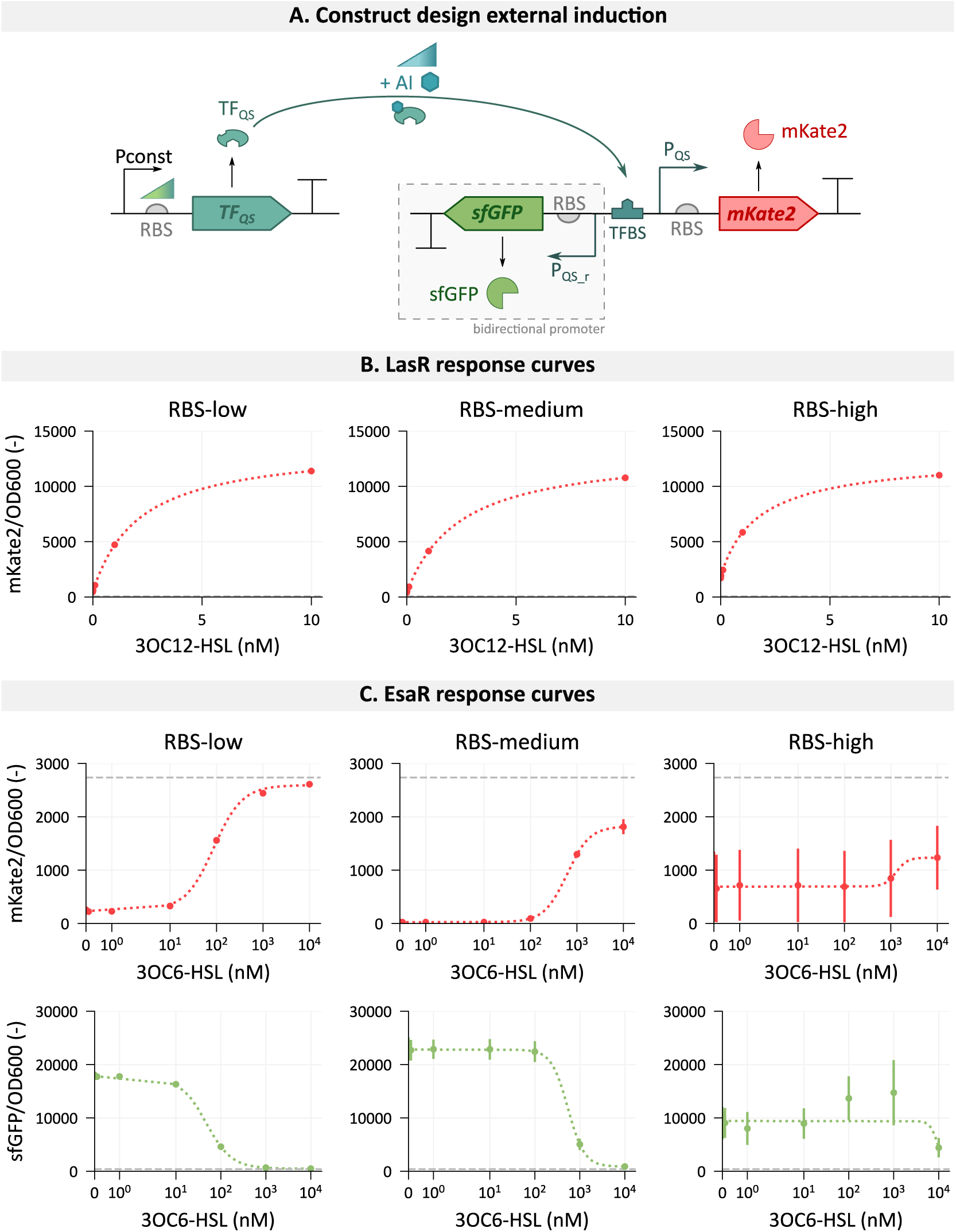
Overview of the characterization of the LasR– and EsaR-based quorum sensing (QS) systems with the extracellular addition of their respective autoinducer molecule. **A.** Construct design for obtaining the response curve at different RBS-strengths for the transcription factor. In response to the presence of autoinducers (AIs), the transcription factor (TF), will have an effect on the quorum sensing-regulated promoter (P_QS_), regulating the expression of a fluorescent reporter protein. For the EsaR-based QS system the bidirectional promoter is indicated by P_QS_ for P_esaR_ and P_QS_r_ for P_esaS_. Genetic parts are depicted according to SBOL conventions [29,30]. **B.** Response curves of the three different variants of the LasR system with its autoinducer 3OC12-HSL. **C.** Response curves of three variants of the EsaR system with its autoinducer 3OC6-HSL. The red fluorescent protein mKate2 is a proxy for P_esaR_ promoter activity, whereas green fluorescence by sfGFP quantifies P_esaS_ promoter activity. Fluorescent values are normalized for cell growth determined by optical density at 600 nm (OD600) and obtained during the stationary growth phase. Error bars represent the standard error for three biological replicates. The gray dashed line depicts the basal promoter activity in the absence of the transcription factor. Pconst = constitutive promoter, RBS = ribosome binding site, TF_QS_ = transcription factor of a quorum sensing system, AI = autoinducer.

For the LasR-based QS system, the three generated RBS sequences have an *in silico*-predicted translation initiation rate of 478, 5,847 and 26,568 a.u. for RBS-low, RBS-medium and RBS-high, respectively [33]. However, the *in vivo* quantification shows large deviations from these predictions. RBS-low and RBS-medium result in similar LasR-sfGFP expression levels, whereas the measured green fluorescence was higher for RBS-high albeit not as much as predicted *in silico* (Figure 3B). Interestingly, no difference in expression level could be observed in the presence or absence of the AI, even though LasR was thought to require its AI to aid in protein folding (Supplementary Figure S1A) [21]. Only with the addition of 10 µM 3OC12-HSL, a decrease in fluorescence could be observed for the highest expression level of LasR, although this could be explained by the large metabolic burden the strain experiences. For each of the three expression levels, the response curve was obtained by measuring mKate2 production, regulated by P_lasI_, with the addition of different 3OC12-HSL concentrations, ranging from 0.01 to 10 nM. The corresponding response curves (Figure 4B) only show small differences between the three variants, which can be explained by the small difference in RBS strength. In all cases, the activation of LasR happens at low 3OC12-HSL concentrations, around 1-2 nM. The full details of the response curves together with ID-sheets of this 3OC12-HSL bioswitch can be found in Supplementary File 2, as was described in Demeester *et al.* (2023) [11].

It can be observed that in the absence of the AI, there is already an increase in P_lasI_ activity compared to the promoter activity without LasR expression. This increase is the most distinct for the highest expression level of LasR (Figure 4B). This indicates that some LasR is present in an active confirmation without 3OC12-HSL. This result, combined with the earlier observation of similar LasR-sfGFP levels in the absence of 3OC12-HSL, raises questions about the actual molecular mechanism of LasR. Further microscopic analysis indicates that the protein was homogeneously present in the cytoplasm of the cells both in the presence and absence of 3OC12-HSL (Supplementary Figure S2). This does not agree with the general assumption that LasR, in the absence of its AI, is nonfunctional and present in the insoluble fraction, such as inclusion bodies [21,34,35]. Possibly LasR folds into an inactive soluble protein that is activated upon signal binding as was demonstrated by Sappington *et al.* (2011) [25]. At high protein levels, some of this soluble LasR might succeed in binding and activating P_lasI_.

Similar to the characterization of the LasR-based QS system, the response curves of EsaR were obtained at three different expression levels, defined by the RBS strength. Similar RBS strengths were sought for controlling the translation of EsaR as for LasR, resulting in translation initiation rates of 529, 3,115 and 29,036 a.u. for the low, medium and high strength RBS, respectively. In this case, the *in vivo* strength was more in line with the *in silico* design, covering a large range of possible expression levels of this transcription factor (Figure 3B). Additionally, the presence of the AI did not influence the EsaR concentration (Supplementary Figure S1B).

To follow the activity of the bidirectional EsaR-regulated promoter, two different fluorescent reporter proteins were used, namely mKate2 and sfGFP, for P_esaR_ and P_esaS_ activity, respectively. The increase in RBS strength results in a shift of the response curves of P_esaR_ and P_esaS_ to higher 3OC6-HSL concentrations (Figure 4C). Additionally, the leaky expression and maximal promoter activity are influenced by the EsaR expression level. However, with the highest expression level of EsaR, the growth is severely hampered, resulting in large biological variability on the corresponding response curves. This is probably due to the burden caused by overexpressing EsaR, as was observed earlier by Shong *et al.* (2013) [36]. A summary of the parameterized response curves can be found in (Supplementary File 2).

In nature, a broad range of AHL-concentrations can be found, going from 1 nM to 50 µM [37]. For 3OC12-HSL produced by *Pseudomonas aeruginosa* PAO1, a range of 0.5 to 5 µM was detected in laboratory cultures [38] and up to 600 µM in biofilms [39]. To induce exopolysaccharide production in *Pantoea stewartii*, a concentration of 2 µM 3OC6-HSL was required [40]. In our synthetic system, around 1-2 nM was required for activating LasR. Hence, this concentration is more than 100 times lower than what is generally observed in nature. The main difference is that in our synthetic system both the transcription factor and the promoter are expressed from a plasmid instead of genomic. Generally, it can be assumed that higher LasR levels are obtained. With LasR functioning as an activator, this bigger pool of unbound LasR has a higher probability of binding lower AHL-concentrations [32]. However, the opposite would be expected for the EsaR-based system in which EsaR releases the promoter region in the presence of the AI. Hence, higher EsaR and promoter levels, compared to genomic expression, would require more AI to titrate the EsaR away and induce a response. However, this is not confirmed by our data, in which concentrations ranging from 46 to 600 nM are required to switch the promoter activity compared to 2 µM 3OC6-HSL in nature (Supplementary File 2) [40].

### 3.2 Characterization of the quorum sensing systems with autoinduction

The obtained response curves provide insight into the interplay between the transcription factor, autoinducer and promoter. For both LasR and EsaR, the lowest expression levels resulted in functional, responsive systems with minimal burden on the host. Therefore, further characterization of the systems was done with the lowest expression level for each of the transcription factors. Subsequently, the corresponding synthase was added to each of the plasmids regulated by the constitutive J23108 promoter and the Bba_B0031 RBS from the iGEM parts registry, resulting in a LasI/LasR and a EsaI/EsaR quorum sensing strain, referred to as LasQS and EsaQS, respectively (Figure 5A). Similar as for the response curves, the strains LasQS and EsaQS were monitored for mKate2 or mKate2 and sfGFP expression, respectively.

**Figure 5.**
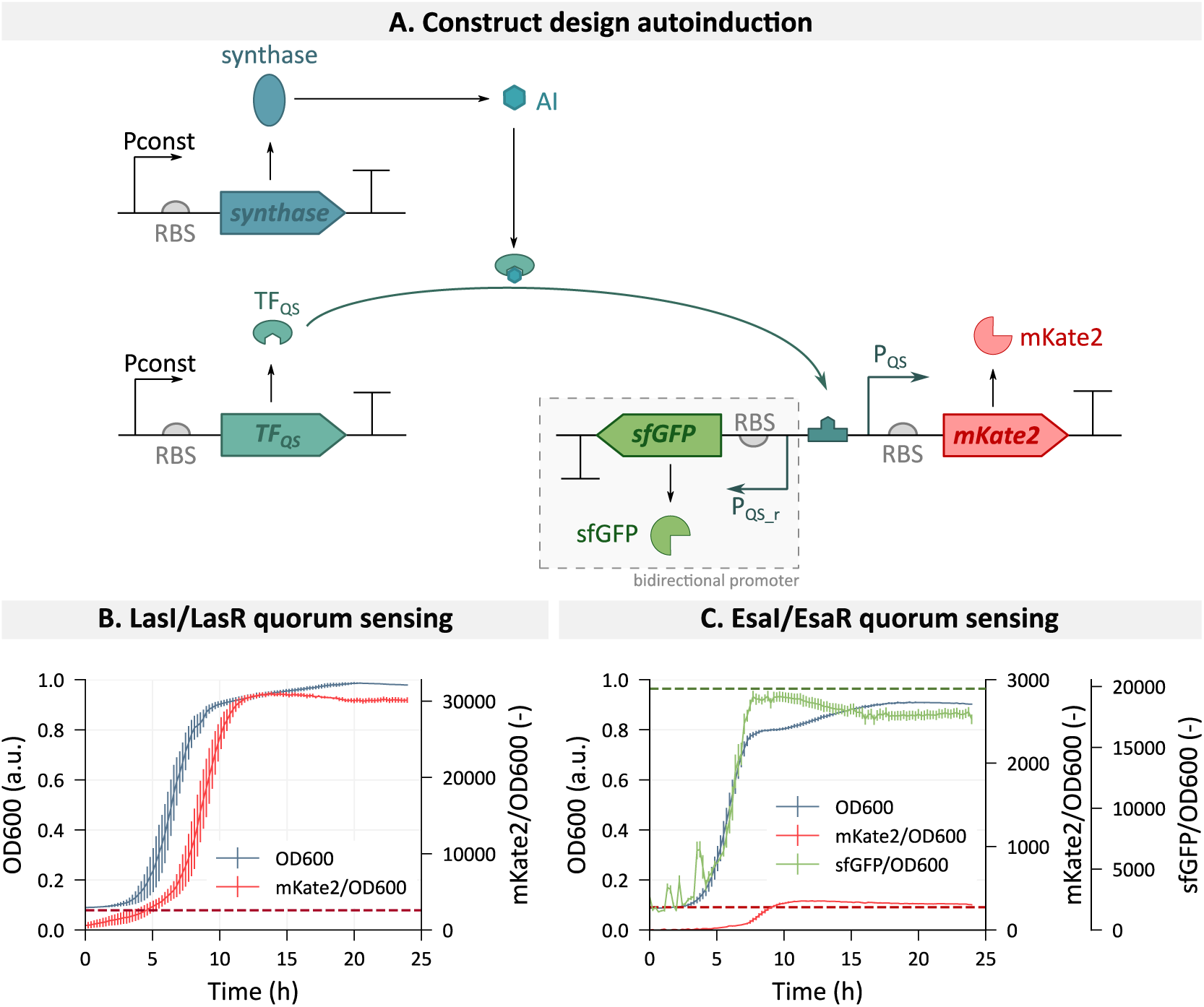
**A**. Construct design for assessing the functionality of the quorum sensing (QS) strains. The autoinducer (AI) produced by the synthase will bind to the quorum sensing transcription factor (TF_QS_), thereby influencing its activity. This will result in a change in activity of the promoter it regulates (P_QS_), measured by fluorescent protein production. Genetic parts are depicted according to SBOL conventions [29,30]. **B.** P_lasI_ activity, quantified by red fluorescent protein mKate2 production, when autoinduced by LasR in the presence of autoinducer produced by LasI. Both LasR and LasI are constitutively expressed. The red dashed line depicts the promoter activity of P_lasI_ with LasR constitutively expressed in the absence of 3-oxo-dodecanoyl homoserine lactone (3OC12-HSL). **C.** P_esaR_ and P_esaS_ activity, quantified by red fluorescent protein mKate2 and green fluorescent sfGFP production, respectively. The strain constitutively expresses EsaR, which, in the absence of its AI, binds the promoter region leading to an activation of P_esaS_ and repression of P_esaR_. Additionally, the respective synthase EsaI is constitutively expressed. The green and red dashed line depict the promoter activity of EsaR-bound P_esaR_ and P_esaS_, respectively, in the absence of 3-oxo-hexanoyl homoserine lactone (3OC6-HSL). Fluorescent values are normalized for cell growth determined by optical density at 600 nm (OD600). Error bars represent the standard error for three biological replicates. Pconst = constitutive promoter, RBS = ribosome binding site.

A functional LasQS strain was obtained with the autoinduced expression of mKate2 regulated by P_lasI_ as can be seen in Figure 5B. The normalized fluorescence is produced at levels that even surpass the maximal promoter activity observed in the response curves (Figure 4B). The normalized mKate2 fluorescence shows a slight delay compared to the growth, caused by 3OC12-HSL accumulation, fluorescent protein production time and/or maturation. In contrast, the EsaQS system does not switch to the induced phenotype as was expected (Figure 5C). The sfGFP levels stay high, while mKate2 is only produced at levels that correspond to leaky expression. This indicates that the EsaI synthase is not sufficiently expressed to result in the required 3OC6-HSL concentration for induction.

To create a functional EsaQS-strain, a library of 24 RBS-variants was created with the Salis RBS library Calculator to regulate EsaI production [33] (Supplementary Table S5). It was demonstrated earlier that adapting the expression strength of the synthase is a successful strategy for tuning the timing of induction [41,42]. Additionally, an LVA-degradation tag was added to sfGFP [43]. Since sfGFP is a highly stable protein, this tag is required to see a decrease in protein production when the EsaQS-strain switches to the induced state and P_esaS_ gets deactivated. Furthermore, the RBS regulating EsaR expression was exchanged for RBS-medium, since the response curve of the strain expressing EsaR at this level reaches the highest sfGFP levels (Figure 4C). This to ensure that sfGFP was still produced within the measurable range despite the increased turnover by the degradation tag. The EsaI-RBS-library was screened and the results could be divided into three groups (Figure 6 and Supplementary Figures S3-8). In Group 1, no increase in mKate2 could be observed and the decrease in sfGFP could only be attributed to the degradation tag. Hence, there is not enough 3OC6-HSL accumulation to induce the switch, most likely because of insufficient EsaI expression. In contrast, strains from Group 2 did not produce sfGFP in the detectable range and the mKate2 increased rapidly. It is expected that these strains have too high EsaI expression, resulting in the rapid accumulation of 3OC6-HSL and induction at the start of the growth. Interestingly, high expression levels of EsaI, resulted in visible detrimental effects on the cell growth such as an increased lag phase and lower maximal growth rate (Figure 6B and Supplementary Figures S5-8), possibly caused by either recombinant protein overexpression or the actual 3OC6-HSL production that tapped into the cellular resources. In a different study, this burden was also observed in a library expressing the synthase LuxI at different levels [42]. Finally, in Group 3 the desired phenotype could be observed in which mKate2 expression follows up on the sfGFP production. This phenotype could be linked to the RBS-sequences with a theoretical translation initiation rate of 45 or 103 a.u., corresponding to the theoretically weakest RBS sequences in the library. The RBS sequence with an *in vivo* predicted translation initiation rate of 45 a.u. will be used for the regulation of *esaI* translation in the rest of this research.

**Figure 6.**
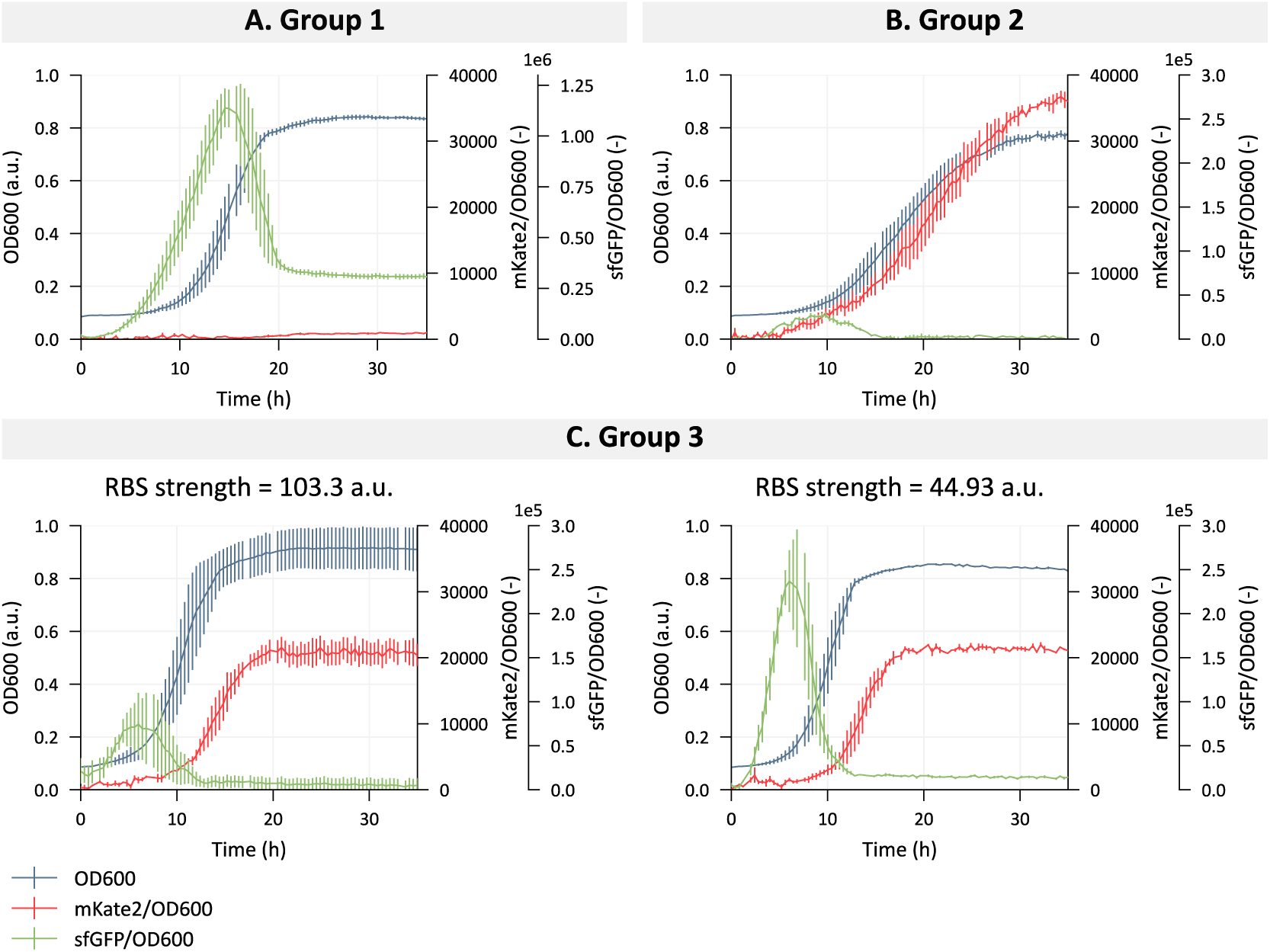
Selection of strains from RBS-library for tuning the expression of EsaI in the EsaI/EsaR quorum sensing strain. The strains are grouped based on the output. **A.** Group 1 retained maximal sfGFP expression, while only producing leaky mKate2. **B.** In Group 2, mKate2 reaches maximal values but no sfGFP can be observed. **C.** In Group 3, mKate2 production follows sfGFP production, with a switch during the early exponential phase. The theoretical translation initiation rate of the RBS sequences in the two strains in this group is provided above the plot. Fluorescent values are normalized for cell growth determined by optical density at 600 nm (OD600). Error bars represent the standard error for three biological replicates. Note that the scale of the y-axis for sfGFP/OD600 (-) differs in Group 1 compared to the other two groups.

### 3.3 Orthogonality of the two characterized quorum sensing systems

The simultaneous use of the two QS systems ideally occurs without any interference between the two systems. However, crosstalk can occur on three levels: promoter crosstalk in which the TF is able to regulate a non-canonical promoter (Figure 7A); signal crosstalk caused by low specificity of the TF for the AI molecules (Figure 7B); or synthase crosstalk where the synthase produces a mixture of AIs (Figure 7C). Due to the high similarity between the QS systems belonging to the LuxR-family, crosstalk on any of these levels is likely. However, once crosstalk is identified, orthogonality can be sought by engineering the relevant protein or promoter sequences. The crosstalk between the LasI/LasR and EsaI/EsaR systems was fully assessed at each level. By replacing the presence of the synthase protein with the extracellular addition of the AIs, signal and promoter crosstalk can be analyzed while excluding the influence of possible synthase crosstalk on the results.

**Figure 7.**
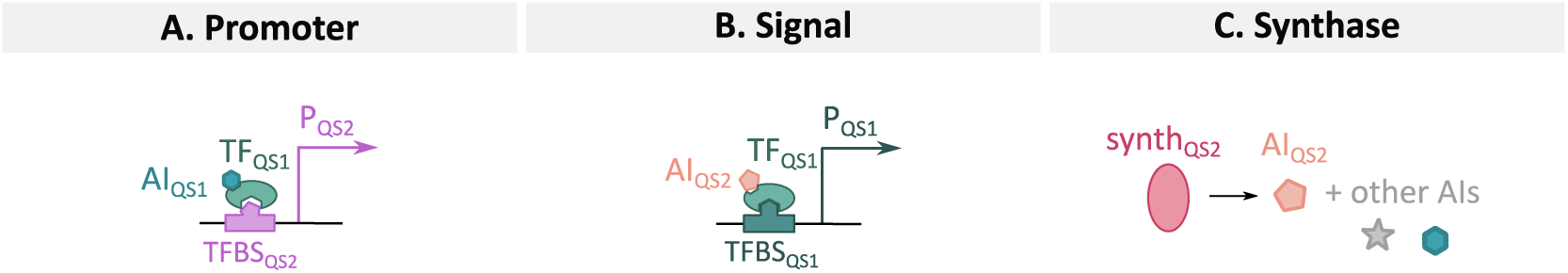
Illustration of the three types of crosstalk that can occur between different acyl-homoserine lactone-based quorum sensing systems. **A.** Promoter crosstalk: The transcription factor of one system (TF_QS1_) is able to bind and influence activity of the promoter of another system (P_QS2_). **B.** Signal crosstalk: The transcription factor of one system can bind to the autoinducer of the second system (AI_QS2_) due to the promiscuity of the transcription factor. **C.** Synthase crosstalk: The synthase of one system (synth_QS2_) produces a range of different AI molecules. TFBS = transcription factor binding site.

#### Promoter crosstalk

The specificity of a TF for its transcription factor binding site (TFBS) is defined by the DNA-binding domain of the transcription factor and the TFBS sequence located in the promoter region. For TFs from the LuxR-family, this TFBS is a palindromic sequence of about 20 base pairs with high homology between the different systems (Figure 8) [23,44,45]. Therefore, non-specific binding of a TF to a non-cognate TFBS is likely to occur. To assess this possible promoter crosstalk, each transcription factor, LasR and EsaR, was combined with its non-cognate promoter, P_esaR/esaS_ and P_lasI_, respectively. Their cognate AI was added in a range of concentrations from 0.01 nM to 10 µM and promoter activity was again assessed by measuring the fluorescent output of the reporter proteins (Figure 9A). Additionally, these results were compared with strains containing the quorum sensing-dependent promoter regulating the reporter protein, but lacking the necessary transcription factor, as a quantification of the basal expression level. Since EsaR works as a repressor of P_esaR_, high promoter activity is expected in the absence of EsaR.

**Figure 8.**
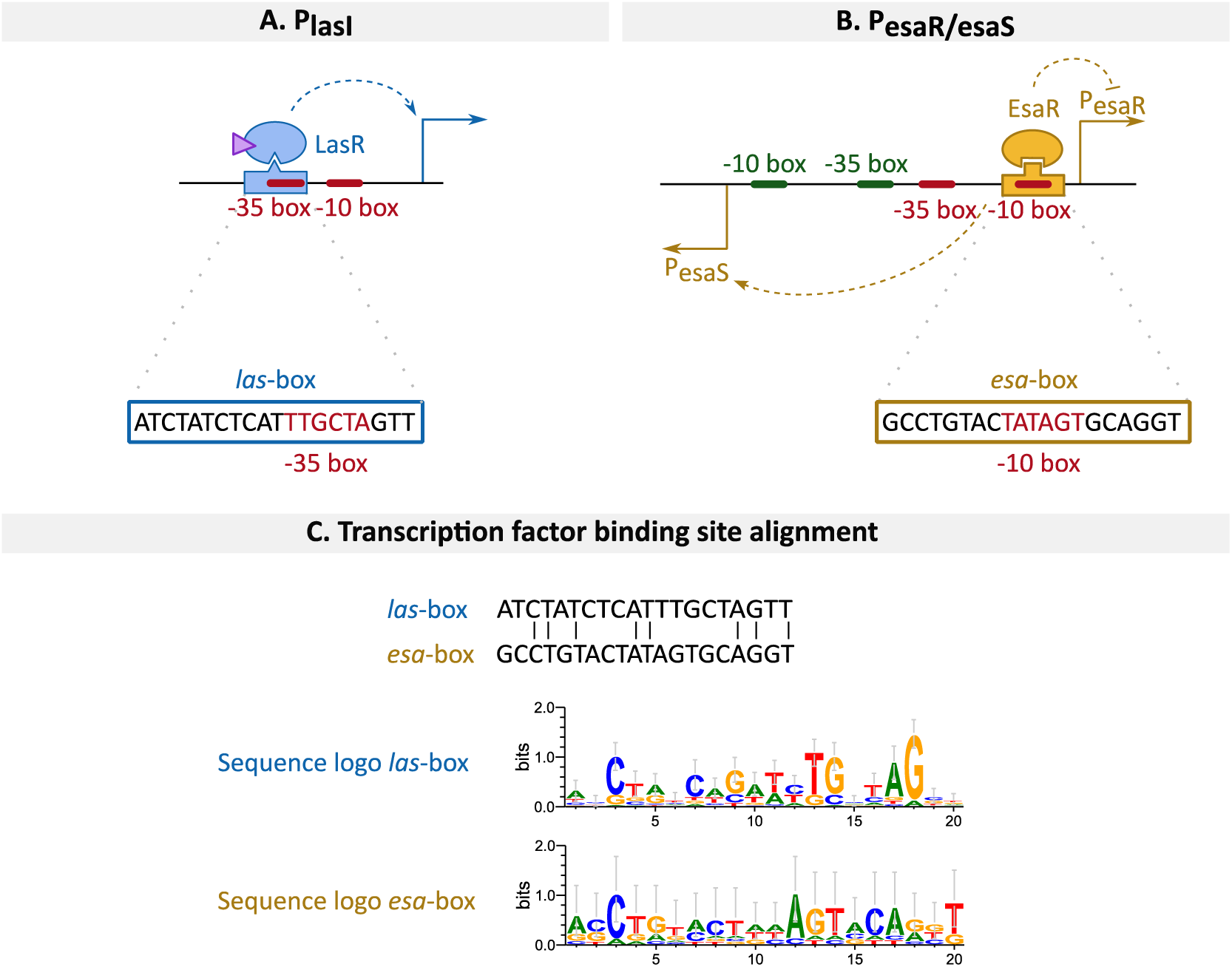
**A**. Promoter architecture of the P_lasI_ promoter with the *las*-box, the transcription factor binding site of LasR. This transcription factor, when bound to its autoinducer, will function as an activator. The sequence of the *las*-box is highlighted. **B.** Promoter architecture of the P_esaR/esaS_ promoter with the *esa*-box, the transcription factor binding site of EsaR. This transcription factor functions as an activator of P_esaS_ and repressor of P_esaR_ simultaneously. The sequence of the *esa*-box is highlighted. **C.** Comparison of the sequences of the *las*– and *esa*-box used in this research. Additionally, the sequence logos of both boxes are depicted for further insight into the sequence conservation [46–48].

**Figure 9.**
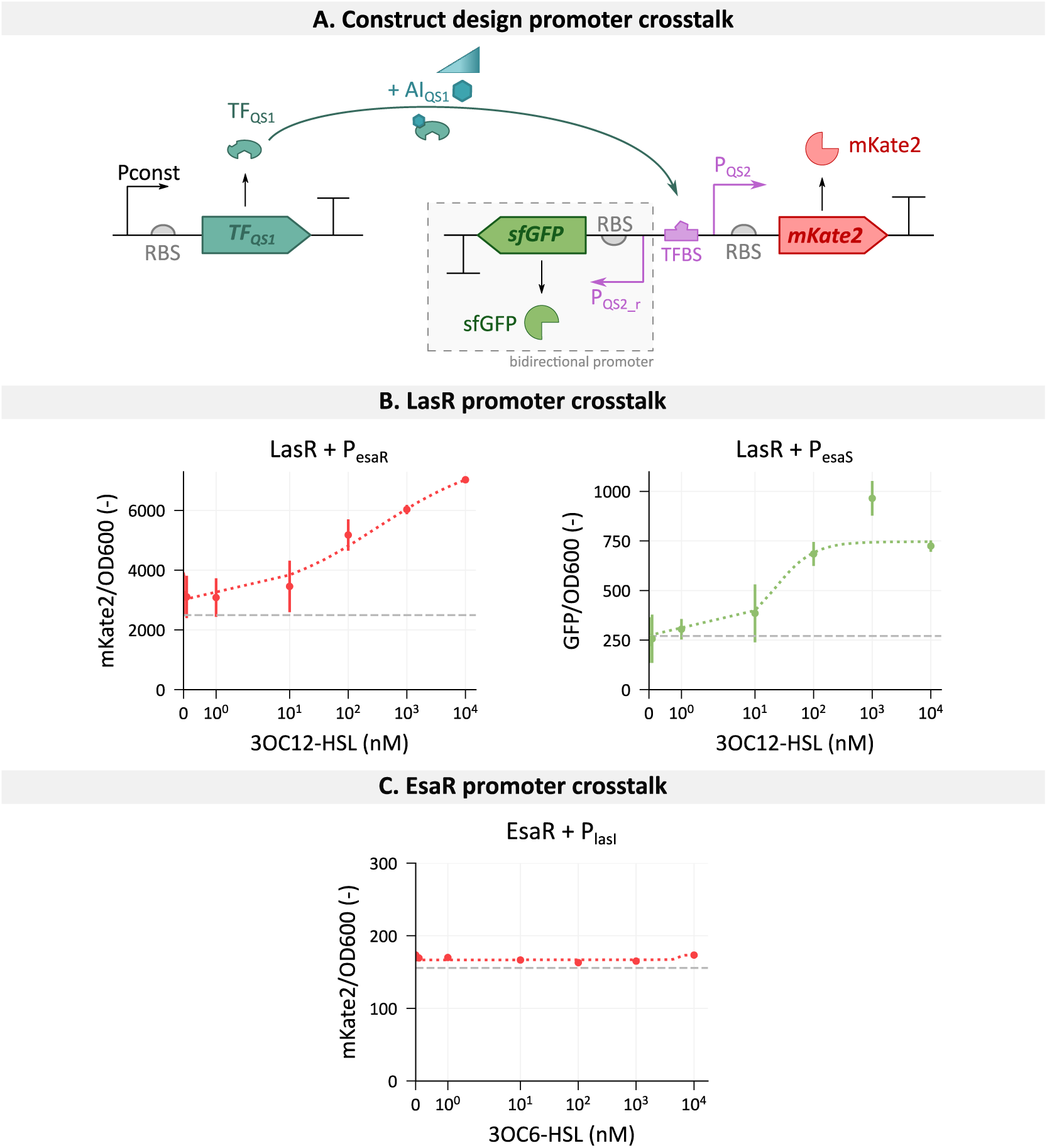
Assessment of the promoter crosstalk between the two quorum sensing (QS) systems. **A.** Construct design for checking promoter crosstalk. A constitutively expressed transcription factor of the first QS system (TF_QS1_) is combined with the quorum sensing-regulated promoter of the second system (P_QS2_), which can be bidirectional (P_QS2_r_). Different concentrations of the autoinducer of system one (AI_QS1_) are added to the medium and the change in promoter response is followed by measuring the fluorescence intensity of the reporter proteins. Genetic parts are depicted according to SBOL conventions [29,30]. **B.** Crosstalk assessment of the transcription factor LasR with the P_esaR_ and P_esaS_ promoter of the EsaR-based quorum sensing system. The promoters P_esaR_ and P_esaS_ regulate the transcription of *mKate2* and *sfGFP*, respectively, to allow quantification of their promoter activity. **C.** Crosstalk assessment of the transcription factor EsaR with the P_lasI_ promoter of the LasR-based quorum sensing system. The promoter P_lasI_ regulates the transcription of *mKate2* to allow quantification of its promoter activity. The gray dashed lines depict the basal promoter activity in the absence of the transcription factor. Fluorescent values are normalized for cell growth determined by optical density at 600 nm (OD600) and obtained during the stationary growth phase. Error bars represent the standard error for three biological replicates, except for the EsaR promoter crosstalk measurement with 0 nM 3OC6-HSL, where only two replicates remained. Pconst = constitutive promoter, RBS = ribosome binding site. An overview of the statistical analysis is given in Supplementary Table S6-8.

The *esa*-box, situated in the P_esaR_ promoter, overlaps with its –10 box, thereby causing steric hindrance for the RNA-polymerase (Figure 2B). Hence, in the absence of EsaR, an active promoter can be observed. However, when combining with active LasR, P_esaR_ gets activated to levels surpassing the lack of EsaR-repression (Figure 9B). Similarly, P_esaS_ activation increases with increasing 3OC12-HSL concentration, corresponding to more activated LasR. However, the effect remains limited compared to the basal promoter activity. ANOVA tests confirmed a significant effect of increasing 3OC12-HSL on P_esaR_ and P_esaS_ activity (p = 0.000847 and p = 0.000194, respectively). Full statistical details can be found in Supplementary Tables S6 and S7. Despite the observed LasR promoter crosstalk, it is likely that in a system with EsaR, the higher affinity of EsaR for its cognate promoter would outcompete possible binding of LasR. Similarly, LasR has a higher affinity for its own cognate promoter P_lasI_, as can be seen by the lower 3OC12-HSL concentration required to obtain sufficient active LasR to activate the promoter (Figure 4B). Therefore, in the presence of P_lasI_, the promoter crosstalk is expected to be lower.

In contrast with LasR, no significant effect of 3OC6-HSL binding to EsaR on the activity of P_lasI_ could be revealed (Figure 9C and Supplementary Table S8). Additionally, the observed leaky expression corresponds to the same mKate2-expression of a strain containing only the P_lasI_, in the absence of LasR. Therefore, we can conclude that there is no EsaR-P_lasI_ promoter crosstalk.

#### Signal crosstalk

To assess signal crosstalk, different concentrations ranging from 0.01 nM to 10 µM of the cognate and non-cognate AI molecule were added to the strains LasR-P_lasI_ or EsaR-P_esaR/esaS_ (Figure 10A). Additionally, to verify the specificity of the transcription factors, two more AHLs were checked, namely 3-oxo-octanoyl homoserine lactone (3OC8-HSL) and 3-oxo-decanoyl homoserine lactone (3OC10-HSL).

**Figure 10.**
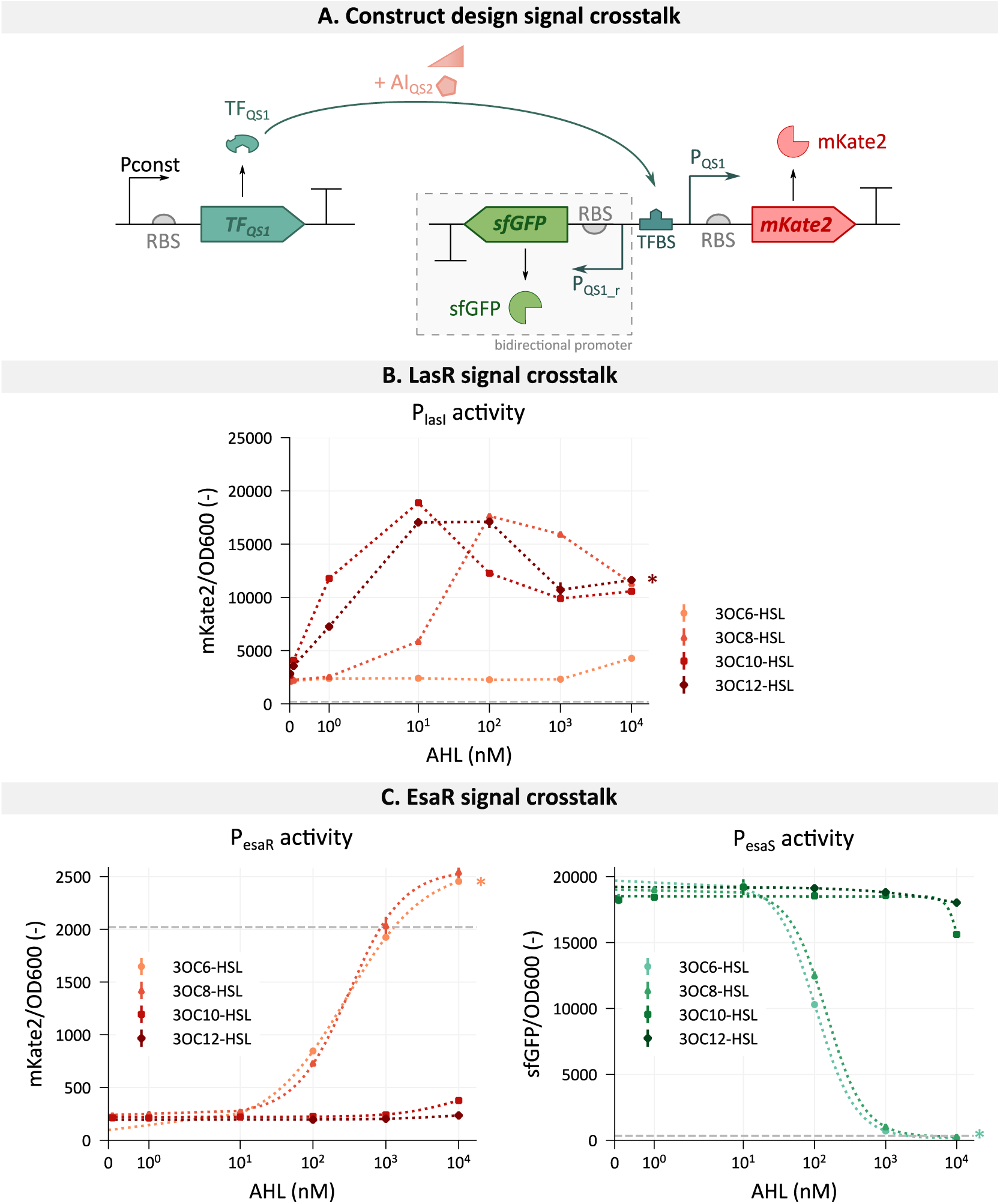
Assessment of the signal crosstalk between the two quorum sensing (QS) systems. **A.** Construct design for checking signal crosstalk. A constitutively expressed transcription factor (TF_QS1_) is combined with its own quorum sensing-regulated promoter (P_QS1_), which can be bidirectional (P_QS1_r_). Different concentrations of the autoinducer of a second quorum sensing system (AI_QS2_) are added to the medium and the change in promoter response is followed by measuring the fluorescence intensity of the reporter proteins. Genetic parts are depicted according to SBOL conventions [29,30]. **B.** Crosstalk of the transcription factor LasR with different acyl-homoserine lactone molecules (AHLs). The promoter P_lasI_ regulates the transcription of *mKate2* to allow quantification of the promoter activity influenced by AHL-bound LasR. **C.** Crosstalk of the transcription factor EsaR with different acyl-homoserine lactone molecules (AHLs). The promoters P_esaR_ and P_esaS_ regulate the transcription of *mKate2* and *sfGFP*, respectively, to allow quantification of the promoter activity influenced by AHL-bound EsaR. The gray dashed lines depict the basal promoter activity in the absence of the transcription factor. The response curves of the TF with its cognate AI are indicated with an asterisk. Fluorescent values are normalized for cell growth determined by optical density at 600 nm (OD600) and obtained during the stationary growth phase. Error bars represent the standard error for three biological replicates. Pconst = constitutive promoter, RBS = ribosome binding site. An overview of the statistical analysis is given in Supplementary Table S9-17.

Figure 10B shows that the ligand binding of LasR is not limited to 3OC12-HSL. A similar response curve can be observed for 3OC10-HSL with maximal induction around 10 nM. The decrease in fluorescence for the highest concentrations of 3OC10– and 3OC12-HSL can be attributed to the hampered growth, possibly caused by a large quantity of activated LasR in the cytoplasm. A further decrease in acyl chain length of the AI required higher concentrations to activate the P_lasI_ promoter. Nevertheless, at 10 µM, signal crosstalk could be observed for all tested AHLs. Full statistical details can be found in Supplementary Tables S9-11. In literature, comparable results were obtained in which LasR is responsive to high concentrations of 3OC6-HSL [13,14,26]. This implies that the combination of this quorum sensing system with EsaI might lead to low levels of crosstalk caused by the production of 3OC6-HSL by EsaI.

Similarly, the EsaR signal crosstalk was assessed by measuring P_esaR_ and P_esaS_ activity in the presence of different AHL molecules (Figure 10C). A similar response was observed for the natural AI, 3OC6-HSL, as for 3OC8-HSL for both promoters. However, for longer acyl chain lengths of the AHL molecules, the effect on EsaR-binding decreased. With the addition of 3OC10-HSL, a change in promoter activity can only be observed at high concentrations (1 and 10 µM). In the presence of the highest concentration of 3OC12-HSL, the effect is close to promoter activity in the absence of any inducer. Nevertheless, a significant influence can be found of 3OC12-HSL binding to EsaR on the P_esaS_ promoter activity (p = 0.009712). Full statistical details are found in Supplementary Tables S12-17.

#### Synthase crosstalk

To determine synthase crosstalk between the two systems, the transcription factor and promoter of one QS system were combined with the synthase of the second system (Figure 11A). However, the results should be interpreted with care, since it is not possible to distinguish synthase crosstalk from signal crosstalk from this experiment alone. To achieve this, the data should be combined with the results obtained from the signal crosstalk assessment (Figure 10).

**Figure 11.**
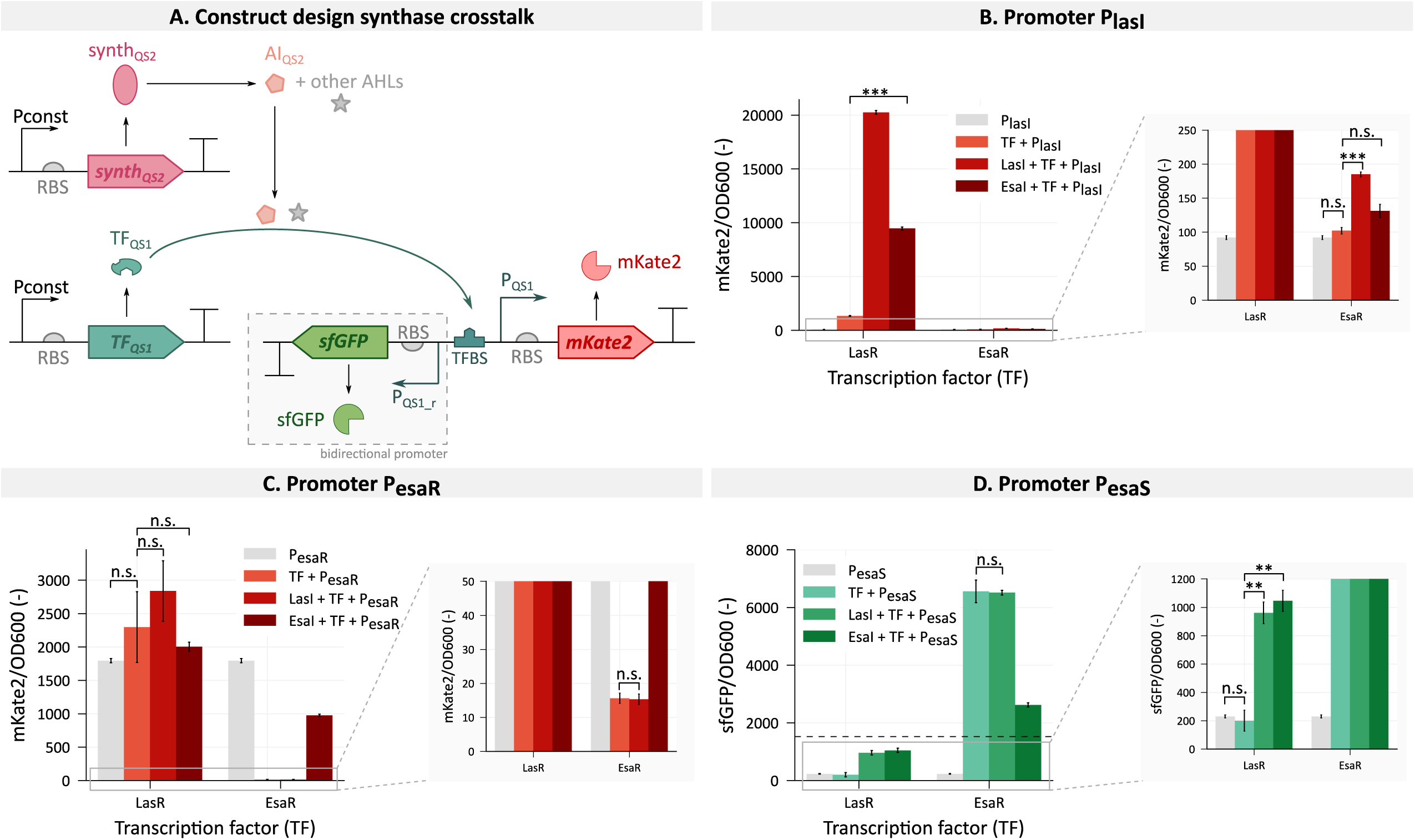
Assessment of the synthase crosstalk between the two quorum sensing (QS) systems. **A.** Construct design for checking synthase crosstalk. A constitutively expressed transcription factor (TF_QS1_) is combined with its own quorum sensing-regulated promoter (P_QS1_), which can be bidirectional (P_QS1_r_). The synthase from the second quorum sensing system (synth_QS2_) is constitutively expressed and the acyl-homoserine lactones (AHLs) produced by it might bind and influence the activity of the transcription factor (TF). Transcription factor activity will directly influence the activity of the quorum sensing-regulated promoter, controlling the expression of a fluorescent reporter protein. Genetic parts are depicted according to SBOL conventions [29,30]. **B.** Promoter activity of P_lasI_, controlling mKate2 expression, influenced by the different combinations of the transcription factors, LasR and EsaR, and the synthases, LasI and EsaI. **C.** Promoter activity of P_esaR_, controlling mKate2 expression, influenced by the different combinations of the transcription factors LasR and EsaR and the synthases LasI and EsaI. **D.** Promoter activity of P_esaS_, controlling sfGFP expression, influenced by the different combinations of the transcription factors, LasR and EsaR, and the synthases, LasI and EsaI. The black dashed line depicts the background fluorescence of the growth medium. Fluorescent values are normalized for cell growth determined by optical density at 600 nm (OD600). For mKate2 measurement, datapoints were chosen during the stationary growth phase. For sfGFP, the maximal production was visualized, since the highest P_esaS_ activity is reached during the exponential phase. Error bars represent the standard error for three biological replicates. The synthase crosstalk was analyzed using a Welsch two-sample t-test. The corresponding outcome is given with p-values * < 0.05, ** < 0.01, *** < 0.001 and n.s. > 0.05. An overview of the statistical analysis is given in Supplementary Table S18.

As was shown earlier in Figure 10B, LasR showed a low response to 10 µM 3OC6-HSL. However, it also responded to lower concentrations of AHLs with longer acyl chains. Clear crosstalk between LasR and EsaI can be observed in Figure 11B, with a large increase in P_lasI_ activity. Hence, it can be concluded that EsaI is not fully specific and produces other AHLs with longer acyl chains besides 3OC6-HSL. Gould *et al.* (2006) analyzed the AHLs produced by EsaI and besides 3OC6– and C6-HSL, low levels of C8– and 3OC8-HSL could be found [49]. Additionally, it was reported that low concentrations of 3OC12-HSL were produced by EsaI [50]. Combined with the signal crosstalk of LasR with these AHL molecules, the EsaI-LasR-P_lasI_ system displays strong crosstalk. This was also observed by Tekel *et al.* (2019) and Jiang *et al.* (2020) [13,15]. From Figure 11C and D, we can conclude that EsaR does not respond to AHLs produced by LasI. This indicates that LasI does not produce 3OC6– and 3OC8-HSL, since EsaR is highly responsive to those AHLs (Figure 10C). Additionally, inhibition of the signals produced by LasI on the 3OC6-HSL recognition by EsaR could not be observed (Supplementary Figure S9A).

Besides testing the synthase crosstalk, all different combinations of synthase, transcription factor and promoter of the two different systems were created to verify if there are no unexpected forms of crosstalk, such as promoter crosstalk, when autoinduced. In contrast to what was concluded earlier, AHL-bound EsaR is able to significantly increase P_lasI_ activity. However, this increase in activity is negligible compared to the range of induction by LasR. Even in the absence of its cognate synthase, inactive LasR is able to activate P_lasI_ almost 15-fold, while AHL-bound EsaR only increases the promoter activity with 29.8% (Figure 11B). Interestingly, given the location of the *las*-box within the promoter site, it would be expected of EsaR to function as an activator rather than a repressor [36]. Nevertheless, the increase in P_lasI_ activity is observed in the presence of EsaI, when EsaR theoretically releases the promoter. This would indicate that EsaR would still function as a repressor. However, care must be taken when interpreting these results, given the slight increase in activity and the absence of this effect during the promoter crosstalk assessment.

The earlier observed promoter crosstalk between LasR and P_esaR_ (Figure 9B) could not be confirmed by this experiment (Figure 11C). Possible explanations are the low level of crosstalk that cannot always be distinguished because of biological variability. Increased P_esaS_ activity could be observed in the presence of AHL-bound LasR. However, the obtained maximal sfGFP values are lower than the background fluorescence of the growth medium. Therefore, we cannot attribute this increase to promoter crosstalk based on these data.

#### Modifications for improved orthogonality: LasR mutant

While assessing the orthogonality of LasR with the 3OC6-HSL inducer of the EsaI/EsaR system, crosstalk could be observed. Not only did LasR respond to the highest tested 3OC6-HSL concentration, it also became activated by 3OC8-HSL at lower concentrations. Furthermore, the high activation of P_lasI_ by LasR combined with EsaI, indicated that EsaI also produces other inducers besides 3OC6-HSL. Therefore, to guarantee orthogonality, a new mutant of the LasR protein was sought with increased specificity for its cognate autoinducer, 3OC12-HSL.

From literature, three mutants of LasR were selected for further orthogonality assessment [15,26,51]. These three mutants, consisting of changes in one amino acid each, could increase the signal specificity of LasR. The first tested mutation, exchanging the proline at position 117 by a serine (P117S), is located at the N-terminal ligand-binding domain, but is not involved in 3OC12-HSL binding. It was hypothesized by Jiang *et al.* (2020) that this mutation hinders multimerization of the DNA-binding domain, thereby increasing the required inducer concentration for dimerization [15]. The second mutant, S129N, replaces the serine at position 129 by an asparagine. This serine is known to be responsible for the hydrogen bond with 3OC12-HSL [26,51]. Lastly, replacing the threonine at position 222 by an isoleucine (T222I) was shown to increase specificity of LasR and is located in the DNA-binding domain of LasR [15]. These three mutants, together with P_lasI_ regulating *mKate2* transcription, were combined with EsaI for assessing orthogonality or with LasI to check for retained functionality (Figure 12A). Although all three mutations successfully led to reduced crosstalk with EsaI, the affinity toward its cognate inducer was also impacted leading to reduced P_lasI_ activation when combined with LasI for LasR(P117S) (p = 0.00017) and LasR(T222I) (p = 8.2*1e-8). Nevertheless, in the strain containing LasR(P117S), crosstalk was reduced more than tenfold, with only a 25.3% decrease in mKate2-expression for the LasI-LasR(P117S)-P_lasI_ strain, leading to the highest fold change between EsaI and LasI induction of this mutant. Therefore, this LasR mutant is the most successful in reducing crosstalk while retaining functionality. Next, LasR(P117S) was combined with different inducers to verify its signal specificity (Figure 12B). LasR(P117S) does not respond to any of the tested 3OC6-HSL concentrations. Furthermore, P_lasI_ activity was only observed for the highest tested 3OC8-HSL concentration, namely 10 µM. Additionally, the affinity of LasR(P117S) toward 3OC10– and 3OC12-HSL was clearly retained, although higher inducer concentrations were required to have the same response compared to the wild type LasR (Figure 10B). Finally, this new mutant was shown not to experience inhibitory effects of the inducers produced by EsaI for binding to its cognate AI, 3OC12-HSL (Supplementary Figure S9B).

**Figure 12.**
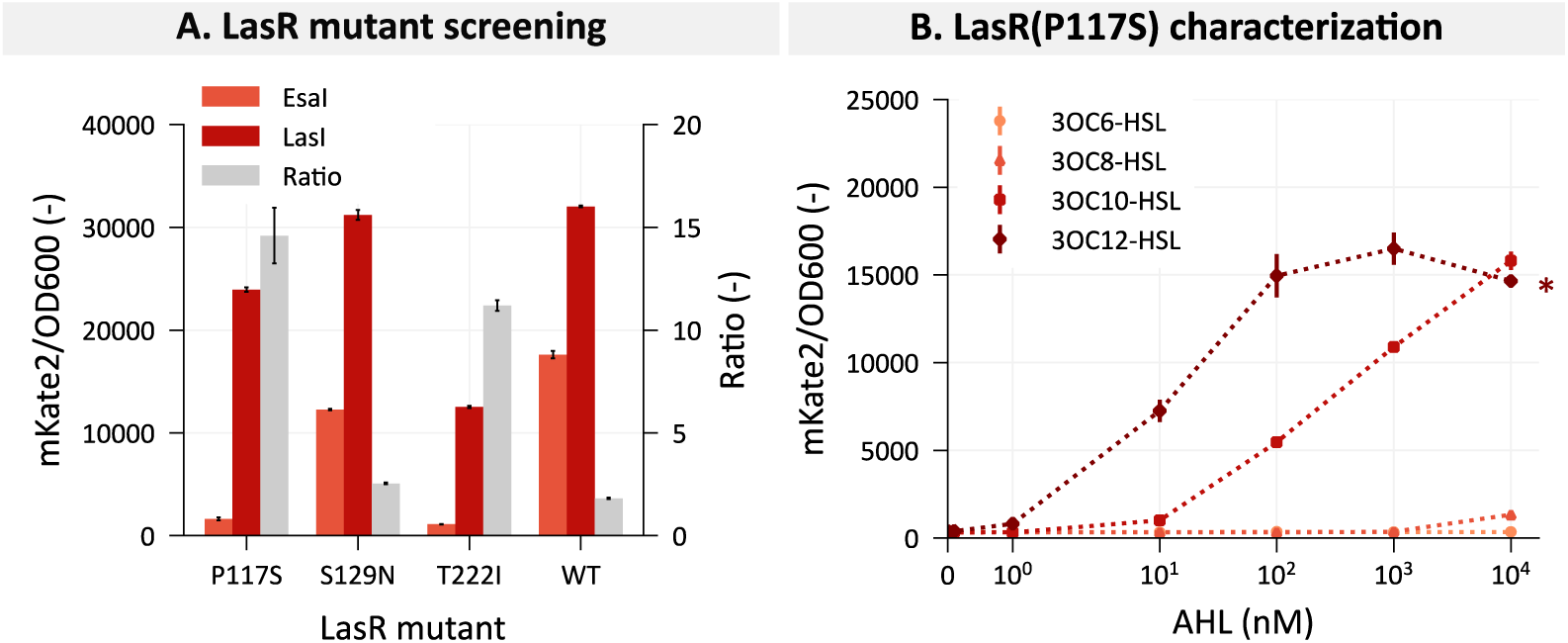
Screening for a LasR mutant with decreased signal crosstalk. **A.** Three different mutant versions of LasR were tested for functionality and orthogonality by combining them with the synthases LasI and EsaI, respectively. The response was measured by following the P_lasI_ activity, expressing mKate2. The wild type LasR (WT) was taken for comparison. **B.** Signal crosstalk of the transcription factor LasR(P117S) with different acyl-homoserine lactone molecules (AHLs). The promoter P_lasI_ regulates the transcription of *mKate2*, to allow quantification of its promoter activity influenced by AHL-bound LasR(P117S). Fluorescent values are normalized for cell growth determined by optical density at 600 nm (OD600). Bars represent the mean and error bars the standard error of the mean based on three biological replicates.

#### Modifications for improved orthogonality: P_esaR/esaS_ mutant

As discussed earlier, low promoter crosstalk could be observed between LasR and both P_esaR_ and P_esaS_. This can be explained by the high similarity in the palindromic transcription factor binding sites of different members of the LuxR-family [23,44,45]. To reduce this crosstalk, a new *esa*-box could be created which retains affinity toward EsaR but is no longer bound by LasR. To achieve this, the DNA-sequence of the used *esa*– and *las*-box were compared. Furthermore, the sequence logos, shown in Figure 8B, can be used to identify important nucleotides for TF-recognition. A few nucleotides appear to be highly conserved over multiple *las*-boxes found in nature, namely the cytosine on position 3 (3C), the thymine on position 13 (13T), the guanine on position 14 (14G), the adenine on position 17 (17A) and lastly the guanine on position 18 (18G). Of these possibly important nucleotides, 3C, 17A and 18G can be found in the *esa*-box of the P_esaR/esaS_ promoter. An indication of the importance of these nucleotides for EsaR-binding can be done by looking at the *esa*-box sequence logo. This shows that 3C is also highly conserved over the compared *esa*-boxes. 17A also appears to be conserved, although to a lesser extent. Lastly, on position 18, there is an equal presence of adenine compared to guanine [46]. Therefore, changing this nucleotide into adenine appears to be the most promising strategy to decrease LasR-binding and retain EsaR-affinity. The influence of this nucleotide change on LasR affinity is also found by Li *et al.* (2007), where it was shown to lead to an 80% reduction in promoter activity [52]. It is generally agreed upon that the nucleotides at position 3 to 5 and 16 to 18 play an important role in the interaction with the transcription factor within the LuxR-family [14,23]. Even though 18G appears to be highly conserved in the *lux*-box as well, replacing this with an adenine has been shown to retain functionality while decreasing promoter crosstalk with LasR [14].

The new promoter region with mutation G18A, referred to as P_esaR/esaS_*, was tested for decreased promoter crosstalk (Figure 13). Even though a decrease in LasR-induced P_esaR_* activity compared to P_esaR_, could be observed, no significant difference could be found, probably due to the large biological variance (p = 0.0939). However, a significant increase in P_esaR_* activity with and without active LasR could be observed. Hence, from these results it is difficult to claim whether this mutation resulted in improved promoter orthogonality between LasR and P_esaR_*. From the results shown in Figure 13, it appears as if the mutation made the promoter crosstalk even worse for P_esaS_*. However, these values are very close to the medium background fluorescence and it is hard to assess whether the observed results are due to actual biological differences or can be explained by the technical set-up of the experiment. The promoter orthogonality was also checked with the new LasR(P117S) variant and no significant up-or downregulation by LasR(P117S) of P_esaR_* or P_esaS_*, respectively, was obtained (Supplementary Figure S10).

**Figure 13.**
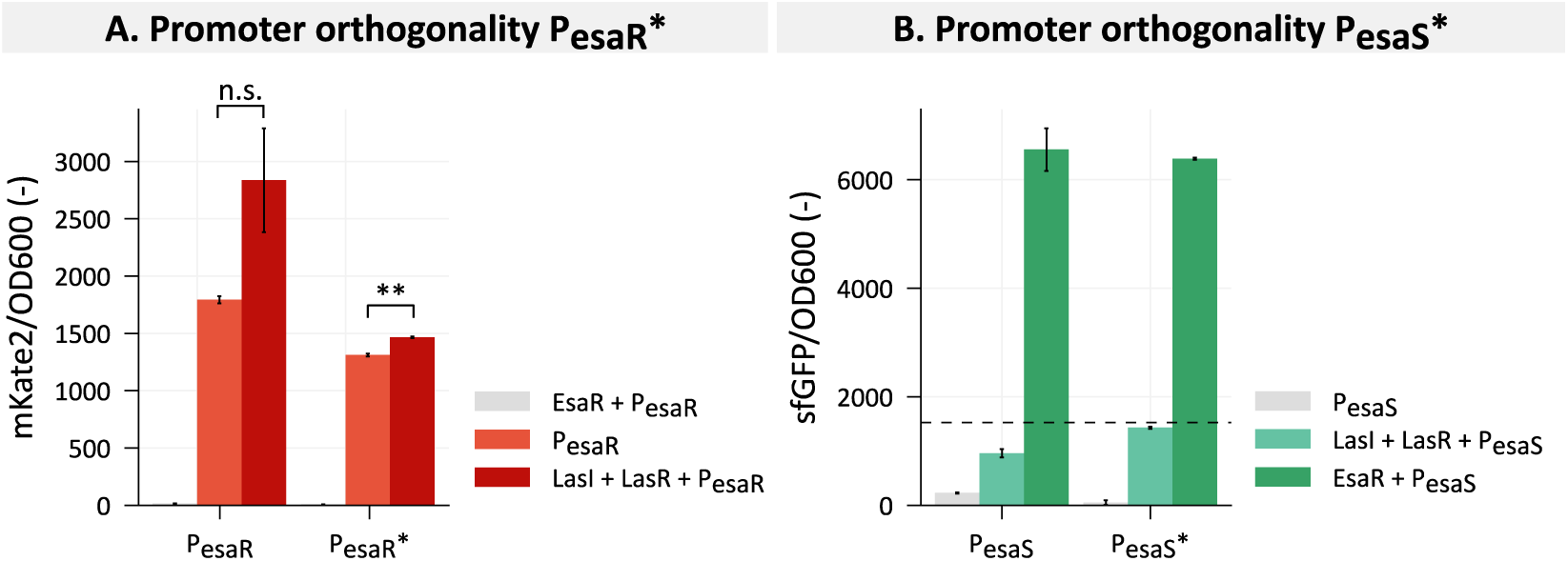
Testing for decreased promoter crosstalk of the mutant promoter P_esaR/esaS_* with the transcription factor LasR, shown for P_esaR_*. (**A.**) and P_esaS_* (**B.**), regulating the expression of the fluorescent proteins mKate2 and sfGFP, respectively. The new promoter was tested for leaky expression and combined with its cognate transcription factor EsaR for reference. The black dashed line depicts the background fluorescence of the growth medium. Fluorescent values are normalized for cell growth determined by optical density at 600 nm (OD600). Bars represent the mean and error bars the standard error of the mean based on three biological replicates. The promoter crosstalk between LasR and P_esaR_ and P_esaR_* was analyzed using a Welsch two-sample t-test. The corresponding outcome is given with p-values * < 0.05, ** < 0.01, *** < 0.001 and n.s. > 0.05. An overview of the statistical analysis is given in Supplementary Table S19.

## 4. Conclusions

To fully unlock the potential of QS systems for synthetic biology applications, thorough characterization is needed to minimize optimization times. As such, we have fully characterized two interesting QS systems on multiple levels. The first chosen QS system, namely the LasI/LasR system, had drawn a lot of attention from biologists for its importance in the virulence of *Pseudomonas aeruginosa*. Therefore, this characterization might be beneficial both for applications and fundamental biological research. The second QS system, the EsaI/EsaR system, has an interesting mechanism of action, because of its bidirectional promoter with opposite regulation.

To conduct this in-depth characterization, we have first excluded the synthase protein from the characterization and replaced it with the extracellular addition of the autoinducer of each specific system. For both QS systems, response curves were obtained. Additionally, it could be concluded that the expression level of the transcription factor influenced the shape of the response curve and the sensitivity of the system for its autoinducers. When the respective synthases were included in the systems, autoinduction could be observed for the LasI/LasR quorum sensing system. Further tuning of the EsaI expression level was required to obtain the desired autoinduction for the EsaI/EsaR system. This highlights the importance of the tight balance between the synthase and transcription factor level in this dynamic system.

Next, we assessed the full orthogonality between the two QS systems. This ensures that the two systems are functioning independently inside of the same cell or population without interfering with each other. The orthogonality was checked on all three levels: promoter, signal and synthase crosstalk. Promoter crosstalk could be observed of LasR with P_esaR/esaS_. By mutating one nucleotide in the *esa*-box situated within this promoter region, reduced affinity of LasR for this sequence was expected. However, the results are inconclusive whether a more orthogonal promoter was obtained. Additionally, LasR could also respond to the AHLs produced by EsaI by a combination of signal and synthase crosstalk. This was resolved with a mutant version of LasR, namely LasR(P117S), which no longer responded to EsaI-induction. With this improvement in the orthogonality, we assume that the two systems can now be simultaneously applied. Even with the possible low levels of promoter crosstalk between LasR and P_esaR/esaS_ remaining, it is assumed that in the presence of P_lasI_, this transcription factor will preferably bind to its own cognate promoter. Similarly, the presence of EsaR would likely outcompete the affinity of LasR for the P_esaR/esaS_ promoter region. Altogether, we succeeded in characterizing two orthogonal quorum sensing systems with great potential for synthetic biology applications. Furthermore, the created workflow can be applied to different quorum sensing systems to further expand the toolbox of characterized AHL-based quorum sensing systems.

## 5. Author contributions

J. De Baets designed and performed the experiments. J. De Baets analyzed the data and wrote the manuscript. B. De Paepe and M. De Mey were involved in the conceptualization and design of this work and in correcting and revising the manuscript.

## 8. Supplementary data

Supplementary File 1 contains:

– Supplementary Figures S1-10
– Supplementary Tables S1-19

Supplementary File 2 contains the ID-sheets of LasR and EsaR biosensors with different ribosome binding site sequences.

