## Supplementary File 1 for "Characterization and orthogonality assessment of two quorum sensing systems for synthetic biology applications"

### Characterization and orthogonality assessment of two quorum sensing systems for synthetic biology applications

Jasmine De Baets, Brecht De Paepe, Marjan De Mey

*Centre for Synthetic Biology, Ghent university, 9000 Ghent, Belgium*

**Overview**

**Supplementary Figure S1:** Quantification of the LasR and EsaR transcription factors for different RBS-strength and at different autoinducer concentrations.
**Supplementary Figure S2:** Confocal light scanning microscopy images of LasR in the absence and presence of its autoinducer.
**Supplementary Figure S3:** Strain from the RBS-library controlling EsaI expression with too low EsaI expression.
**Supplementary Figure S4:** Overview of the strains from the RBS-library controlling EsaI expression, resulting in the desired response.
**Supplementary Figure S5:** Overview of the first part of the strains from the RBS-library controlling EsaI expression with too high EsaI expression.
**Supplementary Figure S6:** Overview of the second part of the strains from the RBS-library controlling EsaI expression with too high EsaI expression.
**Supplementary Figure S7:** Overview of the third part of the strains from the RBS-library controlling EsaI expression with too high EsaI expression.
**Supplementary Figure S8:** Overview of the fourth part of the strains from the RBS-library controlling EsaI expression with too high EsaI expression.
**Supplementary Figure S9:** Assessment of the signal inhibition between the EsaI/EsaR and LasI/LasR quorum sensing systems.
**Supplementary Figure S10:** Assessment of the orthogonality of the LasR(P117S) variant with the new $P_{esaR/esaS}*$ mutant.

---

**Supplementary Table S1:** Overview of the plasmids used in this research.
**Supplementary Table S2:** Overview of the DNA-sequence of all regulatory parts used in this research.
**Supplementary Table S3:** Overview of the genes used in this research.
**Supplementary Table S4:** Overview of the primers used in this research.
**Supplementary Table S5:** Sequence of the RBS-library controlling *esaI* translation.
**Supplementary Table S6:** Statistical tests for the binding of LasR to $P_{esaR}$.
**Supplementary Table S7:** Statistical tests for the binding of LasR to $P_{esaS}$.
**Supplementary Table S8:** Statistical tests for the binding of EsaR to $P_{lasI}$.
**Supplementary Table S9:** Statistical tests for the binding of LasR to 3OC6-HSL.
**Supplementary Table S10:** Statistical tests for the binding of LasR to 3OC8-HSL.
**Supplementary Table S11:** Statistical tests for the binding of LasR to 3OC10-HSL.
**Supplementary Table S12:** Statistical tests for the influence of 3OC8-HSL addition on EsaR repression of $P_{esaR}$.

**Supplementary Table S13:** Statistical tests for the influence of 3OC10-HSL addition on EsaR repression of $P_{esaR}$.
**Supplementary Table S14:** Statistical tests for the influence of 3OC12-HSL addition on EsaR repression of $P_{esaR}$.
**Supplementary Table S15:** Statistical tests for the influence of 3OC8-HSL addition on EsaR activation of $P_{esaS}$.
**Supplementary Table S16:** Statistical tests for the influence of 3OC10-HSL addition on EsaR activation of $P_{esaS}$.
**Supplementary Table S17:** Statistical tests for the influence of 3OC12-HSL addition on EsaR activation of $P_{esaS}$.
**Supplementary Table S18:** Statistical tests for the synthase crosstalk.
**Supplementary Table S19:** Statistical tests for the orthogonality of LasR with the new $P_{esaR/esaS}*$ promoter.

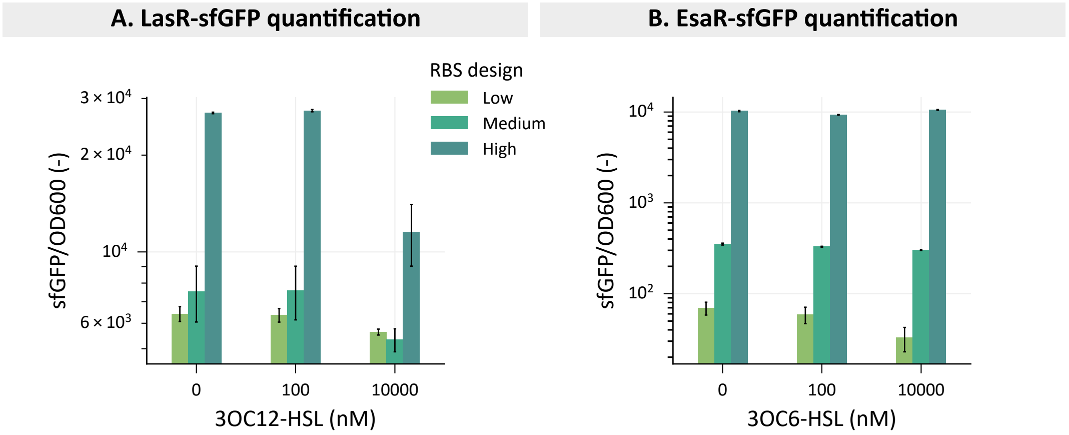


**Figure S1:** Quantification of the expression level of transcription factor-sfGFP fusion, influenced by three different ribosome binding site (RBS) sequences for both LasR (**A.**) and EsaR (**B.**) at different concentrations of 3-oxo-dodecanoyl homoserine lactone (3OC12-HSL) and 3-oxo-hexanoyl homoserine lactone (3OC6-HSL), respectively. Fluorescent values were obtained in the stationary phase and normalized for cell growth determined by optical density at 600 nm (OD600). Error bars represent the standard error for 3 biological replicates.

#### A. Positive control: sfGFP in cytoplasm

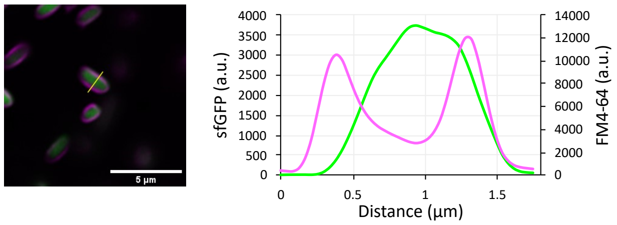


#### B. Positive control: sfGFP targeted to membrane

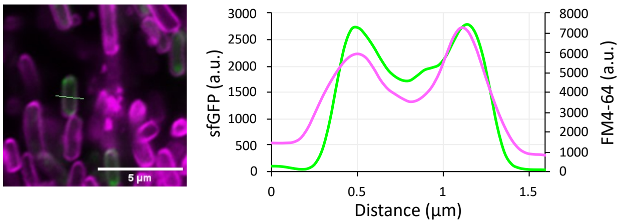


#### C. LasR-$P_{lasI}$ + 10 µM 3OC12-HSL

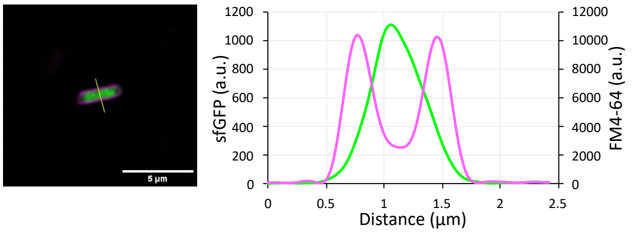


#### D. LasR-$P_{lasI}$ + 10 nM 3OC12-HSL

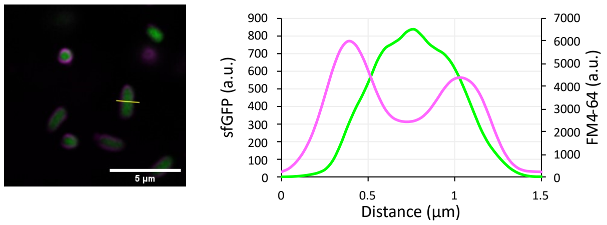


#### E. LasR-$P_{lasI}$ + 0 nM 3OC12-HSL

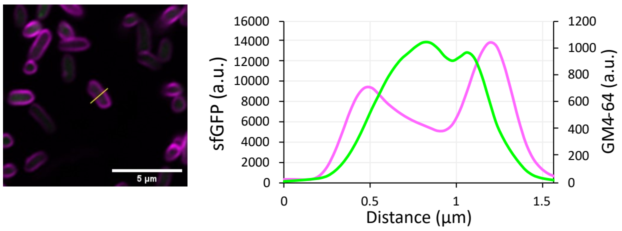


**Figure S2:** Confocal light scanning microscopy images for the LasR-sfGFP fusion at different 3-oxo-dodecanoyl homoserine lactone (3OC12-HSL) concentrations with intensity cross-section profiles for both the red FM4-64 membrane dye (purple) and sfGFP (green). The cross-sections analyzed are indicated in yellow in the images. **A.** and **B.** represent positive controls for cytoplasmic and membrane-targeted sfGFP expression, respectively.

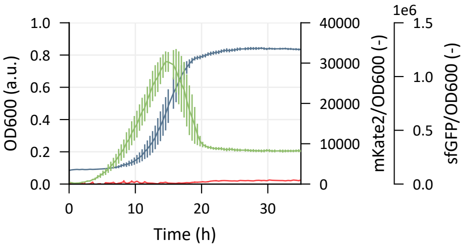


**Figure S3:** Strain from group 1 of the RBS-library for tuning the expression of EsaI in the EsaR/EsaI quorum sensing strain. This strain retained maximal sfGFP expression, while only producing leaky mKate2. Fluorescent values are normalized for cell growth determined by optical density at 600 nm (OD600). Error bars represent the standard error for 3 biological replicates.

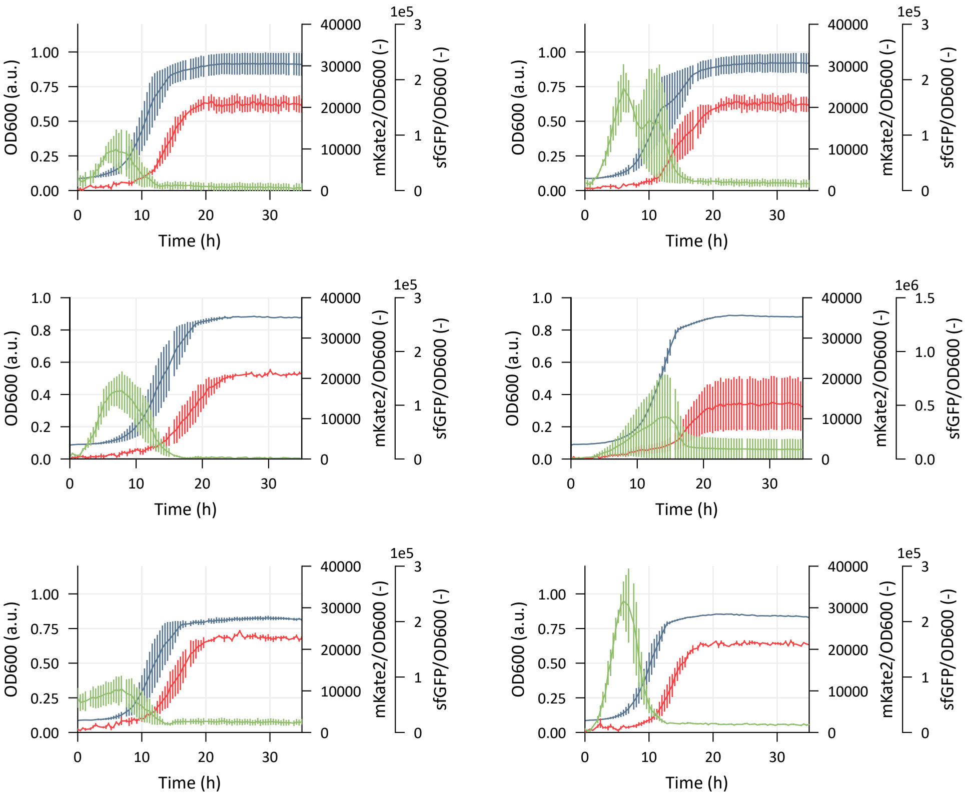


**Figure S4:** Strains from group 3 of the RBS-library for tuning the expression of EsaI in the EsaR/EsaI quorum sensing strain. In strains from this group, mKate2 production follows sfGFP production, with a switch during the early exponential phase. Fluorescent values are normalized for cell growth determined by optical density at 600 nm (OD600). Error bars represent the standard error for 3 biological replicates.

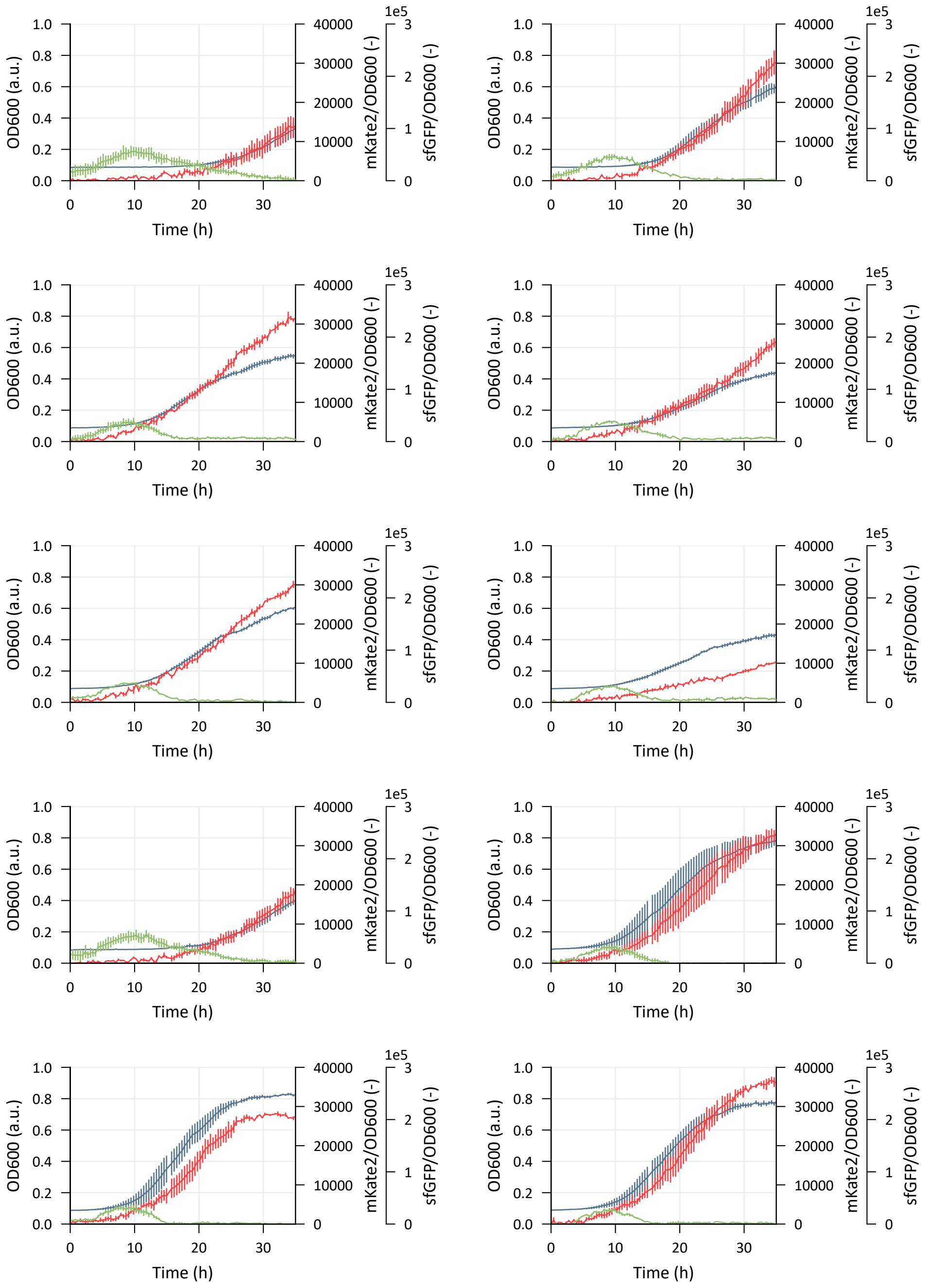


**Figure S5:** Strains from group 2 of the RBS-library for tuning the expression of EsaI in the EsaR/EsaI quorum sensing strain. In strains from this group, the mKate2 reaches maximal values but no sfGFP can be observed. Fluorescent values are normalized for cell growth determined by optical density at 600 nm (OD600). Error bars represent the standard error for 3 biological replicates.

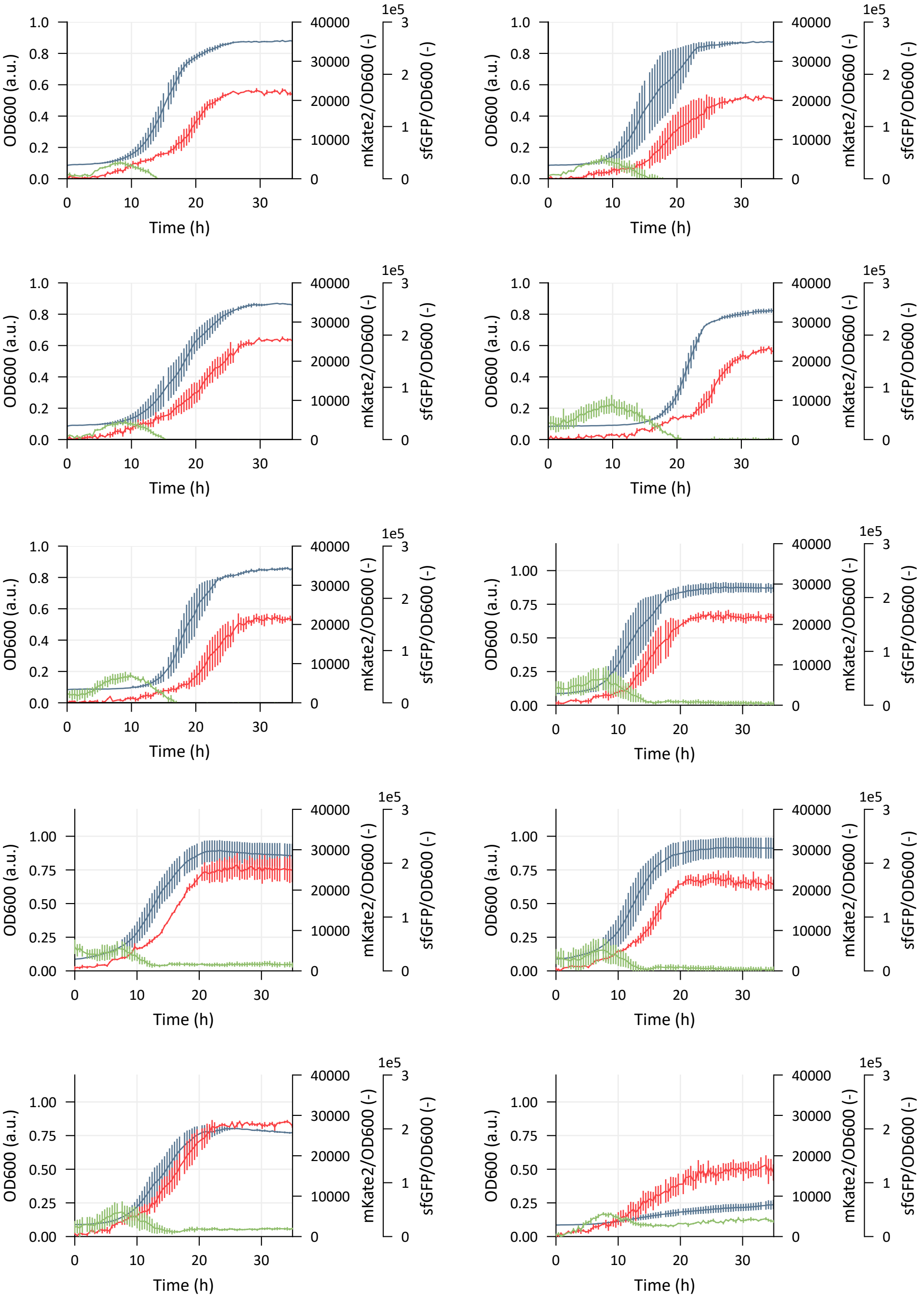


**Figure S6:** Continued from Figure S5. Strains from group 2 of the RBS-library for tuning the expression of EsaI in the EsaR/EsaI quorum sensing strain. In strains from this group, the mKate2 reaches maximal values but no sfGFP can be observed. Fluorescent values are normalized for cell growth determined by optical density at 600 nm (OD600). Error bars represent the standard error for 3 biological replicates.

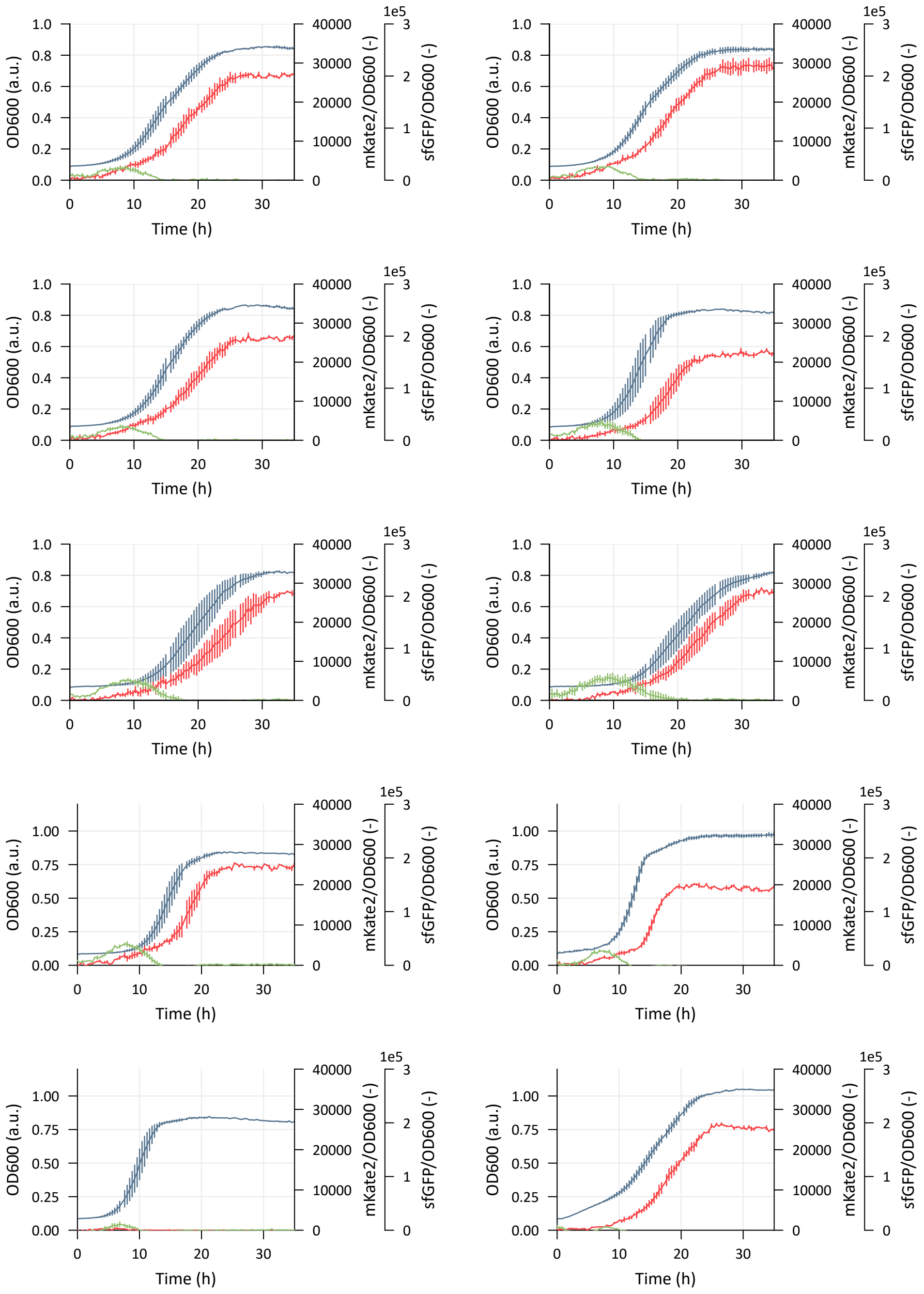


**Figure S7:** Continued from Figure S6. Strains from group 2 of the RBS-library for tuning the expression of EsaI in the EsaR/EsaI quorum sensing strain. In strains from this group, the mKate2 reaches maximal values but no sfGFP can be observed. Fluorescent values are normalized for cell growth determined by optical density at 600 nm (OD600). Error bars represent the standard error for 3 biological replicates.

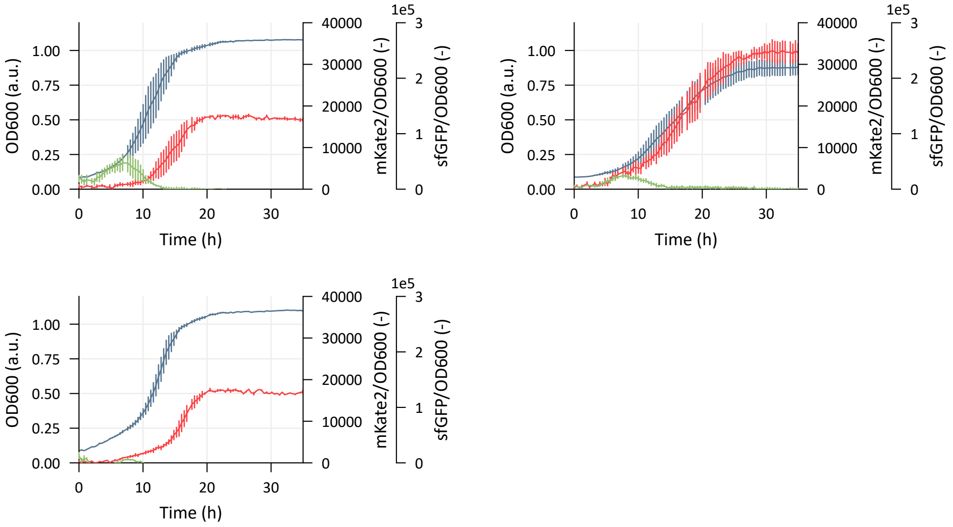


**Figure S8:** Continued from Figure S7. Strains from group 2 of the RBS-library for tuning the expression of EsaI in the EsaR/EsaI quorum sensing strain. In strains from this group, the mKate2 reaches maximal values but no sfGFP can be observed. Fluorescent values are normalized for cell growth determined by optical density at 600 nm (OD600). Error bars represent the standard error for 3 biological replicates.

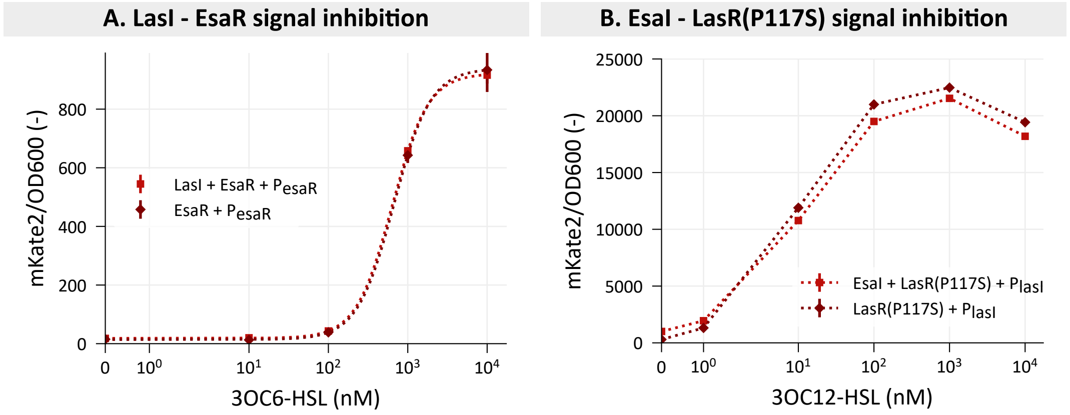


**Figure S9:** Assessment of the signal inhibition between the EsaI/EsaR and LasI/LasR quorum sensing systems. **A.** The response curve of the EsaR-$P_{esaR}$ system with 3-oxo-hexanoyl homoserine lactone (3OC6-HSL) in the presence and the absence of LasI. **B.** The response curve of the LasR(P117S)-$P_{lasI}$ system with 3-oxo-dodecanoyl homoserine lactone (3OC12-HSL) in the presence and the absence of EsaI. Fluorescent values are normalized for cell growth determined by optical density at 600 nm (OD600). Error bars represent the standard error for 3 biological replicates.

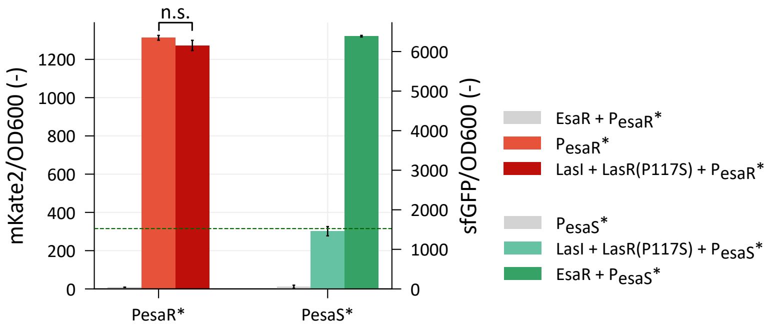


**Figure S10:** Testing for decreased promoter crosstalk of the mutant promoter $P_{esaR/esaS}*$ with the transcription factor LasR(P117S), shown for $P_{esaR}*$ (**A.**) and $P_{esaS}*$ (**B.**), regulating the fluorescent proteins mKate2 and sfGFP expression, respectively. The new promoter was tested for leaky expression and combined with its cognate transcription factor EsaR for reference. The green dashed line depicts the background fluorescence of the growth medium. Fluorescent values are normalized for cell growth determined by optical density at 600 nm (OD600). Bars represent the mean and error bars the standard error of the mean for 3 biological replicates. The promoter crosstalk between LasR and $P_{esaR}$ and $P_{esaR}*$ was analyzed using a Welsch two-sample t-test. The corresponding outcome is given with p-values * < 0.05, ** < 0.01, *** < 0.001 and n.s. > 0.05. An overview of the statistical analysis is given in Supplementary Table S19.

**Table S1:** Overview of the plasmids used this research. All plasmids are made in the MoBioS platform [11].

| Plasmid | Plasmid details: p[Ori][Marker]-[insert 1: promoter - RBS - gene - terminator]-[insert 2] |
|---|---|
| | **Response curves** |
| LasR(low)-Plas | p[BBR1-MCS2][Kan]-[MoBioS-M4][P22-RBS[TIRlow]-LasR-TT3_rrnD1_T1]-[PlasI-Bba_B0030-mKate2-FAB391’] |
| LasR(med)-Plas | p[BBR1-MCS2][Kan]-[MoBioS-M4][P22-RBS[TIRmedium]-LasR-TT3_rrnD1_T1]-[PlasI-Bba_B0030-mKate2-FAB391’] |
| LasR(high)-Plas | p[BBR1-MCS2][Kan]-[MoBioS-M4][P22-RBS[TIRhigh]-LasR-TT3_rrnD1_T1]-[PlasI-Bba_B0030-mKate2-FAB391’] |
| EsaR(low)-Pesa | p[BBR1-MCS2][Kan]-[MoBioS-M4][P22-RBS[TIRlow]-EsaR-TT3_rrnD1_T1]-[TT3_rrnB_T1’-sfGFP-B0032-PesaR/S-B0030-mKate2-FAB391’] |
| EsaR(med)-Pesa | p[BBR1-MCS2][Kan]-[MoBioS-M4][P22-RBS[TIRmedium]-EsaR-TT3_rrnD1_T1]-[TT3_rrnB_T1’-sfGFP-B0032-PesaR/S-B0030-mKate2-FAB391’] |
| EsaR(high)-Pesa | p[BBR1-MCS2][Kan]-[MoBioS-M4][P22-RBS[TIRhigh]-EsaR-TT3_rrnD1_T1]-[TT3_rrnB_T1’-sfGFP-B0032-PesaR/S-B0030-mKate2-FAB391’] |
| **sfGFP fusions** | |
| LasR(low)_sfGFP | p[BBR1-MCS2][Kan]-[MoBioS-M4][P22-RBS[TIRlow]-LasR-linker_sfGFP-TT3_rrnD1_T1]-[Pjunk-mKate2-FAB391’] |
| LasR(med)_sfGFP | p[BBR1-MCS2][Kan]-[MoBioS-M4][P22-RBS[TIRmedium]-LasR-linker_sfGFP-TT3_rrnD1_T1]-[Pjunk-mKate2-FAB391’] |
| LasR(high)_sfGFP | p[BBR1-MCS2][Kan]-[MoBioS-M4][P22-RBS[TIRhigh]-LasR-linker_sfGFP-TT3_rrnD1_T1]-[Pjunk-mKate2-FAB391’] |
| EsaR(low)_sfGFP | p[BBR1-MCS2][Kan]-[MoBioS-M4][P22-RBS[TIRlow]-EsaR-linker_sfGFP-TT3_rrnD1_T1]-[Pjunk-mKate2-FAB391’] |
| EsaR(med)_sfGFP | p[BBR1-MCS2][Kan]-[MoBioS-M4][P22-RBS[TIRmedium]-EsaR-linker_sfGFP-TT3_rrnD1_T1]-[Pjunk-mKate2-FAB391’] |
| EsaR(high)_sfGFP | p[BBR1-MCS2][Kan]-[MoBioS-M4][P22-RBS[TIRhigh]-EsaR-linker_sfGFP-TT3_rrnD1_T1]-[Pjunk-mKate2-FAB391’] |
| **Synthases** | |
| LasQS | p[BBR1-MCS2][Kan]-[MoBioS-M4][Bba_J23108-Bba_B0031-LasI-TT9_RNA_IT]-[P22-RBS[TIRlow]-LasR-TT3_rrnD1_T1]-[PlasI-Bba_B0030-mKate2-FAB391’] |

**Table S1:** Continued overview of plasmids used in this research.

| Plasmid | Plasmid details: p[Ori][Marker]-[insert 1: promoter - RBS - gene - terminator]-[insert 2] |
|---|---|
| EsaQS | p[BBR1-MCS2][Kan]-[MoBioS-M4][Bba_J23108-Bba_B0031-EsaI-TT8-T3TE]-[P22-RBS[TIRlow]-EsaR-TT3_rrnD1_T1]-[TT3_rrnB_T1'-sfGFP-B0032-PesaR/S-B0030-mKate2-FAB391'] |
| **EsaI-library** | |
| EsaI-lib_EsaR_Pesa(LVA) | p[BBR1-MCS2][Kan]-[MoBioS-M4][Bba_J23108-RBS(lib)-EsaI-TT8-T3TE]-[P22-RBS[TIRmedium]-EsaR-TT3_rrnD1_T1]-[TT3_rrnB_T1'-sfGFP(LVA)-B0032-PesaR/S-B0030-mKate2-FAB391'] |
| **Orthogonality** | |
| LasR_Pesa | p[BBR1-MCS2][Kan]-[MoBioS-M4][P22-RBS[TIRlow]-LasR-TT3_rrnD1_T1]-[TT3_rrnB_T1'-sfGFP-B0032-PesaR/S-B0030-mKate2-FAB391'] |
| EsaR_Plas | p[BBR1-MCS2][Kan]-[MoBioS-M4][P22-RBS[TIRmedium]-EsaR-TT3_rrnD1_T1]-[PlasI-Bba_B0030-mKate2-FAB391'] |
| EsaI_LasR_Plas | p[BBR1-MCS2][Kan]-[MoBioS-M4][Bba_J23108-RBS45-EsaI-TT8-T3TE]-[P22-RBS[TIRlow]-LasR-TT3_rrnD1_T1]-[PlasI-Bba_B0030-mKate2-FAB391'] |
| LasI_EsaR_Pesa(LVA) | p[BBR1-MCS2][Kan]-[MoBioS-M4][Bba_J23108-Bba_B0031-LasI-TT9_RNA_IT]-[P22-RBS[TIRmedium]-EsaR-TT3_rrnD1_T1]-[TT3_rrnB_T1'-sfGFP(LVA)-B0032-PesaR/S-B0030-mKate2-FAB391'] |
| LasI_LasR_Pesa(LVA) | p[BBR1-MCS2][Kan]-[MoBioS-M4][Bba_J23108-Bba_B0031-LasI-TT9_RNA_IT]-[P22-RBS[TIRlow]-LasR-TT3_rrnD1_T1]-[TT3_rrnB_T1'-sfGFP(LVA)-B0032-PesaR/S-B0030-mKate2-FAB391'] |
| EsaI_EsaR_Plas | p[BBR1-MCS2][Kan]-[MoBioS-M4][Bba_J23108-RBS45-EsaI-TT8-T3TE]-[P22-RBS[TIRmedium]-EsaR-TT3_rrnD1_T1]-[PlasI-Bba_B0030-mKate2-FAB391'] |
| **LasR-mutant** | |
| EsaI_LasR(S129N)_Plas | p[BBR1-MCS2][Kan]-[MoBioS-M4][Bba_J23108-RBS45_EsaI-TT8-T3TE]-[P22-LasR_S129N[TIRLow]-TT3_rrnD1_T1]-[PlasI-B0030-mKate2-FAB391'] |
| EsaI_LasR(P117S)_Plas | p[BBR1-MCS2][Kan]-[MoBioS-M4][Bba_J23108-RBS45_EsaI-TT8-T3TE]-[P22-LasR_P117S[TIRLow]-TT3_rrnD1_T1]-[PlasI-B0030-mKate2-FAB391'] |
| EsaI_LasR(T222I)_Plas | p[BBR1-MCS2][Kan]-[MoBioS-M4][Bba_J23108-RBS45_EsaI-TT8-T3TE]-[P22-LasR_T222I[TIRLow]-TT3_rrnD1_T1]-[PlasI-B0030-mKate2-FAB391'] |
| LasI_LasR(S129N)_Plas | p[BBR1-MCS2][Kan]-[MoBioS-M4][Bba_J23108-Bba_B0031-LasI-TT9_RNA_IT]-[P22-LasR_S129N[TIRLow]-TT3_rrnD1_T1]-[TFBS[Las]-B0030-mKate2-FAB391'] |
| LasI_LasR(P117S)_Plas | p[BBR1-MCS2][Kan]-[MoBioS-M4][Bba_J23108-Bba_B0031-LasI-TT9_RNA_IT]-[P22-LasR_P117S[TIRLow]-TT3_rrnD1_T1]-[PlasI-B0030-mKate2-FAB391'] |

**Table S1:** Continued overview of plasmids used in this research.

| Plasmid | Plasmid details: p[Ori][Marker]-[insert 1: promoter - RBS - gene - terminator]-[insert 2] |
|---|---|
| LasI_LasR(T222I)_Plas | p[BBR1-MCS2][Kan]-[MoBioS-M4][Bba_J23108-Bba_B0031-LasI-TT9_RNA_IT]-[P22-LasR_T222I[TIRLow]-TT3_rrnD1_T1]-[PlasI-B0030-mKate2-FAB391’] |
| **Pesa-mutant** | |
| EsaR_Pesa-orth(LVA) | p[BBR1-MCS2][Kan]-[MoBioS-M4][p22-EsaR[TIRLow]-TT3_rrnD1_T1]-[TT3_rrnB_T1’-sfGFP(LVA)-B0032-PesaR/S[orth]-B0030-mKate2-FAB391’] |
| Pesa-orth(LVA) | p[BBR1-MCS2][Kan]-[MoBioS-M4][TF_junk]-[TT3_rrnB_T1’-sfGFP(LVA)-B0032-PesaR/S[orth]-B0030-mKate2-FAB391’] |
| LasI_LasR_Pesa-orth(LVA) | p[BBR1-MCS2][Kan]-[MoBioS-M4][Bba_J23108-Bba_B0031-LasI][P22-LasR[TIRLow]-TT3_rrnD1_T1]-[TT3_rrnB_T1’-sfGFP(LVA)-B0032-PesaR/S[orth]-B0030-mKate2-FAB391’] |
| **LasR-mut orthogonality** | |
| LasI_LasR(P117S)_Pesa(LVA) | p[BBR1-MCS2][Kan]-[MoBioS-M4][Bba_J23108-Bba_B0031-LasI-TT9_RNA_IT]-[P22-LasR_P117S[TIRLow]-TT3_rrnD1_T1]-[TT3_rrnB_T1’-sfGFP(LVA)-B0032-PesaR/S-B0030-mKate2-FAB391’] |
| EsaI_LasR(P117S)_Pesa(LVA) | p[BBR1-MCS2][Kan]-[MoBioS-M4][Bba_J23108-RBS45-EsaI-TT8-T3TE][P22-LasR_P117S[TIRLow]-TT3_rrnD1_T1]-[TT3_rrnB_T1’-sfGFP(LVA)-B0032-PesaR/S-B0030-mKate2-FAB391’] |

**Table S2:** Overview of the DNA sequence of all regulatory parts used in this research.

| Part | DNA sequence |
|---|---|
| **Promoters** | |
| P22 | TTGACATTTTGGAATAGATGTGATATAATGTGTACATAT |
| J23108 | CTGACAGCTAGCTCAGTCCTAGGTATAATGCTAGC |
| PesaR/esaS | CCGCTAAACAACTGAAGCCATTGTAACCTCTGAATGATTCATTGTAAGTTACTCTTAAGTATCATCTTGCCTGTACTATAGTGCAGGTTAAGTCCACGTTAAGTAAAAGAAGCAGC |
| PlasI | TTCGAGCCTAGCAAGGGTCCGGGTTCACCGAAATCTATCTCATTTGCTAGTTATAAAATTATGAAATTTGCATAAATTCTTCA |
| PesaR/esaS(orth) | CCGCTAAACAACTGAAGCCATTGTAACCTCTGAATGATTCATTGTAAGTTACTCTTAAGTATCATCTTGCCTGTACTATAGTGCAAGTTAAGTCCACGTTAAGTAAAAGAAGCAGC |
| **RBS** | |
| RBS_low(EsaR) | CGTCACACTACCCGCTAACTGAAGCGGGTCATAGACCGCGTCTCT |
| RBS_medium(EsaR) | CGTCACACTACCCGCTAGCCCCCCACAAATTTAAGATACGTCTCT |
| RBS_high(EsaR) | CGTCACACTACCCGCTGCTACTAACATCTACGAGGAGGCGTCTCT |
| RBS_low(LasR) | CGTCACACTACCCGCTAACAACAACCAAACTAAGGACGCGTCTCT |
| RBS_medium(LasR) | CGTCACACTACCCGCTACGGGTTGAGGAAGACAGGGGTCGTCTCT |
| RBS_high(LasR) | CGTCACACTACCCGCTAGCTGGCGGGGAACTGAAGGAGCGTCTCT |
| Bba_B030 | TCTAGAGATTAAAGAGGAGAAATACTAG |
| Bba_B032 | ATACTAGAGTCACACAGGAAAGTACTAG |
| RBS45 | ACACGATCTTCGAAGGACGTACAT |
| **Terminators** | |
| TT3-rrnD1-T1 | GGGAACTGCCAGACATCAAATAAAACAAAAGGCTCAGTCGGAAGACTGGGCCTTTTGTTTTATCTGTTGTTTGTCGGTGAACACTCTCCC |
| TT8-T3 TE | CCCTCAAGAGAAAATGTAACCAACTCACTGGCTCACCTTCACGGGTGGGCCTTTCTTCGTTCCGGGCATTAACCCTCACTAACAGGAGA |
| TT9-RNAIT | GAGTTGGTAGCTCTTGATCCGGCAAACAAACCACCGTTGGTAGCGGTGGTTTTTTTGTTTGCAAGCAGCAGATTACGCGCAGAAAAAAAGG |
| BioFab terminator_FAB391’ | TCGGTCAGTTTCACCTGATTTACGTAAAAACCCGCTTCGGCGGGTTTTTGCTTTTGGAGGGGCAGAAAGATGAATGACTGTC |

**Table S3:** Overview of the coding sequence of all genes used in this research.

| Part | Coding sequence |
|---|---|
| LasR | 1 ATGGCACTGG TTGATGGTTT TCTGGAACTG GAACGTAGCA GCGGTAAACT GGAATGGTCA GCAATCCTGC<br>71 AGAAAATGGC AAGCGATCTG GGTTTTAGCA AAATTCTGTT TGGTCTGCTG CCGAAAGATA GCCAGGATTA<br>141 TGAAAATGCC TTTATCGTGG GTAATTATCC GGCAGCATGG CGTGAACATT ATGATCGTGC AGGTTATGCA<br>211 CGTGTTGATC CGACCGTTAG CCATTGTACC CAGAGCGTTC TGCCGATTTT TTGGGAACCG AGCATTTATC<br>281 AGACCCGTAA ACAGCACGAA TTTTTTGAAG AAGCAAGCGC AGCAGGTCTG GTTTATGGTC TGACCATGCC<br>351 GCTGCATGGC GCACGTGGTG AACTGGGTGC ACTGAGCCTG AGCGTTGAAG CAGAAAATCG TGCCGAAGCA<br>421 AATCGTTTTA TGGAAAGCGT GCTGCCGACA CTGTGGATGC TGAAAGATTA TGCACTGCAG AGCGGTGCCG<br>491 GTCTGGCATT TGAACATCCG GTTAGCAAAC CGGTGGTTCT GACCAGCCGT GAAAAAGAAG TTCTGCAGTG<br>561 GTGTGCAATT GGTAAAACCA GCTGGGAAAT TAGCGTTATT TGTAATTGTA GCGAAGCCAA CGTGAACTTT<br>631 CATATGGGTA ATATTCGTCG CAAATTTGGT GTTACCAGCC GTCGTGTTGC AGCAATTATG GCAGTTAATC<br>701 TGGGTCTGAT TACCCTGTAA |
| EsaR | 1 ATGTTCTCGT TCTTCCTGGA AAACCAGACC ATTACGGATA CGCTTCAGAC TTACATACAG AGAAAGTTAT<br>71 CTCCGCTGGG TAGTCCGGAT TACGCTTACA CTGTTGTGAG CAAAAAAAAT CCTTCAAATG TTCTGATTAT<br>141 TTCCAGTTAT CCTGACGAAT GGATTAGGTT ATACCGCGCT AACAACTTTC AGCTGACCGA TCCGGTTATT<br>211 CTCACGGCCT TTAAACGCAC CTCGCCGTTT GCCTGGGATG AGAATATTAC GCTGATGTCC GACCTGCGGT<br>281 TCACCAAAAT TTTCTCTTTA TCCAAGCAAT ACAACATCGT TAACGGCTTT ACCTATGTCC TGCATGACCA<br>351 CATGAACAAC CTTGCTCTGT TGTCCGTGAT CATTAAAGGC AACGATCAGA CTGCGCTGGA GCAACGCCTT<br>421 GCTGCCGAAC AGGGCACGAT GCAGATGCTG CTGATTGATT TTAACGAGCA GATGTACCGC CTGGCCGGTA<br>491 CCGAAGGCGA GCGAGCCCCG GCGTTAAATC AGAGCGCGGA CAAAACGATA TTTTCCTCGC GTGAAAATGA<br>561 GGTGTTGTAC TGGGCGAGTA TGGGCAAAAC CTATGCTGAG ATTGCCGCTA TTACGGGCAT TTCTGTGAGT<br>631 ACCGTGAAGT TTCACATCAA GAATGTGGTC GTGAAACTGG GCGTCAGTAA CGCCCGACAG GCTATCAGAC<br>701 TGGGTGTAGA ACTGGATCTT ATCAGACCGG CAGCATCAGC TGCAAGGTAG |

**Table S3:** Continued overview of the coding sequences used in this research.

| Part | Coding sequence |
|---|---|
| LasI | 1 ATGATCGTAC AGATTGGTCG TCGCGAAGAG TTCGATAAAA AACTGCTGGG TGAAATGCAC AAACTGCGTG |
| | 71 CTCAGGTTTT CAAAGAACGT AAAGGTTGGG ACGTTAGCGT CATCGACGAA ATGGAAATCG ATGGTTATGA |
| | 141 CGCACTGAGC CCGTATTACA TGCTGATCCA GGAAGATACT CCGGAAGCCC AGGTTTTCGG TTGCTGGCGT |
| | 211 ATTCTCGATA CCACCGGTCC GTACATGCTG AAAAACACCT TCCCGGAACT GCTGCACGGT AAAGAAGCGC |
| | 281 CTTGCTCGCC GCACATCTGG GAACTGAGCC GTTTCGCCAT CAACTCTGGT CAGAAAGGTT CCCTGGGTTT |
| | 351 TTCCGACTGT ACCCTGGAAG CGATGCGTGC GCTGGCCCGT TACAGCCTGC AGAACGACAT CCAGACCCTG |
| | 421 GTTACCGTTA CCACCGTTGG TGTTGAAAAA ATGATGATCC GTGCCGGTCT GGACGTTTCG CGTTTCGGTC |
| | 491 CGCACCTGAA AATCGGTATC GAACGTGCGG TTGCCCTGCG CATCGAACTG AATGCTAAAA CCCAGATCGC |
| | 561 GCTGTACGGT GGTGTTCTGG TTGAACAGCG TCTGGCGGTT TCATAA |
| EsaI | 1 ATGCTTGAAC TGTTTGACGT CAGTTACGAA GAACTGCAAA CCACCCGTTC AGAAGAACTT TATAAACTTC |
| | 71 GCAAGAAAAC ATTTAGCGAT CGTCTGGGAT GGGAAGTCAT TTGCAGTCAG GGAATGGAGT CCGATGAATT |
| | 141 TGATGGGCCC GGTACACGTT ATATTCTGGG AATCTGCGAA GGACAATTAG TGTGCAGCGT ACGTTTTACC |
| | 211 AGCCTCGATC GTCCCAACAT GATCACGCAC ACTTTTCAGC ACTGCTTCAG TGATGTCACC CTGCCCGCCT |
| | 281 ATGGTACCGA ATCCAGCCGT TTTTTTGTCG ACAAAGCCCG CGCACGTGCG CTGTTAGGTG AGCACTACCC |
| | 351 TATCAGCCAG GTCCTGTTTT TAGCGATGGT GAACTGGGCG CAAAATAATG CCTACGGCAA TATCTATACG |
| | 421 ATTGTCAGCC GCGCGATGTT GAAAATTCTC ACTCGCTCTG GCTGGCAAAT CAAAGTCATT AAAGAGGCTT |
| | 491 TCCTGACCGA AAAGGAACGT ATCTATTTGC TGACGCTGCC AGCAGGTCAG GATGACAAGC AGCAACTCGG |
| | 561 TGGTGATGTG GTGTCACGTA CGGGCTGTCC GCCCGTCGCA GTCACTACCT GGCCGCTGAC GCTGCCGGTC |
| | 631 TGA |
| LasR(P117S) | 1 ATGGCACTGG TTGATGGTTT TCTGGAACTG GAACGTAGCA GCGGTAAACT GGAATGGTCA GCAATCCTGC |
| | 71 AGAAAATGGC AAGCGATCTG GGTTTTAGCA AAATTCTGTT TGGTCTGCTG CCGAAAGATA GCCAGGATTA |
| | 141 TGAAAATGCC TTTATCGTGG GTAATTATCC GGCAGCATGG CGTGAACATT ATGATCGTGC AGGTTATGCA |
| | 211 CGTGTTGATC CGACCGTTAG CCATTGTACC CAGAGCGTTC TGCCGATTTT TTGGGAACCG AGCATTTATC |
| | 281 AGACCCGTAA ACAGCACGAA TTTTTTGAAG AAGCAAGCGC AGCAGGTCTG GTTTATGGTC TGACCATGTC |
| | 351 ACTGCATGGC GCACGTGGTG AACTGGGTGC ACTGAGCCTG AGCGTTGAAG CAGAAAATCG TGCCGAAGCA |
| | 421 AATCGTTTTA TGGAAAGCGT GCTGCCGACA CTGTGGATGC TGAAAGATTA TGCACTGCAG AGCGGTGCCG |
| | 491 GTCTGGCATT TGAACATCCG GTTAGCAAAC CGGTGGTTCT GACCAGCCGT GAAAAAGAAG TTCTGCAGTG |
| | 561 GTGTGCAATT GGTAAAACCA GCTGGGAAAT TAGCGTTATT TGTAATTGTA GCGAAGCCAA CGTGAACTTT |
| | 631 CATATGGGTA ATATTCGTCG CAAATTTGGT GTTACCAGCC GTCGTGTTGC AGCAATTATG GCAGTTAATC |
| | 701 TGGGTCTGAT TACCCTGTAA |

**Table S3:** Continued overview of the coding sequences used in this research.

| Part | Coding sequence |
|---|---|
| LasR(S129N) | 1 ATGGCACTGG TTGATGGTTT TCTGGAACTG GAACGTAGCA GCGGTAAACT GGAATGGTCA GCAATCCTGC |
| | 71 AGAAAATGGC AAGCGATCTG GGTTTTAGCA AAATTCTGTT TGGTCTGCTG CCGAAAGATA GCCAGGATTA |
| | 141 TGAAAATGCC TTTATCGTGG GTAATTATCC GGCAGCATGG CGTGAACATT ATGATCGTGC AGGTTATGCA |
| | 211 CGTGTTGATC CGACCGTTAG CCATTGTACC CAGAGCGTTC TGCCGATTTT TTGGGAACCG AGCATTTATC |
| | 281 AGACCCGTAA ACAGCACGAA TTTTTTGAAG AAGCAAGCGC AGCAGGTCTG GTTTATGGTC TGACCATGCC |
| | 351 GCTGCATGGC GCACGTGGTG AACTGGGTGC ACTGAATCTG AGCGTTGAAG CAGAAAATCG TGCCGAAGCA |
| | 421 AATCGTTTTA TGGAAAGCGT GCTGCCGACA CTGTGGATGC TGAAAGATTA TGCACTGCAG AGCGGTGCCG |
| | 491 GTCTGGCATT TGAACATCCG GTTAGCAAAC CGGTGGTTCT GACCAGCCGT GAAAAAGAAG TTCTGCAGTG |
| | 561 GTGTGCAATT GGTAAAACCA GCTGGGAAAT TAGCGTTATT TGTAATTGTA GCGAAGCCAA CGTGAACTTT |
| | 631 CATATGGGTA ATATTCGTCG CAAATTTGGT GTTACCAGCC GTCGTGTTGC AGCAATTATG GCAGTTAATC |
| | 701 TGGGTCTGAT TACCCTGTAA |
| LasR(T222I) | 1 ATGGCACTGG TTGATGGTTT TCTGGAACTG GAACGTAGCA GCGGTAAACT GGAATGGTCA GCAATCCTGC |
| | 71 AGAAAATGGC AAGCGATCTG GGTTTTAGCA AAATTCTGTT TGGTCTGCTG CCGAAAGATA GCCAGGATTA |
| | 141 TGAAAATGCC TTTATCGTGG GTAATTATCC GGCAGCATGG CGTGAACATT ATGATCGTGC AGGTTATGCA |
| | 211 CGTGTTGATC CGACCGTTAG CCATTGTACC CAGAGCGTTC TGCCGATTTT TTGGGAACCG AGCATTTATC |
| | 281 AGACCCGTAA ACAGCACGAA TTTTTTGAAG AAGCAAGCGC AGCAGGTCTG GTTTATGGTC TGACCATGCC |
| | 351 GCTGCATGGC GCACGTGGTG AACTGGGTGC ACTGAGCCTG AGCGTTGAAG CAGAAAATCG TGCCGAAGCA |
| | 421 AATCGTTTTA TGGAAAGCGT GCTGCCGACA CTGTGGATGC TGAAAGATTA TGCACTGCAG AGCGGTGCCG |
| | 491 GTCTGGCATT TGAACATCCG GTTAGCAAAC CGGTGGTTCT GACCAGCCGT GAAAAAGAAG TTCTGCAGTG |
| | 561 GTGTGCAATT GGTAAAACCA GCTGGGAAAT TAGCGTTATT TGTAATTGTA GCGAAGCCAA CGTGAACTTT |
| | 631 CATATGGGTA ATATTCGTCG CAAATTTGGT GTTATTAGCC GTCGTGTTGC AGCAATTATG GCAGTTAATC |
| | 701 TGGGTCTGAT TACCCTGTAA |

**Table S3:** Continued overview of the coding sequences used in this research.

| Part | Coding sequence |
|---|---|
| sfGFP | 1 ATGAGCAAGG GCGAAGAGCT TTTTACCGGT GTTGTGCCGA TTTTAGTAGA ACTGGACGGA GACGTGAACG |
| | 71 GTCATAAGTT CTCTGTTCGT GGCGAAGGAG AGGGAGATGC CACCAATGGT AAGCTGACCC TGAAGTTCAT |
| | 141 CTGTACCACC GGTAAGCTGC CCGTGCCTTG GCCGACGCTG GTCACAACGT TGACGTATGG CGTCCAATGC |
| | 211 TTTTCACGCT ATCCAGATCA CATGAAACGC CACGACTTTT TTAAAAGCGC AATGCCTGAA GGTTATGTGC |
| | 281 AGGAACGGAC TATTAGCTTC AAAGACGATG GGACGTATAA GACCCGCGCG GAAGTGAAAT TTGAAGGCGA |
| | 351 TACCTTAGTT AACCGCATTG AATTAAAAGG TATCGATTTC AAAGAGGATG GGAATATCCT GGGGCACAAA |
| | 421 TTGGAATACA ACTTTAATTC GCACAACGTA TACATTACAG CGGATAAACA GAAAAATGGC ATCAAAGCCA |
| | 491 ACTTTAAAAT CCGTCATAAC GTAGAGGACG GTTCCGTGCA GCTGGCTGAT CATTACCAGC AGAATACTCC |
| | 561 GATTGGCGAT GGCCCCGTTC TGCTCCCGGA TAATCATTAC CTGTCTACAC AAAGCGTTCT TAGTAAAGAC |
| | 631 CCAAACGAGA AGCGTGACCA TATGGTCCTG TTGGAATTCG TCACGGCAGC GGGGATTACT CATGGCATGG |
| | 701 ATGAACTCTA TAAGTAA |
| sfGFP(LVA) | 1 ATGAGCAAGG GCGAAGAGCT TTTTACCGGT GTTGTGCCGA TTTTAGTAGA ACTGGACGGA GACGTGAACG |
| | 71 GTCATAAGTT CTCTGTTCGT GGCGAAGGAG AGGGAGATGC CACCAATGGT AAGCTGACCC TGAAGTTCAT |
| | 141 CTGTACCACC GGTAAGCTGC CCGTGCCTTG GCCGACGCTG GTCACAACGT TGACGTATGG CGTCCAATGC |
| | 211 TTTTCACGCT ATCCAGATCA CATGAAACGC CACGACTTTT TTAAAAGCGC AATGCCTGAA GGTTATGTGC |
| | 281 AGGAACGGAC TATTAGCTTC AAAGACGATG GGACGTATAA GACCCGCGCG GAAGTGAAAT TTGAAGGCGA |
| | 351 TACCTTAGTT AACCGCATTG AATTAAAAGG TATCGATTTC AAAGAGGATG GGAATATCCT GGGGCACAAA |
| | 421 TTGGAATACA ACTTTAATTC GCACAACGTA TACATTACAG CGGATAAACA GAAAAATGGC ATCAAAGCCA |
| | 491 ACTTTAAAAT CCGTCATAAC GTAGAGGACG GTTCCGTGCA GCTGGCTGAT CATTACCAGC AGAATACTCC |
| | 561 GATTGGCGAT GGCCCCGTTC TGCTCCCGGA TAATCATTAC CTGTCTACAC AAAGCGTTCT TAGTAAAGAC |
| | 631 CCAAACGAGA AGCGTGACCA TATGGTCCTG TTGGAATTCG TCACGGCAGC GGGGATTACT CATGGCATGG |
| | 701 ATGAACTCTA TAAGGCAGCA AACGACGAAA ACTACGCTTT AGTAGCTTAA |

**Table S3:** Continued overview of the coding sequences used in this research.

| Part | Coding sequence |
|---|---|
| mKate2 | 1 ATGGTTAGCG AGCTGATCAA AGAAAACATG CACATGAAAC TGTATATGGA AGGCACCGTG AATAACCACC |
| | 71 ACTTTAAATG TACCAGCGAA GGTGAAGGTA AACCGTATGA AGGCACCCAG ACCATGCGTA TTAAAGCAGT |
| | 141 TGAAGGTGGT CCGCTGCCGT TTGCATTTGA TATTCTGGCA ACCAGCTTTA TGTATGGCAG CAAAACCTTT |
| | 211 ATTAACCATA CCCAGGGTAT CCCGGATTTT TTCAAACAGA GCTTTCCGGA AGGTTTTACC TGGGAACGTG |
| | 281 TTACCACCTA TGAAGATGGT GGTGTTCTGA CCGCAACCCA GGATACCAGT CTGCAGGATG GTTGTCTGAT |
| | 351 TTATAATGTG AAAATTCGCG GTGTGAACTT TCCGAGCAAT GGTCCGGTTA TGCAGAAAAA AACCCTGGGT |
| | 421 TGGGAAGCAA GCACCGAAAC CCTGTATCCG GCAGATGGTG GTCTGGAAGG TCGTGCAGAT ATGGCACTGA |
| | 491 AACTGGTTGG TGGTGGTCAT CTGATTTGCA ATCTGAAAAC CACCTATCGT AGCAAAAAAC CGGCAAAAAA |
| | 561 TCTGAAAATG CCTGGCGTGT ATTATGTTGA TCGTCGTCTG GAACGTATTA AAGAGGCAGA TAAAGAAACC |
| | 631 TATGTGGAAC AGCATGAAGT TGCAGTTGCA CGTTATTGTG ATCTGCCGAG CAAACTGGGT CACCGCTGA |

**Table S4:** Overview of the DNA sequence of all primers used for plasmid construction in this research.

| Primer | Sequence (5' → 3') |
|---|---|
| oMEMO9997_Rv_EsaR_sfGFPoverhang | CATACCAGAACCACCACCAGAACCACCCCTTGCAGCTGATGCTGCCGGTCTGATAAG |
| oMEMO9998_Rv_LasR_sfGFPoverhang | CATACCAGAACCACCACCAGAACCACCCAGGGTAATCAGACCCAGATTAACTGCC |
| oMEMO9999_Rv_BB_sfGFPoverhang_pMoBioS-M4 | ACTCATGGCATGGATGAACTCTATAAGTAACAATAGTCTTTCAGGGCCGTATGCAC |
| oMEMO10083_Rv_BB_pMoBioS-M4 | GAGTGGTGATTGATTGAGC |
| oMEMO10084_Fw_SacB_BsaI | GCTCAATCAATCACCACTCAAGGCGAGACCGGCCGCCTTCATTCTATAAG |
| oMEMO10085_Rv_SacB_BsaI | GCCTACACGGGAGAGTGTTCACTCCGAGACCCCACACTAC |
| oMEMO10098_Rv_SacB_BsaI_GG2 | GAACACTCTCCCGTGTAGGCCACTCCGAGACCCCACACTAC |
| oMEMO10101_Rv_internal_pMoBioS | CCGCTAGCCCATGGTTATC |
| oMEMO10152_Fw_BsaI_pMoBioS | TGTGGGGTCTCGGAGTGGCCTACACGGGAGAGTGTTC |
| oMEMO10153_Rv_BsaI_pMoBioS | GCGGCCGGTCTCGCCTTGAGTGGTGATTGATTGAGC |
| oMEMO10235_Fw_RBSlib_EsaI | GTCCTAGGTATAATGCTAGCACACGWTMTTSGAAGGAVGTACATATGCTTGAACT-GTTTGACG |
| oMEMO10236_Rv_J23108_overlap | CGTGTGCTAGCATTATACCTAGGAC |
| oMEMO10237_Fw_EsaI_overlap | GTACATATGCTTGAACTGTTTGACG |
| oMEMO10263_sfGFP_ssrA_RV | CGAGCATTCACCTGCGGTACTTATTAAGCTACTAAAGCGTAGTTTTCGTCGTTTGCTGCCT-TATAGAGTTCATCCATGCCATGAG |
| oMEMO10289_RBS_EsaI | GTCCTAGGTATAATGCTAGCACACGATCTTCGAAGGACGTACATATGCTTGAACT-GTTTGACG |
| oMEMO10335_Fw_LasR_T222I | GCAACACGACGGCTAATAACACCAAATTTGC |
| oMEMO10336_Rv_LasR_T222I | CAAATTTGGTGTTATTAGCCGTCGTGTTGC |
| oMEMO10337_Fw_LasR_S129N | GCTTCAACGCTCAGATTCAGTGCACCCAG |
| oMEMO10338_Rv_LasR_S129N | CTGGGTGCACTGAATCTGAGCGTTGAAGC |
| oMEMO10339_Fw_LasR_P117S | GTGCGCCATGCAGTGACATGGTCAGACCATAAAC |
| oMEMO10340_Rv_LasR_P117S | GTTTATGGTCTGACCATGTCACTGCATGGCGCAC |
| oMEMO11118_Pesa_orth_overhang_Fw | CATCTTGCCTGTACTATAGTGCAAGTTAAGTCCACGTTAAGTAAAAGAAGCAGC |
| oMEMO11119_Pesa_orth_Rv | TGCACTATAGTACAGGCAAGATG |
| **Internal primers** | |
| oMEMO8529_Rv_BB1 | CTCGGATGGAAGCCGGTCTTGTCG |
| oMEMO8530_Fw_BB2 | CGACAAGACCGGCTTCCATCCGAG |
| oMEMO9754_pBBR1MCS2-split_FW | CCTACCGCATGGAGATAAGC |
| oMEMO9755_pBBR1MCS2-split_RV | GCTTATCTCCATGCGGTAGGG |
| oMEMO10101_Rv_internal_pMoBioS | CCGCTAGCCCATGGTTATC |
| oMEMO9662_pBBR1MCS2-Psyn-BB-FW | TAACCATGGGCTAGCGGTTTG |

**Table S5:** Overview of the sequences in the RBS-library used to control the translation of *esaI*. TIR = translation initiation rate.

| **Sequence** | **TIR** |
|---|---|
| ACACG**W**T**M**TT**S**GAAGGA**V**GTACAT | Library |
| ACACGATATTGGAAGGAGGTACAT | 35252.33216 |
| ACACGTTATTGGAAGGAGGTACAT | 21584.65324 |
| ACACGATATTCGAAGGAGGTACAT | 17867.1502 |
| ACACGATCTTGGAAGGAGGTACAT | 17867.1502 |
| ACACGTTATTCGAAGGAGGTACAT | 9472.569872 |
| ACACGTTCTTGGAAGGAGGTACAT | 5324.627492 |
| ACACGATCTTCGAAGGAGGTACAT | 2822.941332 |
| ACACGATATTGGAAGGAAGTACAT | 2615.018711 |
| ACACGATATTCGAAGGAAGTACAT | 2014.246509 |
| ACACGATCTTGGAAGGAAGTACAT | 1608.372313 |
| ACACGTTATTGGAAGGAAGTACAT | 1319.433997 |
| ACACGTTATTCGAAGGAAGTACAT | 1163.218083 |
| ACACGATATTGGAAGGACGTACAT | 954.2501268 |
| ACACGTTATTGGAAGGACGTACAT | 864.2964327 |
| ACACGTTCTTCGAAGGAGGTACAT | 841.2705793 |
| ACACGTTCTTGGAAGGAAGTACAT | 653.8565285 |
| ACACGATCTTCGAAGGAAGTACAT | 346.6530987 |
| ACACGATATTCGAAGGACGTACAT | 284.378052 |
| ACACGATCTTGGAAGGACGTACAT | 284.378052 |
| ACACGTTATTCGAAGGACGTACAT | 150.7678346 |
| ACACGTTCTTCGAAGGAAGTACAT | 103.3068062 |
| ACACGTTCTTGGAAGGACGTACAT | 84.74812726 |
| ACACGATCTTCGAAGGACGTACAT | 44.9306532 |
| ACACGTTCTTCGAAGGACGTACAT | 13.38988255 |

**Table S6:** Statistical comparison of the binding of LasR to $P_{esaR}$. This is assessed by the addition of different concentrations of 3OC12-HSL to form the ligand-bound active transcription factor complex. One-way ANOVA was performed and Tukey HSD was conducted to correct for multiple comparison. Significant p-values are highlighted in bold.

| **Statistics** | | |
|---|---|---|
| ANOVA:<br>F = 6.663253<br>p = **0.000847** | | |
| **Concentration group 1 (nM)** | **Concentration group 2 (nM)** | **p-adj** |
| 0 | 0.01 | 1 |
| 0 | 0.1 | 1 |
| 0 | 1 | 1 |
| 0 | 10 | 1 |
| 0 | 100 | 0.349 |
| 0 | 1000 | 0.0688 |
| 0 | 10000 | **0.0075** |
| 0.01 | 0.1 | 1 |
| 0.01 | 1 | 1 |
| 0.01 | 10 | 1 |
| 0.01 | 100 | 0.33 |
| 0.01 | 1000 | 0.0638 |
| 0.01 | 10000 | **0.0069** |
| 0.1 | 1 | 1 |
| 0.1 | 10 | 0.9999 |
| 0.1 | 100 | 0.3019 |
| 0.1 | 1000 | 0.0569 |
| 0.1 | 10000 | **0.0061** |
| 1 | 10 | 0.9998 |
| 1 | 100 | 0.2921 |
| 1 | 1000 | 0.0545 |
| 1 | 10000 | **0.0059** |
| 10 | 100 | 0.5133 |
| 10 | 1000 | 0.1191 |
| 10 | 10000 | **0.0136** |
| 100 | 1000 | 0.9698 |
| 100 | 10000 | 0.4302 |
| 1000 | 10000 | 0.9335 |

**Table S7:** Statistical comparison of the binding of LasR to $P_{esaS}$. This is assessed by the addition of different concentrations of 3OC12-HSL to form the ligand-bound active transcription factor complex. One-way ANOVA was performed and Tukey HSD was conducted to correct for multiple comparison. Significant p-values are highlighted in bold.

| **Statistics** | | |
|---|---|---|
| ANOVA:<br>F = 8.649191<br>p = **0.000194** | | |
| **Concentration group 1 (nM)** | **Concentration group 2 (nM)** | **p-adj** |
| 0 | 0.01 | 1 |
| 0 | 0.1 | 1 |
| 0 | 1 | 0.9993 |
| 0 | 10 | 0.9421 |
| 0 | 100 | 0.0598 |
| 0 | 1000 | **0.001** |
| 0 | 10000 | **0.034** |
| 0.01 | 0.1 | 1 |
| 0.01 | 1 | 0.9999 |
| 0.01 | 10 | 0.9661 |
| 0.01 | 100 | 0.0737 |
| 0.01 | 1000 | **0.0013** |
| 0.01 | 10000 | 0.0422 |
| 0.1 | 1 | 0.9999 |
| 0.1 | 10 | 0.9737 |
| 0.1 | 100 | 0.0803 |
| 0.1 | 1000 | **0.0014** |
| 0.1 | 10000 | **0.0461** |
| 1 | 10 | 0.9983 |
| 1 | 100 | 0.1509 |
| 1 | 1000 | **0.0028** |
| 1 | 10000 | 0.0894 |
| 10 | 100 | 0.3783 |
| 10 | 1000 | **0.0091** |
| 10 | 10000 | 0.2458 |
| 100 | 1000 | 0.4516 |
| 100 | 10000 | 1 |
| 1000 | 10000 | 0.6262 |

**Table S8:** One-way ANOVA was performed for the statistical comparison of the binding of EsaR to $P_{lasI}$. This is assessed by the addition of different concentrations of 3OC6-HSL, to form the ligand-bound active transcription factor complex.

| **Statistics** |
|---|
| ANOVA:<br>F = 1.011252<br>p = 0.459494 |

**Table S9:** Statistical assessment of the signal crosstalk of LasR to 3OC6-HSL. This is assessed by looking at the promoter activity of $P_{lasI}$. One-way ANOVA was performed and Tukey HSD was conducted to correct for multiple comparison. Significant p-values are highlighted in bold.

| **Statistics** | | |
|---|---|---|
| ANOVA:<br>F = 136.431781<br>p = **1.328327e-11** | | |
| **Concentration group 1 (nM)** | **Concentration group 2 (nM)** | **p-adj** |
| 0.01 | 0.1 | 0.6189 |
| 0.01 | 1 | 0.9982 |
| 0.01 | 10 | 0.975 |
| 0.01 | 100 | 0.9917 |
| 0.01 | 1000 | 1 |
| 0.01 | 10000 | **0** |
| 0.1 | 1 | 0.341 |
| 0.1 | 10 | 0.2103 |
| 0.1 | 100 | 0.9375 |
| 0.1 | 1000 | 0.7229 |
| 0.1 | 10000 | **0** |
| 1 | 10 | 0.9998 |
| 1 | 100 | 0.8852 |
| 1 | 1000 | 0.9908 |
| 1 | 10000 | **0** |
| 10 | 100 | 0.7255 |
| 10 | 1000 | 0.9387 |
| 10 | 10000 | **0** |
| 100 | 1000 | 0.9984 |
| 100 | 10000 | **0** |
| 1000 | 10000 | **0** |

**Table S10:** Statistical assessment of the signal crosstalk of LasR to 3OC8-HSL. This is assessed by looking at the promoter activity of $P_{lasI}$. One-way ANOVA was performed and Tukey HSD was conducted to correct for multiple comparison. Significant p-values are highlighted in bold.

| **Statistics** | | |
|---|---|---|
| ANOVA:<br>F = 1588.835419<br>p = **5.236138e-19** | | |
| **Concentration group 1 (nM)** | **Concentration group 2 (nM)** | **p-adj** |
| 0.01 | 0.1 | 0.998 |
| 0.01 | 1 | 0.5713 |
| 0.01 | 10 | **0** |
| 0.01 | 100 | **0** |
| 0.01 | 1000 | **0** |
| 0.01 | 10000 | **0** |
| 0.1 | 1 | 0.8532 |
| 0.1 | 10 | **0** |
| 0.1 | 100 | **0** |
| 0.1 | 1000 | **0** |
| 0.1 | 10000 | **0** |
| 1 | 10 | **0** |
| 1 | 100 | **0** |
| 1 | 1000 | **0** |
| 1 | 10000 | **0** |
| 10 | 100 | **0** |
| 10 | 1000 | **0** |
| 10 | 10000 | **0** |
| 100 | 1000 | **0.0001** |
| 100 | 10000 | **0** |
| 1000 | 10000 | **0** |

**Table S11:** Statistical assessment of the signal crosstalk of LasR to 3OC10-HSL. This is assessed by looking at the promoter activity of $P_{lasI}$. One-way ANOVA was performed and Tukey HSD was conducted to correct for multiple comparison. Significant p-values are highlighted in bold.

| **Statistics** | | |
|---|---|---|
| ANOVA:<br>F = 1209.44445<br>p = **3.521393e-18** | | |
| **Concentration group 1 (nM)** | **Concentration group 2 (nM)** | **p-adj** |
| 0 | 0.01 | **0.0003** |
| 0 | 0.1 | **0** |
| 0 | 1 | **0** |
| 0 | 10 | **0** |
| 0 | 100 | **0** |
| 0 | 1000 | **0** |
| 0 | 10000 | **0** |
| 0.01 | 0.1 | **0** |
| 0.01 | 1 | **0** |
| 0.01 | 10 | **0** |
| 0.01 | 100 | **0** |
| 0.01 | 1000 | **0** |
| 0.01 | 10000 | 0.3459 |
| 0.1 | 1 | **0** |
| 0.1 | 10 | **0.0012** |
| 0.1 | 100 | **0** |
| 0.1 | 1000 | **0** |
| 0.1 | 10000 | **0** |
| 1 | 10 | **0** |
| 1 | 100 | **0** |
| 1 | 1000 | 0.097 |
| 1 | 10000 | **0.0163** |
| 10 | 100 | 0.9997 |
| 10 | 1000 | 0.9777 |
| 10 | 10000 | 0.0532 |
| 100 | 1000 | 0.9997 |
| 100 | 10000 | 0.1244 |
| 1000 | 10000 | 0.2642 |

**Table S12:** Statistical assessment of the signal crosstalk of EsaR to 3OC8-HSL. This is assessed by looking at the promoter activity of $P_{esaR}$. One-way ANOVA was performed and Tukey HSD was conducted to correct for multiple comparison. Significant p-values are highlighted in bold.

| **Statistics** | | |
|---|---|---|
| ANOVA:<br>F = 555.175443<br>p = **9.273916e-14** | | |
| **Concentration group 1 (nM)** | **Concentration group 2 (nM)** | **p-adj** |
| 0.1 | 1 | 0.9999 |
| 0.1 | 10 | 0.9931 |
| 0.1 | 100 | **0** |
| 0.1 | 1000 | **0** |
| 0.1 | 10000 | **0** |
| 1 | 10 | 0.9995 |
| 1 | 100 | **0.0001** |
| 1 | 1000 | **0** |
| 1 | 10000 | **0** |
| 10 | 100 | **0.0001** |
| 10 | 1000 | **0** |
| 10 | 10000 | **0** |
| 100 | 1000 | **0** |
| 100 | 10000 | **0** |
| 1000 | 10000 | **0** |

**Table S13:** Statistical assessment of the signal crosstalk of EsaR to 3OC10-HSL. This is assessed by looking at the promoter activity of $P_{esaR}$. One-way ANOVA was performed and Tukey HSD was conducted to correct for multiple comparison. Significant p-values are highlighted in bold.

| **Statistics** | | |
|---|---|---|
| ANOVA:<br>F = 102.116977<br>p = **2.086524e-09** | | |
| **Concentration group 1 (nM)** | **Concentration group 2 (nM)** | **p-adj** |
| 0.1 | 1 | 1 |
| 0.1 | 10 | 0.9565 |
| 0.1 | 100 | 0.6141 |
| 0.1 | 1000 | **0.0626** |
| 0.1 | 10000 | **0** |
| 1 | 10 | 0.905 |
| 1 | 100 | 0.5089 |
| 1 | 1000 | **0.0459** |
| 1 | 10000 | **0** |
| 10 | 100 | 0.9674 |
| 10 | 1000 | 0.231 |
| 10 | 10000 | **0** |
| 100 | 1000 | 0.5989 |
| 100 | 10000 | **0** |
| 1000 | 10000 | **0** |

**Table S14:** One-way ANOVA was performed for the statistical assessment of the signal crosstalk of EsaR to 3OC12-HSL. This is assessed by looking at the promoter activity of $P_{esaR}$.

| **Statistics** |
|---|
| ANOVA:<br>F = 1.918009<br>p = 0.132925 |

**Table S15:** Statistical assessment of the signal crosstalk of EsaR to 3OC8-HSL. This is assessed by looking at the promoter activity of $P_{esaS}$. One-way ANOVA was performed and Tukey HSD was conducted to correct for multiple comparison. Significant p-values are highlighted in bold.

| **Statistics** | | |
|---|---|---|
| ANOVA:<br>F = 1119.15288<br>p = **1.404036e-15** | | |
| **Concentration group 1 (nM)** | **Concentration group 2 (nM)** | **p-adj** |
| 0.1 | 1 | 0.4173 |
| 0.1 | 10 | 0.1226 |
| 0.1 | 100 | **0** |
| 0.1 | 1000 | **0** |
| 0.1 | 10000 | **0** |
| 1 | 10 | 0.9503 |
| 1 | 100 | **0** |
| 1 | 1000 | **0** |
| 1 | 10000 | **0** |
| 10 | 100 | **0** |
| 10 | 1000 | **0** |
| 10 | 10000 | **0** |
| 100 | 1000 | **0** |
| 100 | 10000 | **0** |
| 1000 | 10000 | 0.4062 |

**Table S16:** Statistical assessment of the signal crosstalk of EsaR to 3OC10-HSL. This is assessed by looking at the promoter activity of $P_{esaS}$. One-way ANOVA was performed and Tukey HSD was conducted to correct for multiple comparison. Significant p-values are highlighted in bold.

| **Statistics** | | |
|---|---|---|
| ANOVA:<br>F = 15.8041<br>p = **0.000064** | | |
| **Concentration group 1 (nM)** | **Concentration group 2 (nM)** | **p-adj** |
| 0.1 | 1 | 0.9951 |
| 0.1 | 10 | 0.3164 |
| 0.1 | 100 | 0.9758 |
| 0.1 | 1000 | 0.9678 |
| 0.1 | 10000 | **0.0009** |
| 1 | 10 | 0.5738 |
| 1 | 100 | 0.9999 |
| 1 | 1000 | 0.9998 |
| 1 | 10000 | **0.0004** |
| 10 | 100 | 0.6969 |
| 10 | 1000 | 0.7249 |
| 10 | 10000 | **0** |
| 100 | 1000 | 1 |
| 100 | 10000 | **0.0003** |
| 1000 | 10000 | **0.0003** |

**Table S17:** Statistical assessment of the signal crosstalk of EsaR to 3OC12-HSL. This is assessed by looking at the promoter activity of $P_{esaS}$. One-way ANOVA was performed and Tukey HSD was conducted to correct for multiple comparison. Significant p-values are highlighted in bold.

| **Statistics** | | |
|---|---|---|
| ANOVA:<br>F = 4.052984<br>p = **0.009712** | | |
| **Concentration group 1 (nM)** | **Concentration group 2 (nM)** | **p-adj** |
| 0 | 0.01 | 0.959 |
| 0 | 0.1 | 0.9952 |
| 0 | 1 | 0.9988 |
| 0 | 10 | 0.9228 |
| 0 | 100 | 0.709 |
| 0 | 1000 | 0.4397 |
| 0 | 10000 | **0.0053** |
| 0.01 | 0.1 | 1 |
| 0.01 | 1 | 0.9996 |
| 0.01 | 10 | 1 |
| 0.01 | 100 | 0.9981 |
| 0.01 | 1000 | 0.9526 |
| 0.01 | 10000 | **0.0405** |
| 0.1 | 1 | 1 |
| 0.1 | 10 | 0.9996 |
| 0.1 | 100 | 0.9758 |
| 0.1 | 1000 | 0.8397 |
| 0.1 | 10000 | 0.0214 |
| 1 | 10 | 0.998 |
| 1 | 100 | 0.95 |
| 1 | 1000 | 0.7694 |
| 1 | 10000 | 0.0163 |
| 10 | 100 | 0.9997 |
| 10 | 1000 | 0.9777 |
| 10 | 10000 | 0.0532 |
| 100 | 1000 | 0.9997 |
| 100 | 10000 | 0.1244 |
| 1000 | 10000 | 0.2642 |

**Table S18:** Two sample t-test for the assessment of synthase crosstalk. Strains with the promoter and transcription factor are compared with strains that have either one of the synthase proteins or only the promoter region. Significant p-values (< 0.05) are highlighted in bold.

| Strain 1 | Strain 2 | t-value | p-value |
|---|---|---|---|
| **PlasI** | | | |
| LasR-PlasI | PlasI | -33.006995139325184 | **0.000857019530233585** |
| | LasI-LasR-PlasI | 111.26981823098576 | **3.49240710160141e-05** |
| | EsaI-LasR-PlasI | 57.555966426934 | **0.00010750460662436118** |
| EsaR-PlasI | PlasI | -1.8122340908375705 | 0.15901963425131524 |
| | LasI-EsaR-PlasI | 13.885947003419048 | **0.0002472113450992815** |
| | EsaI-EsaR-PlasI | 2.67990101665882 | 0.07760134811177463 |
| **PesaR** | | | |
| LasR-PesaR | PesaR | -0.9468785298482081 | 0.44304713420048997 |
| | LasI-LasR-PesaR | 0.7761049617000624 | 0.482000462097577 |
| | EsaI-LasR-PesaR | -0.5455454785989141 | 0.6386217143287847 |
| EsaR-PesaR | PesaR | 56.58854013633201 | **0.00030230693910849953** |
| | LasI-EsaR-PesaR | -0.13622495968352216 | 0.8982253193583881 |
| | EsaI-EsaR-PesaR | 56.170141258935395 | **0.000284328241347685** |
| **PesaS** | | | |
| LasR-PesaS | PesaS | 0.4012619653072439 | 0.725628355364238 |
| | LasI-LasR-PesaS | 7.141947439145252 | **0.0020462246595919042** |
| | EsaI-LasR-PesaS | 8.038243939482804 | **0.0013017228570557555** |
| EsaR-PesaS | PesaS | -16.094805487429227 | **0.00381360449037832** |
| | LasI-EsaR-PesaS | -0.10814563334415234 | 0.9230349815833094 |
| | EsaI-EsaR-PesaS | -9.856481118082817 | **0.008155164336609136** |

**Table S19:** Two sample t-test for the assessment of the functionality and synthase orthogonality of the new LasR mutants. And the two sample t-test for the assessment of the promoter crosstalk of LasR and LasR(P117S) with the new $P_{esaR/esaS}*$ mutant. Significant p-values (< 0.05) are highlighted in bold.

| Strain 1 | Strain 2 | t-value | p-value |
|---|---|---|---|
| **LasR mutants** | | | |
| LasI-LasR-PlasI | LasI-LasR(P117S)-PlasI | -34.25215157195646 | **0.00017421143133470083** |
| | LasI-LasR(S129N)-PlasI | -1.6744690885797533 | 0.22889886662030032 |
| | LasI-LasR(T222I)-PlasI | -134.58663197738616 | **8.16098974039902e-08** |
| LasR(P117S)-PlasI | EsaI-LasR(P117S)-PlasI | -6.791801504606722 | **0.020721906496406598** |
| **PesaR/esaS mutant** | | | |
| PesaR | LasI-LasR-PesaR | 2.298268564257825 | 0.14712132649685125 |
| PesaR* | LasI-LasR-PesaR* | 10.673875338407004 | **0.0013254507483923075** |
| **LasR(P117S) and PesaR/esaS mutant** | | | |
| PesaR* | LasI-LasR(P117S)-PesaR* | 1.36048727 | 0.27164464 |
