## Supplementary File 2 for "Characterization and orthogonality assessment of two quorum sensing systems for synthetic biology applications"

#### Characterization and orthogonality assessment of two quorum sensing systems for synthetic biology applications

Jasmine De Baets, Brecht De Paepe, Marjan De Mey

*Centre for Synthetic Biology, Ghent university, 9000 Ghent, Belgium*

**Overview**

ID-sheets of LasR and EsaR biosensors with different ribosome binding site sequences. They contain information on the ligand, response curve, meta-data about the strain and experiment and the DNA-sequences of the transcription factor and promoter part.

### LasR:3OC12-HSL

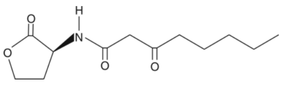


Ligand: 3-oxo-dodecanoyl-L-homoserine lactone (3OC12-HSL)
Source: Sigma-Aldrich O9139
CAS ID: 168982-69-2
Stock: 1 mM
Solvent: DMSO
Storage: 2 months at -80°C

#### Response curve: SynLasR(low)

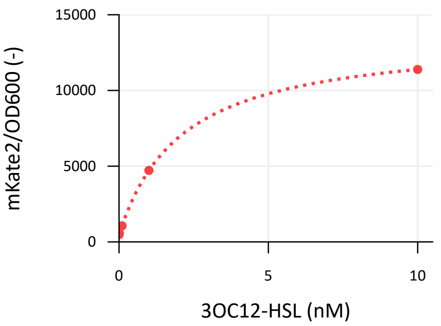


Error bars depict standard errors over four biological replicates (n=3)
a = the basal normalized fluorescent signal (leaky expression, a.u.)
k = the maximum normalized fluorescent signal relative to a (a.u.)
M = a + k = the maximum normalized fluorescent signal (a.u.)
n = the Hill coefficient (cooperativity, sigmoid character)
K = the Hill constant (TF-ligand affinity, nM)
C = the ligand concentration in the growth medium (nM)

**Hill function**

$$\left(\frac{\text{Fluo}}{\overline{\text{OD}_{600}}}\right)_{cor} = f(C) = a + k\left(\frac{C^n}{C^n + K^n}\right)$$

**Parameters Hill function**

$a_{(a.u)} = 4.82*10^2 \pm 0.19*10^2$
$k_{(a.u)} = 1.32*10^4 \pm 0.06*10^4$
$M_{(a.u)} = 1.37*10^4 \pm 0.06*10^4$
$n = 1.01 \pm 0.03$
$K_{(nM)} = 2.09 \pm 0.19$

#### Meta-data

| | |
|---|---|
| Host | *Escherichia coli TOP10 (F- mcrA Δ(mrr-hsdRMS-mcrBC) Φ80lacZΔM15 ΔlacX74 recA1 araD139 Δ(araleu)7697 galU galK rpsL (StrR) endA1 nupG)* |
| Origin | *Pseudomonas aeruginosa PAO1* |
| Parts | LasR; Gene ID: 881789 codon harmonized |
| | $P_{LasI}$; Gene ID: 881777, promoter upstream of *lasI* |
| Architecture | Synthetic |
| Experiment | 150 µL EZ Rich medium, 96-well black microtiter plate<br>PerkinElmer Ensight, exc. 588nm, emm. 633nm at 30°C<br>Fluorescence after …h of growth (early stationary phase) |
| Notes | - |

#### LasR:3OC12-HSL

| | |
|---|---|
| *lasR* | GATGATGGCGAGCATTCACCTGCGGTAGCTA**TTACAGGGTAATCAGACCCAGATTAACT**<br>PaqCI recognition sequence PaqCI restriction site<br>**GCCATAATTGCTGCAACACGACGGCTGGTAACACCAAATTTGCGACGAATATTACCC**<br>**ATATGAAAGTTCACGTTGGCTTCGCTACAATTACAAATAACGCTAATTTCCCAGCTG**<br>**GTTTTACCAATTGCACACCACTGCAGAACTTCTTTTTCACGGCTGGTCAGAACCACC**<br>**GGTTTGCTAACCGGATGTTCAAATGCCAGACCGGCACCGCTCTGCAGTGCATAATCT**<br>**TTCAGCATCCACAGTGTCGGCAGCACGCTTTCCATAAAACGATTTGCTTCGGCACGA**<br>**TTTTCTGCTTCAACGCTCAGGCTCAGTGCACCCAGTTCACCACGTGCGCCATGCAGC**<br>*lasR*<br>**GGCATGGTCAGACCATAAACCAGACCTGCTGCGCTTGCTTCTTCAAAAAATTCGTGC**<br>**TGTTTACGGGTCTGATAAATGCTCGGTTCCCAAAAAATCGGCAGAACGCTCTGGGTA**<br>**CAATGGCTAACGGTCGGATCAACACGTGCATAACCTGCACGATCATAATGTTCACGC**<br>**CATGCTGCCGGATAATTACCCACGATAAAGGCATTTTCATAATCCTGGCTATCTTTC**<br>**GGCAGCAGACCAAACAGAATTTTGCTAAAACCCAGATCGCTTGCCATTTTCTGCAGG**<br>**ATTGCTGACCATTCCAGTTTACCGCTGCTACGTTCCAGTTCCAGAAAACCATCAACC**<br>←<br>**AGTGCCAT**AGAGACGCGTCCTTAGTTTGGTTGTTGTTAGCGGGTAGTGTGACGTGGCGCAG<br>BsaI restriction site BsaI recognition sequence Golden RBS PaqCI restriction site<br>GTGATGGACTTCATGCTGAC<br>PaqCI recognition sequence |
| $P_{lasI}$ | GATGATGGCGAGCATTCACCTGCGGTACTTA**TTCGAGCCTAGCAAGGGTCCGGGTTCAC**<br>PaqCI recognition sequence PaqCI restriction site<br>**CGAAATCTATCTCATTTGCTAGTTATAAAATTATGAAATTTGCATAAATTCTTCA**TCT<br>$P_{LasI}$<br>AGAGATTAAAGAGGAGAAATACTAGATGGTGGCGCAGGTGATGGACTTCATGCTGAC<br>Bba_B0030 PaqCI restriction site PaqCI recognition sequence |

#### LasR:3OC12-HSL

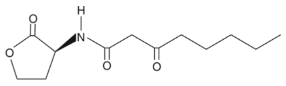


Ligand: 3-oxo-dodecanoyl-L-homoserine lactone (3OC12-HSL)
Source: Sigma-Aldrich O9139
CAS ID: 168982-69-2
Stock: 1 mM
Solvent: DMSO
Storage: 2 months at -80°C

##### Response curve: SynLasR(medium)

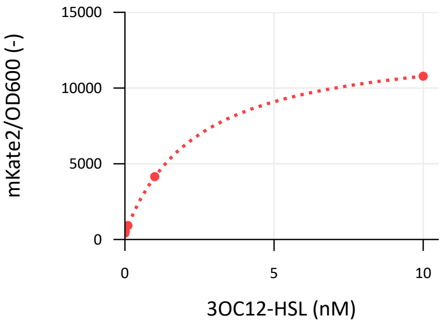


**Hill function**

$$\left(\frac{\text{Fluo}}{\text{OD}_{600}}\right)_{cor} = f\,(C) = a + k\left(\frac{C^n}{C^n+K^n}\right)$$

**Parameters Hill function**

$a_{\,(a.u)} = 4.61*10^2 \pm 0.39*10^2$
$k_{\,(a.u)} = 1.27*10^4 \pm 0.08*10^4$
$M_{\,(a.u)} = 1.32*10^4 \pm 0.08*10^4$
$n = 1.02 \pm 0.09$
$K_{\,(nM)} = 2.41 \pm 0.48$

Error bars depict standard errors over four biological replicates (n=3)
a = the basal normalized fluorescent signal (leaky expression, a.u.)
k = the maximum normalized fluorescent signal relative to a (a.u.)
M = a + k = the maximum normalized fluorescent signal (a.u.)
n = the Hill coefficient (cooperativity, sigmoid character)
K = the Hill constant (TF-ligand affinity, nM)
C = the ligand concentration in the growth medium (nM)

##### Meta-data

| | |
|---|---|
| Host | *Escherichia coli TOP10 (F- mcrA Δ(mrr-hsdRMS-mcrBC) Φ80lacZΔM15 ΔlacX74 recA1 araD139 Δ(araleu)7697 galU galK rpsL (StrR) endA1 nupG)* |
| Origin | *Pseudomonas aeruginosa PAO1* |
| Parts | LasR; Gene ID: 881789 codon harmonized |
| | $P_{LasI}$; Gene ID: 881777, promoter upstream of *lasI* |
| Architecture | Synthetic |
| Experiment | 150 µL EZ Rich medium, 96-well black microtiter plate<br>PerkinElmer Ensight, exc. 588nm, emm. 633nm at 30°C<br>Fluorescence after ...h of growth (early stationary phase) |
| Notes | - |

#### LasR:3OC12-HSL

| | |
|---|---|
| *lasR* | GATGATGGCGAGCATTCACCTGCGGTAGCTA**TTACAGGGTAATCAGACCCAGATTAACT**<br>PaqCI recognition sequence PaqCI restriction site<br>**GCCATAATTGCTGCAACACGACGGCTGGTAACACCAAATTTGCGACGAATATTACCC**<br>**ATATGAAAGTTCACGTTGGCTTCGCTACAATTACAAATAACGCTAATTTCCCAGCTG**<br>**GTTTTACCAATTGCACACCACTGCAGAACTTCTTTTTCACGGCTGGTCAGAACCACC**<br>**GGTTTGCTAACCGGATGTTCAAATGCCAGACCGGCACCGCTCTGCAGTGCATAATCT**<br>**TTCAGCATCCACAGTGTCGGCAGCACGCTTTCCATAAAACGATTTGCTTCGGCACGA**<br>**TTTTCTGCTTCAACGCTCAGGCTCAGTGCACCCAGTTCACCACGTGCGCCATGCAGC**<br>*lasR*<br>**GGCATGGTCAGACCATAAACCAGACCTGCTGCGCTTGCTTCTTCAAAAAATTCGTGC**<br>**TGTTTACGGGTCTGATAAATGCTCGGTTCCCAAAAAATCGGCAGAACGCTCTGGGTA**<br>**CAATGGCTAACGGTCGGATCAACACGTGCATAACCTGCACGATCATAATGTTCACGC**<br>**CATGCTGCCGGATAATTACCCACGATAAAGGCATTTTCATAATCCTGGCTATCTTTC**<br>**GGCAGCAGACCAAACAGAATTTTGCTAAAACCCAGATCGCTTGCCATTTTCTGCAGG**<br>**ATTGCTGACCATTCCAGTTTACCGCTGCTACGTTCCAGTTCCAGAAAACCATCAACC**<br>**AGTGCCAT**AGAGACGACCCCTGTCTTCCTCAACCCGTAGCGGGTAGTGTGACGTGGCGCA<br>BsaI restriction site BsaI recognition sequence Golden RBS PaqCI restriction site<br>GGTGATGGACTTCATGCTGAC<br>PaqCI recognition sequence |
| PlasI | GATGATGGCGAGCATTCACCTGCGGTACTTA**TTCGAGCCTAGCAAGGGTCCGGGTTCAC**<br>PaqCI recognition sequence PaqCI restriction site<br>**CGAAATCTATCTCATTTGCTAGTTATAAAATTATGAAATTTGCATAAATTCTTCA**TCT<br>$P_{LasI}$<br>AGAGATTAAAGAGGAGAAATACTAGATGGTGGCGCAGGTGATGGACTTCATGCTGAC<br>Bba_B0030 PaqCI restriction site PaqCI recognition sequence |

#### LasR:3OC12-HSL

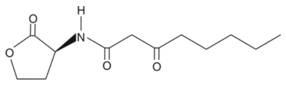


Ligand: 3-oxo-dodecanoyl-L-homoserine lactone (3OC12-HSL)
Source: Sigma-Aldrich O9139
CAS ID: 168982-69-2
Stock: 1 mM
Solvent: DMSO
Storage: 2 months at -80°C

##### Response curve: SynLasR(high)

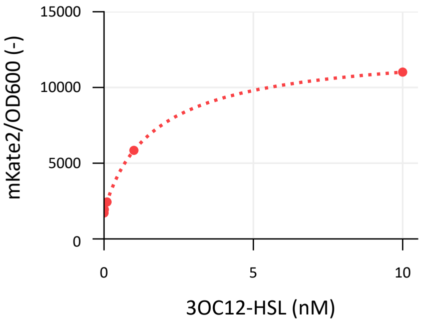


**Hill function**

$$\left(\frac{\text{Fluo}}{\text{OD}_{600}}\right)_{cor} = f\,(C) = a + k\left(\frac{C^n}{C^n + K^n}\right)$$

**Parameters Hill function**

$a_{\text{(a.u)}} = 1.77*10^3 \pm 0.08*10^3$
$k_{\text{(a.u)}} = 1.11*10^4 \pm 0.06*10^4$
$M_{\text{(a.u)}} = 1.29*10^4 \pm 0.07*10^4$
$n = 9.42*10^{-1} \pm 0.98*10^{-1}$
$K_{\text{(nM)}} = 1.77 \pm 0.28$

Error bars depict standard errors over four biological replicates (n=3)
a = the basal normalized fluorescent signal (leaky expression, a.u.)
k = the maximum normalized fluorescent signal relative to a (a.u.)
M = a + k = the maximum normalized fluorescent signal (a.u.)
n = the Hill coefficient (cooperativity, sigmoid character)
K = the Hill constant (TF-ligand affinity, nM)
C = the ligand concentration in the growth medium (nM)

##### Meta-data

| | |
|---|---|
| Host | *Escherichia coli TOP10 (F- mcrA Δ(mrr-hsdRMS-mcrBC) Φ80lacZΔM15 ΔlacX74 recA1 araD139 Δ(araleu)7697 galU galK rpsL (StrR) endA1 nupG)* |
| Origin | *Pseudomonas aeruginosa PAO1* |
| Parts | LasR; Gene ID: 881789 codon harmonized |
| | $P_{LasI}$; Gene ID: 881777, promoter upstream of *lasI* |
| Architecture | Synthetic |
| Experiment | 150 µL EZ Rich medium, 96-well black microtiter plate<br>PerkinElmer Ensight, exc. 588nm, emm. 633nm at 30°C<br>Fluorescence after ...h of growth (early stationary phase) |
| Notes | - |

#### LasR:3OC12-HSL

| | |
|---|---|
| *lasR* | GATGATGGCGAGCATTCACCTGCGGTAGCTA**TTACAGGGTAATCAGACCCAGATTAACT**<br>PaqCI recognition sequence PaqCI restriction site<br>**GCCATAATTGCTGCAACACGACGGCTGGTAACACCAAATTTGCGACGAATATTACCC**<br>**ATATGAAAGTTCACGTTGGCTTCGCTACAATTACAAATAACGCTAATTTCCCAGCTG**<br>**GTTTTACCAATTGCACACCACTGCAGAACTTCTTTTTCACGGCTGGTCAGAACCACC**<br>**GGTTTGCTAACCGGATGTTCAAATGCCAGACCGGCACCGCTCTGCAGTGCATAATCT**<br>**TTCAGCATCCACAGTGTCGGCAGCACGCTTTCCATAAAACGATTTGCTTCGGCACGA**<br>**TTTTCTGCTTCAACGCTCAGGCTCAGTGCACCCAGTTCACCACGTGCGCCATGCAGC**<br>*lasR*<br>**GGCATGGTCAGACCATAAACCAGACCTGCTGCGCTTGCTTCTTCAAAAAATTCGTGC**<br>**TGTTTACGGGTCTGATAAATGCTCGGTTCCCAAAAAATCGGCAGAACGCTCTGGGTA**<br>**CAATGGCTAACGGTCGGATCAACACGTGCATAACCTGCACGATCATAATGTTCACGC**<br>**CATGCTGCCGGATAATTACCCACGATAAAGGCATTTTCATAATCCTGGCTATCTTTC**<br>**GGCAGCAGACCAAACAGAATTTTGCTAAAACCCAGATCGCTTGCCATTTTCTGCAGG**<br>**ATTGCTGACCATTCCAGTTTACCGCTGCTACGTTCCAGTTCCAGAAAACCATCAACC**<br>←<br>**AGTGCCAT**AGAGACGCTCCTTCAGTTCCCCGCCAGCTAGCGGGTAGTGTGACGTGGCGCA<br>BsaI restriction site BsaI recognition sequence Golden RBS PaqCI restriction site<br>GGTGATGGACTTCATGCTGAC<br>PaqCI recognition sequence |
| $P_{lasI}$ | GATGATGGCGAGCATTCACCTGCGGTACTTA**TTCGAGCCTAGCAAGGGTCCGGGTTCAC**<br>PaqCI recognition sequence PaqCI restriction site<br>**CGAAATCTATCTCATTTGCTAGTTATAAAATTATGAAATTTGCATAAATTCTTCA**TCT<br>$P_{LasI}$<br>AGAGATTAAAGAGGAGAAATACTAGATGGTGGCGCAGGTGATGGACTTCATGCTGAC<br>Bba_B0030 PaqCI restriction site PaqCI recognition sequence |

#### EsaR:3OC6-HSL

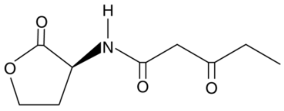


Ligand: 3-oxo-hexanoyl-L-homoserine lactone
Source: Sigma-Aldrich K3007
CAS ID: 143537-62-6
Stock: 1 mM
Solvent: DMSO
Storage: -80°C (2 months)

##### Response curve: SynEsaR(low)

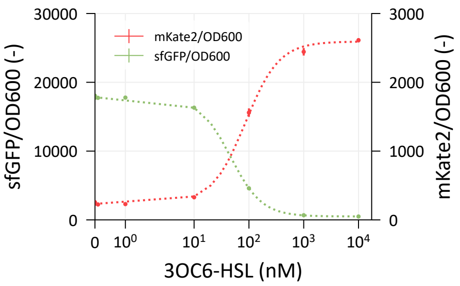


Error bars depict standard errors over three biological replicates (n=3)
a = the basal normalized fluorescent signal (leaky expression, a.u.)
k = the maximum normalized fluorescent signal relative to a (a.u.)
M = a + k = the maximum normalized fluorescent signal (a.u.)
n = the Hill coefficient (cooperativity, sigmoid character)
K = the Hill constant (TF-ligand affinity, nM)
C = the ligand concentration in the growth medium (nM)

**Hill function**

$$\left(\frac{\overline{\text{Fluo}}}{\text{OD}_{600}}\right)_{cor} = f\,(C) = a + k\left(\frac{C^n}{C^n + K^n}\right)$$

**Parameters Hill function $P_{esaR}$**

a (a.u) = $2.32*10^2$ ± $0.12*10^2$
k (a.u) = $2.36*10^3$ ± $0.03*10^3$
M (a.u) = $2.59*10^3$ ± $0.03*10^3$
n = 1.41 ± 0.13
K (nM) = 8.61*10 ± 0.64*10

**Parameters Hill function $P_{esaS}$**

a (a.u) = $4.96*10^2$ ± $0.23*10^2$
k (a.u) = $1.73*10^4$ ± $0.01*10^4$
M (a.u) = $1.78*10^4$ ± $0.01*10^4$
n = 1.52 ± 0.02
K (nM) = 4.63*10 ± 0.07*10

##### Meta-data

| | |
|---|---|
| Host | *Escherichia coli TOP10 (F- mcrA Δ(mrr-hsdRMS-mcrBC) Φ80lacZΔM15 ΔlacX74 recA1 araD139 Δ(araleu)7697 galU galK rpsL (StrR) endA1 nupG)* |
| Origin | *Pantoea stewartii* |
| Parts | EsaR: GenBank L32184.1 partly codon-optimized |
| | $P_{esaR}$: GenBank L32184.1 |
| Architecture | Synthetic |
| Experiment | 150 µL EZ Rich medium, 96-well black microtiter plate<br>PerkinElmer Ensight, exc. 588nm, emm. 633nm at 30°C<br>Fluorescence after ...h of growth (early stationary phase) |
| Notes | EsaR works as a repressor of $P_{esaR}$ in the absence of the ligand. The promoter region is bidirectional and EsaR activates $P_{esaS}$ in the absence of the ligand. |

#### EsaR:3OC6-HSL

| | |
|---|---|
| *esaR* | GATGATGGCGAGCATTCACCTGCGGTAGCTACTACCTTGCAGCTGATGCTGCCGGTCTG<br>PaqCI recognition sequence PaqCI restriction site<br>ATAAGATCCAGTTCTACACCCAGTCTGATAGCCTGTCGGGCGTTACTGACGCCCAGT<br>TTCACGACCACATTCTTGATGTGAAACTTCACGGTACTCACAGAAATGCCCGTAATA<br>GCGGCAATCTCAGCATAGGTTTTGCCCATACTCGCCCAGTACAACACCTCATTTTCA<br>CGCGAGGAAAATATCGTTTTGTCCGCGCTCTGATTTAACGCCGGGGCTCGCTCGCCT<br>TCGGTACCGGCCAGGCGGTACATCTGCTCGTTAAAATCAATCAGCAGCATCTGCATC<br>GTGCCCTGTTCGGCAGCAAGGCGTTGCTCCAGCGCAGTCTGATCGTTGCCTTTAATG<br>*esaR*<br>ATCACGGACAACAGAGCAAGGTTGTTCATGTGGTCATGCAGGACATAGGTAAAGCC<br>GTTAACGATGTTGTATTGCTTGGATAAAGAGAAAATTTTGGTGAACCGCAGGTCGG<br>ACATCAGCGTAATATTCTCATCCCAGGCAAACGGCGAGGTGCGTTTAAAGGCCGTG<br>AGAATAACCGGATCGGTCAGCTGAAAGTTGTTAGCGCGGTATAACCTAATCCATTC<br>GTCAGGATAACTGGAAATAATCAGAACATTTGAAGGATTTTTTTTGCTCACAACAGT<br>GTAAGCGTAATCCGGACTACCCAGCGGAGATAACTTTCTCTGTATGTAAGTCTGAAG<br>CGTATCCGTAATGGTCTGGTTTTCCAGGAAGAACGAGAACATAGAGACGCGGTCTATG<br>BsmBI restriction site BsmBI recognition sequence<br>ACCCGCTTCAGTTAGCGGGTAGTGTGACGTGGCGCAGGTGATGGACTTCATGCTGAC<br>Golden RBS PaqCI restriction site PaqCI recognition sequence |

#### EsaR:3OC6-HSL

| | |
|---|---|
| $P_{esaR}$ | GATGATGGCGAGCATTCACCTGCGGTACTTATTACTTATAGAGTTCATCCATGCCATGAGTAAT<br>PaqCI recognition sequence PaqCI restriction site<br>CCCCGCTGCCGTGACGAATTCCAACAGGACCATATGGTCACGCTTCTCGTTTGGGTCTTTAC<br>TAAGAACGCTTTGTGTAGACAGGTAATGATTATCCGGGAGCAGAACGGGGCCATCGCCAATC<br>GGAGTATTCTGCTGGTAATGATCAGCCAGCTGCACGGAACCGTCCTCTACGTTATGACGGAT<br>TTTAAAGTTGGCTTTGATGCCATTTTTCTGTTTATCCGCTGTAATGTATACGTTGTGCGAATTAA<br>AGTTGTATTCCAATTTGTGCCCCAGGATATTCCCATCCTCTTTGAAATCGATACCTTTTAATTCA<br>ATGCGGTTAACTAAGGTATCGCCTTCAAATTTCACTTCCGCGCGGGTCTTATACGTCCCATCG<br>*sfGFP*<br>TCTTTGAAGCTAATAGTCCGTTCCTGCACATAACCTTCAGGCATTGCGCTTTTAAAAAAGTCGT<br>GGCGTTTCATGTGATCTGGATAGCGTGAAAAGCATTGGACGCCATACGTCAACGTTGTGACC<br>AGCGTCGGCCAAGGCACGGGCAGCTTACCGGTGGTACAGATGAACTTCAGGGTCAGCTTAC<br>CATTGGTGGCATCTCCCTCTCCTTCGCCACGAACAGAGAACTTATGACCGTTCACGTCTCCG<br>TCCAGTTCTACTAAAATCGGCACAACACCGGTAAAAAGCTCTTCGCCCTTGCTCATCTAGTAC<br>TTTCCTGTGTGACTCTAGTAT**CCGCTAAACAACTGAAGCCATTGTAACCTCTGAATGAT**<br>Bba_B0032<br>**TCATTGTAAGTTACTCTTAAGTATCATCTTGCCTGTACTATAGTGCAGGTTAAGTCCA**<br>$P_{esaR/esaS}$<br>**CGTTAAGTAAAAGAAGCAGC**TCTAGAGATTAAAGAGGAGAAATACTAGATGGTGGCGCA<br>Bba_B0030 PaqCI restriction site<br>GGTGATGGACTTCATGCTGAC<br>PaqCI recognition sequence |

#### EsaR:3OC6-HSL

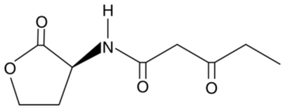


Ligand: 3-oxo-hexanoyl-L-homoserine lactone
Source: Sigma-Aldrich K3007
CAS ID: 143537-62-6
Stock: 1 mM
Solvent: DMSO
Storage: -80°C (2 months)

##### Response curve: SynEsaR(medium)

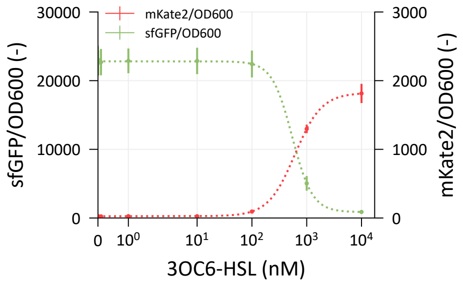


Error bars depict standard errors over three biological replicates (n=3)
a = the basal normalized fluorescent signal (leaky expression, a.u.)
k = the maximum normalized fluorescent signal relative to a (a.u.)
M = a + k = the maximum normalized fluorescent signal (a.u.)
n = the Hill coefficient (cooperativity, sigmoid character)
K = the Hill constant (TF-ligand affinity, nM)
C = the ligand concentration in the growth medium (nM)

**Hill function**

$$\left(\frac{\overline{\text{Fluo}}}{\text{OD}_{600}}\right)_{cor} = f\,(C) = a + k\left(\frac{C^n}{C^n + K^n}\right)$$

**Parameters Hill function $P_{esaR}$**

a (a.u) = $2.52*10 \pm 0.05*10$
k (a.u) = $1.80*10^3 \pm 0.05*10^3$
M (a.u) = $1.83*10^3 \pm 0.03*10^3$
n = $1.79 \pm 0.04$
K (nM) = $6.09*10^2 \pm 0.33*10^2$

**Parameters Hill function $P_{esaS}$**

a (a.u) = $8.60*10^2 \pm 0.21*10^2$
k (a.u) = $2.20*10^4 \pm 0.09*10^4$
M (a.u) = $2.29*10^4 \pm 0.09*10^4$
n = $2.37 \pm 0.25$
K (nM) = $5.43*10^2 \pm 0.35*10^2$

##### Meta-data

| | |
|---|---|
| Host | *Escherichia coli TOP10 (F- mcrA Δ(mrr-hsdRMS-mcrBC) Φ80lacZΔM15 ΔlacX74 recA1 araD139 Δ(araleu)7697 galU galK rpsL (StrR) endA1 nupG)* |
| Origin | *Pantoea stewartii* |
| Parts | EsaR: GenBank L32184.1 partly codon-optimized |
| | $P_{esaR}$: GenBank L32184.1 |
| Architecture | Synthetic |
| Experiment | 150 µL EZ Rich medium, 96-well black microtiter plate<br>PerkinElmer Ensight, exc. 588nm, emm. 633nm at 30°C<br>Fluorescence after ...h of growth (early stationary phase) |
| Notes | EsaR works as a repressor of $P_{esaR}$ in the absence of the ligand. The promoter region is bidirectional and EsaR activates $P_{esaS}$ in the absence of the ligand. |

#### EsaR:3OC6-HSL

| | |
|---|---|
| *esaR* | GATGATGGCGAGCATTCACCTGCGGTAGCTA**CTACCTTGCAGCTGATGCTGCCGGTCTG**<br>PaqCI recognition sequence PaqCI restriction site<br>**ATAAGATCCAGTTCTACACCCAGTCTGATAGCCTGTCGGGCGTTACTGACGCCCAGT**<br>**TTCACGACCACATTCTTGATGTGAAACTTCACGGTACTCACAGAAATGCCCGTAATA**<br>**GCGGCAATCTCAGCATAGGTTTTGCCCATACTCGCCCAGTACAACACCTCATTTTCA**<br>**CGCGAGGAAAATATCGTTTTGTCCGCGCTCTGATTTAACGCCGGGGCTCGCTCGCCT**<br>**TCGGTACCGGCCAGGCGGTACATCTGCTCGTTAAAATCAATCAGCAGCATCTGCATC**<br>**GTGCCCTGTTCGGCAGCAAGGCGTTGCTCCAGCGCAGTCTGATCGTTGCCTTTAATG**<br>*esaR*<br>**ATCACGGACAACAGAGCAAGGTTGTTCATGTGGTCATGCAGGACATAGGTAAAGCC**<br>**GTTAACGATGTTGTATTGCTTGGATAAAGAGAAAATTTTGGTGAACCGCAGGTCGG**<br>**ACATCAGCGTAATATTCTCATCCCAGGCAAACGGCGAGGTGCGTTTAAAGGCCGTG**<br>**AGAATAACCGGATCGGTCAGCTGAAAGTTGTTAGCGCGGTATAACCTAATCCATTC**<br>**GTCAGGATAACTGGAAATAATCAGAACATTTGAAGGATTTTTTTTGCTCACAACAGT**<br>**GTAAGCGTAATCCGGACTACCCAGCGGAGATAACTTTCTCTGTATGTAAGTCTGAAG**<br>**CGTATCCGTAATGGTCTGGTTTTCCAGGAAGAACGAGAACAT**AGAGACGTATCTTAAA<br>BsmBI restriction site BsmBI recognition sequence<br>TTTGTGGGGGGCTAGCGGGTAGTGTGACGTGGCGCAGGTGATGGACTTCATGCTGAC<br>Golden RBS PaqCI restriction site PaqCI recognition sequence |

#### EsaR:3OC6-HSL

| | |
|---|---|
| $P_{esaR}$ | GATGATGGCGAGCATTCACCTGCGGTACTTATTACTTATAGAGTTCATCCATGCCATGAGTAAT<br>PaqCI recognition sequence PaqCI restriction site<br>CCCCGCTGCCGTGACGAATTCCAACAGGACCATATGGTCACGCTTCTCGTTTGGGTCTTTAC<br>TAAGAACGCTTTGTGTAGACAGGTAATGATTATCCGGGAGCAGAACGGGGCCATCGCCAATC<br>GGAGTATTCTGCTGGTAATGATCAGCCAGCTGCACGGAACCGTCCTCTACGTTATGACGGAT<br>TTTAAAGTTGGCTTTGATGCCATTTTTCTGTTTATCCGCTGTAATGTATACGTTGTGCGAATTAA<br>AGTTGTATTCCAATTTGTGCCCCAGGATATTCCCATCCTCTTTGAAATCGATACCTTTTAATTCA<br>ATGCGGTTAACTAAGGTATCGCCTTCAAATTTCACTTCCGCGCGGGTCTTATACGTCCCATCG<br>*sfGFP*<br>TCTTTGAAGCTAATAGTCCGTTCCTGCACATAACCTTCAGGCATTGCGCTTTTAAAAAAGTCGT<br>GGCGTTTCATGTGATCTGGATAGCGTGAAAAGCATTGGACGCCATACGTCAACGTTGTGACC<br>AGCGTCGGCCAAGGCACGGGCAGCTTACCGGTGGTACAGATGAACTTCAGGGTCAGCTTAC<br>CATTGGTGGCATCTCCCTCTCCTTCGCCACGAACAGAGAACTTATGACCGTTCACGTCTCCG<br>TCCAGTTCTACTAAAATCGGCACAACACCGGTAAAAAGCTCTTCGCCCTTGCTCATCTAGTAC<br>TTTCCTGTGTGACTCTAGTAT**CCGCTAAACAACTGAAGCCATTGTAACCTCTGAATGAT**<br>Bba_B0032<br>**TCATTGTAAGTTACTCTTAAGTATCATCTTGCCTGTACTATAGTGCAGGTTAAGTCCA**<br>$P_{esaR/esaS}$<br>**CGTTAAGTAAAAGAAGCAGC**TCTAGAGATTAAAGAGGAGAAATACTAGATGGTGGCGCA<br>Bba_B0030 PaqCI restriction site<br>GGTGATGGACTTCATGCTGAC<br>PaqCI recognition sequence |

#### EsaR:3OC6-HSL

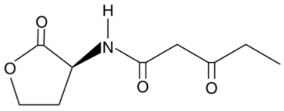

Ligand: 3-oxo-hexanoyl-L-homoserine lactone
Source: Sigma-Aldrich K3007
CAS ID: 143537-62-6
Stock: 1 mM
Solvent: DMSO
Storage: -80°C (2 months)

##### Response curve: SynEsaR(high)

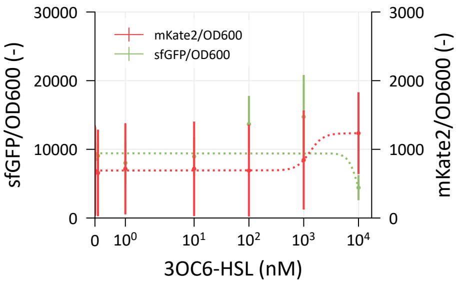


Error bars depict standard errors over three biological replicates (n=3)
a = the basal normalized fluorescent signal (leaky expression, a.u.)
k = the maximum normalized fluorescent signal relative to a (a.u.)
M = a + k = the maximum normalized fluorescent signal (a.u.)
n = the Hill coefficient (cooperativity, sigmoid character)
K = the Hill constant (TF-ligand affinity, nM)
C = the ligand concentration in the growth medium (nM)

**Hill function**

$$\left(\frac{\overline{\text{Fluo}}}{\text{OD}_{600}}\right)_{cor} = f\,(C) = a + k\left(\frac{C^n}{C^n + K^n}\right)$$

**Parameters Hill function $P_{esaR}$**

Due to the high biological variability, the Hill function could not be fitted.

**Parameters Hill function $P_{esaS}$**

Due to the high biological variability, the Hill function could not be fitted.

##### Meta-data

| | |
|---|---|
| Host | *Escherichia coli TOP10 (F- mcrA Δ(mrr-hsdRMS-mcrBC) Φ80lacZΔM15 ΔlacX74 recA1 araD139 Δ(araleu)7697 galU galK rpsL (StrR) endA1 nupG)* |
| Origin | *Pantoea stewartii* |
| Parts | EsaR: GenBank L32184.1 partly codon-optimized |
| | $P_{esaR}$: GenBank L32184.1 |
| Architecture | Synthetic |
| Experiment | 150 µL EZ Rich medium, 96-well black microtiter plate<br>PerkinElmer Ensight, exc. 588nm, emm. 633nm at 30°C<br>Fluorescence after ...h of growth (early stationary phase) |
| Notes | EsaR works as a repressor of $P_{esaR}$ in the absence of the ligand. The promoter region is bidirectional and EsaR activates $P_{esaS}$ in the absence of the ligand. |

#### EsaR:3OC6-HSL

| | |
|---|---|
| *esaR* | GATGATGGCGAGCATTCACCTGCGGTAGCTA**CTACCTTGCAGCTGATGCTGCCGGTCTG**<br>PaqCI recognition sequence PaqCI restriction site<br>**ATAAGATCCAGTTCTACACCCAGTCTGATAGCCTGTCGGGCGTTACTGACGCCCAGT**<br>**TTCACGACCACATTCTTGATGTGAAACTTCACGGTACTCACAGAAATGCCCGTAATA**<br>**GCGGCAATCTCAGCATAGGTTTTGCCCATACTCGCCCAGTACAACACCTCATTTTCA**<br>**CGCGAGGAAAATATCGTTTTGTCCGCGCTCTGATTTAACGCCGGGGCTCGCTCGCCT**<br>**TCGGTACCGGCCAGGCGGTACATCTGCTCGTTAAAATCAATCAGCAGCATCTGCATC**<br>**GTGCCCTGTTCGGCAGCAAGGCGTTGCTCCAGCGCAGTCTGATCGTTGCCTTTAATG**<br>*esaR*<br>**ATCACGGACAACAGAGCAAGGTTGTTCATGTGGTCATGCAGGACATAGGTAAAGCC**<br>**GTTAACGATGTTGTATTGCTTGGATAAAGAGAAAATTTTGGTGAACCGCAGGTCGG**<br>**ACATCAGCGTAATATTCTCATCCCAGGCAAACGGCGAGGTGCGTTTAAAGGCCGTG**<br>**AGAATAACCGGATCGGTCAGCTGAAAGTTGTTAGCGCGGTATAACCTAATCCATTC**<br>**GTCAGGATAACTGGAAATAATCAGAACATTTGAAGGATTTTTTTTGCTCACAACAGT**<br>**GTAAGCGTAATCCGGACTACCCAGCGGAGATAACTTTCTCTGTATGTAAGTCTGAAG**<br>**CGTATCCGTAATGGTCTGGTTTTCCAGGAAGAACGAGAACAT**AGAGACGCCTCCTCGT<br>BsmBI restriction site BsmBI recognition sequence<br>AGATGTTAGTAGCAGCGGGTAGTGTGACGTGGCGCAGGTGATGGACTTCATGCTGAC<br>Golden RBS PaqCI restriction site PaqCI recognition sequence |

#### EsaR:3OC6-HSL

| | |
|---|---|
| $P_{esaR}$ | GATGATGGCGAGCATTCACCTGCGGTACTTATTACTTATAGAGTTCATCCATGCCATGAGTAAT<br>PaqCI recognition sequence PaqCI restriction site<br>CCCCGCTGCCGTGACGAATTCCAACAGGACCATATGGTCACGCTTCTCGTTTGGGTCTTTAC<br>TAAGAACGCTTTGTGTAGACAGGTAATGATTATCCGGGAGCAGAACGGGGCCATCGCCAATC<br>GGAGTATTCTGCTGGTAATGATCAGCCAGCTGCACGGAACCGTCCTCTACGTTATGACGGAT<br>TTTAAAGTTGGCTTTGATGCCATTTTTCTGTTTATCCGCTGTAATGTATACGTTGTGCGAATTAA<br>AGTTGTATTCCAATTTGTGCCCCAGGATATTCCCATCCTCTTTGAAATCGATACCTTTTAATTCA<br>ATGCGGTTAACTAAGGTATCGCCTTCAAATTTCACTTCCGCGCGGGTCTTATACGTCCCATCG<br>*sfGFP*<br>TCTTTGAAGCTAATAGTCCGTTCCTGCACATAACCTTCAGGCATTGCGCTTTTAAAAAAGTCGT<br>GGCGTTTCATGTGATCTGGATAGCGTGAAAAGCATTGGACGCCATACGTCAACGTTGTGACC<br>AGCGTCGGCCAAGGCACGGGCAGCTTACCGGTGGTACAGATGAACTTCAGGGTCAGCTTAC<br>CATTGGTGGCATCTCCCTCTCCTTCGCCACGAACAGAGAACTTATGACCGTTCACGTCTCCG<br>TCCAGTTCTACTAAAATCGGCACAACACCGGTAAAAAGCTCTTCGCCCTTGCTCATCTAGTAC<br>TTTCCTGTGTGACTCTAGTAT**CCGCTAAACAACTGAAGCCATTGTAACCTCTGAATGAT**<br>Bba_B0032<br>**TCATTGTAAGTTACTCTTAAGTATCATCTTGCCTGTACTATAGTGCAGGTTAAGTCCA**<br>$P_{esaR/esaS}$<br>**CGTTAAGTAAAAGAAGCAGC**TCTAGAGATTAAAGAGGAGAAATACTAGATGGTGGCGCA<br>Bba_B0030 PaqCI restriction site<br>GGTGATGGACTTCATGCTGAC<br>PaqCI recognition sequence |
